# Neural drift during rest drives walking direction and memory consolidation in Drosophila

**DOI:** 10.1101/2025.03.20.644317

**Authors:** Andres Flores-Valle, Rolf Honnef, Johannes D. Seelig

## Abstract

Heading is represented in head direction systems across species, including *Drosophila*. However, how navigation decisions are made and how goal memories are maintained, particularly during rest, is little understood. Here, using a navigation learning assay for flies walking in virtual reality, we describe neural dynamics for direction selection and memory. Activity of a subset of neurons in the fan-shaped body, a navigation and memory-related structure, continually drifts when the animal is at rest, centered 180° opposite to the walking direction. Optogenetic activation during rest biases subsequent walking direction. Learning shifts rest activity, revealing a memory of goal direction. Downstream neurons correct the 180° shift, reactivating walking directions during rest; the connectome reveals a compact circuit consistent with reactivation, validated by computational modeling. Thus, rest activity drives behavior, reactivates walking directions, and is shaped by memory, suggesting shared principles for memory consolidation in navigation circuits across species.

---

Animal movement is shaped by memory across species^1^. Many animals, including a variety of insects^2^, navigate over long distances relying on memories of visual cues, and the features attributed to these memories are key for decision making^1–4^. Navigation towards visual cues requires selecting and maintaining a goal in memory during behavior^2,5,6^. The head direction system represents an animal’s orientation with respect to these visual cues and goals^7–11^. Goal related navigation circuits have been described in bats^12^, rats^5,6^, and insects, including *Drosophila*^8,13–17^. While mammalian place and grid cells change their spatial arrangement during goal learning^5,6,8,18^, less is known about head direction cells. In mammalian circuits, decision making and learning are additionally accompanied by navigation-related activity during rest and sleep, such as replay^19–24^.

In *Drosophila*, memories for visual navigation behavior have been located in the central complex (CX)^25–29^, a brain area at the center of sensorimotor integration. The CX contains the fly’s head direction system with circuits that share features of ring attractor networks^9,10,30–34^. Head direction circuits, including neurons that determine goal direction^14^, have been investigated using two-photon imaging in flies during learning behavior^35,36^.

Behavior, physiology, and computational modeling point to the importance of the fan-shaped body, a substructure of the CX, for the encoding of memory, including consolidation through reactivation during sleep^37^, and propose mechanisms for direction selection^6,8,10,16,17,38^. However, where behavioral decisions, such as a choice of navigation direction, occur in the network, how goals are selected, and how memories are consolidated remains poorly understood^6,39,40^.

Here we describe a mechanism for memory consolidation and direction selection in the *Drosophila* fan-shaped body. Using two-photon imaging and optogenetics, we show that PFR neurons track heading direction during walking but exhibit persistent drift across columns during rest. The distribution of drift during rest is shifted by 180°, that is, opposite to the walking direction. Optogenetic stimulation during rest determines subsequent walking direction. Learning biases rest drift towards the opposite direction of the learned goal direction, revealing the emergence of goal memory. We further identify hΔ neurons as downstream PFR targets that also drift during rest but lack the 180° shift between walking and rest, and a connectome-based model proposes a mechanism for how PFR-hΔ connectivity could account for these dynamics. These experiments reveal neural dynamics underlying direction selection, learning, and memory consolidation during rest and navigation.

## PFR neurons show drift during rest

For imaging neural activity in navigating flies, we combined a virtual mechanical arena with a diameter of 0.5 meters (Fig. 1a and Extended Data Fig. 1a) in which flies can perform spatial learning^41^ with two-photon calcium imaging^42,43^. In this setup, tethered flies navigate on an air-supported ball and the readout of ball motion serves to rotate and translate the arena in two dimensions (Fig. 1b, see Methods)^41^. The relative positions and orientations of the fly with respect to the arena thus remain similar to those of an animal walking freely in a stationary arena. In these experiments, the arena featured a single bright vertical stripe on an otherwise dark background (Figs. 1a and b). We validated the setup and closed-loop imaging with GCaMP8m in EPG heading neurons^30^, which reliably tracked the fly’s orientation in the arena (Extended Data Fig. 2 and Supplementary Figs. S1–S3).

**Figure 1.**
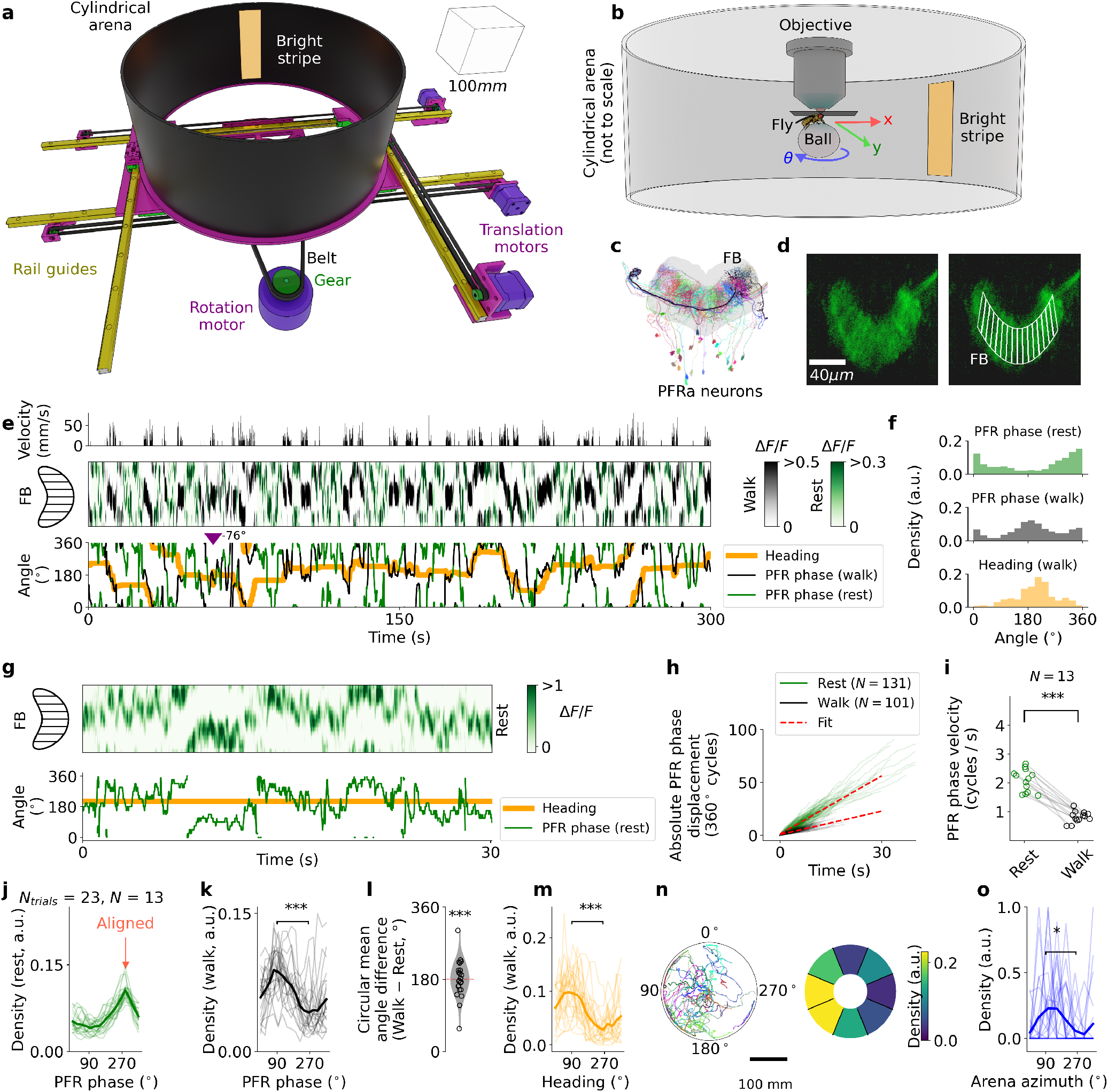
Virtual reality setup and PFR drift. **a** Mechanical virtual reality (colors for visualization). Cube of 100 mm length is shown for scale. A smaller version of the mechanical arena as well as place learning behavior without imaging are described in^41^ and in Methods. **b** A fly walks on an air-suspended ball inside the moving cylindrical arena, enabling 2D navigation (translation in *x*, *y* and rotation around *θ*). The arena, shown here with a bright stripe on a black background, moves while the fly remains stationary during two-photon imaging. **c** PFRa neurons, a subpopulation of PFR neurons (see Supplementary Information), projecting to the FB, each in a different color. Image downloaded from^77,78^ **d** Expression of calcium indicator in PFR neurons with SS02270 (average of 500 frames). Right: ROIs across the FB. **e** Example of PFR calcium activity during navigation. Top: fly velocity. Middle: PFR activity across 16 ROIs (panel d) during rest (green) and walking (black). Bottom: PFR phase during rest (green) and walking (black), and fly heading (orange). Inverted purple triangle indicates switching offset correction (see Methods). **f** Distributions of PFR phase during rest (top), during walking (middle) and heading (bottom) across 16 bins for the fly in e. **g** Example of PFR calcium activity during 30 seconds of rest. Top: PFR activity across 16 ROIs. Bottom: PFR phase during rest (green) and heading (orange). **h** Absolute PFR phase displacement from 13 flies during *≥* 5 seconds of continuous rest (green) and walking (black) epochs, with red lines representing linear fit (see Methods). **i** PFR phase velocity during rest (green) and walking (black) in cycles per second (cycles across the 16 ROIs) for 13 flies. **j** PFR phase density during rest for all flies, with maximum aligned to 270°. **k** PFR phase density during walking after alignment (see Methods). **l** Difference in circular mean PFR phase between walking and resting conditions. Points represent single flies. **m** Fly heading distribution across 16 bins after alignment. **n** Left: aligned fly trajectories. Right: trajectory density across 8 azimuthal bins at radial distance *≥* 50 mm. **o** Aligned arena azimuth density (8 bins, radial distance *≥* 50 mm) for all flies (see angles in panel n, left). Thick lines in panels j, k, m and o represent mean across flies. Three asterisks in panels i and l show statistical significance based on paired two-sided t-test across flies (p < 0.00005). Asterisks in density panels (k, m and o) indicate significant differences between 90° and 270° (Kolmogorov-Smirnov test: p < 0.005 (**), p < 0.0005 (***)). Data from all flies is shown in Supplementary Figs. S7-S11

PFR neurons, which innervate the Fan-shaped Body (FB), project into different FB columns (Fig. 1c, see Methods) and receive input from hΔB neurons, which encode the traveling direction of the fly in allocentric coordinates (Supplementary Fig. S4a)^44–46^. PFR neurons therefore show similar dynamics to hΔB neurons^46^. To record activity during navigation we expressed the calcium indicator GCaMP8m^47^ in PFR neurons using the split-GAL4 line SS02270^48^ (SS02270 > UAS-GCaMP8m, Fig. 1d). In these experiments, flies could freely explore the arena (in closed loop) for 5 minutes per trial, across 3 trials, starting at the center of the arena.

PFR activity formed a moving bump or cosine-like profile across the FB during walking (see Supplementary Figs. S5d, S6, and Methods) with its phase (see Methods) tracking heading direction of the fly relative to the bright stripe as previously described^44–46^. Heading direction signals display an arbitrary offset relative to landmarks that can switch in head-fixed conditions^30,49^. Therefore, the offset between the bright stripe orientation and PFR phase was first removed (allowing for one offset switch, see Methods, Extended Data Fig. 3a-e). We occasionally observed phase jumps in PFR neurons during walking, which have been identified as reflecting changes in traveling direction, such as diagonal, sideways, or backward walking^45,46^. As a consequence, PFR heading-phase correlation was lower than in EPG neurons (Extended Data Fig. 2d). However, we did not analyze these jumps further, since flies mostly walked forward or rotated, and in this situation the phase simply follows the heading direction.

We found that when the fly is standing still, PFR activity nevertheless drifts (Figs. 1e and f, green)^50^, maintaining a cosine-like activity profile similar to that during walking (see Supplementary Fig. S5d and Methods). This differs from other head direction cells, where bump position is stable during periods of rest over tens of seconds^30^. Drift in PFR neurons has been observed anecdotally^46^, but we find here that drift persists over long bouts of immobility (Fig. 1g). Drift activity moved at about twice the speed of activity observed during walking (Fig. 1h and i, see Methods).

## PFR phase shifts by 180° between rest and walking

Comparing PFR activity during rest and walking epochs in a trial of 5 minutes of navigation (Fig. 1e) shows that the distribution of PFR phase during rest is approximately opposite to the phase distribution during walking (Fig. 1f), that is, highest in FB columns that are inactive during walking and *vice versa*. Since the PFR phase corresponds to the heading of the fly in the arena during walking, the heading distribution is also opposite to the distribution during rest (Fig. 1f, orange).

To compare activity distributions across trials and flies (see Supplementary Figs. S7-S11), we first removed the arbitrary phase offset intrinsic to heading direction signals (see Methods). We then aligned the imaging data using the phase distribution during rest, setting the peak of this distribution arbitrarily to 270° (Fig. 1j), in the following referred to as the “rest-phase offset” (see Extended Data Fig. 3f-g and Methods). After alignment, the PFR phase distribution during walking was centered around 90° across flies (Fig. 1k), revealing a consistent 180° shift relative to the mean phase during rest (Fig. 1l).

Similarly, after rotating all headings (behavior) to their corresponding rest-phase offset, we found that the heading direction of flies was shifted by 180° compared to the peak of the PFR phase distribution during rest (Fig. 1m, centered around 90°). In these experiments, flies mostly walked outwards from the center of the arena at the beginning of the trial towards the rim of the arena at the end of the trial. Therefore, after rotating each trajectory by its corresponding rest-phase offset (see Extended Data Fig. 3h-i and Methods), flies mostly walked in the direction opposite to the peak PFR phase distribution at rest (Fig. 1n and o, centered around 90°). This shows that the 180° phase shift between walking and rest also occurs during directional walking. Flies therefore not only orient themselves 180° opposite to the rest PFR phase distribution but also walk towards that direction. Heading is the orientation of the fly, but walking direction can be independent of orientation, since flies can walk backwards, diagonally, or sideways while maintaining the same heading.

Using a second GAL4 line (R27G06^51^), which also labels PFR neurons^44^ along with other neurons in the ellipsoid body (Extended Data Fig. 4a–l), we reproduced all findings, including persistent drift during rest with a cosine-like activity profile (which was again faster than during walking), a 180° phase shift between drift during rest and activity during walking, heading, and walking direction (Extended Data Fig. 4a–l, Supplementary Fig. S12). Thus, the 180° activity shift in PFR neurons between rest and walking suggests different computations across behavioral states.

## Optogenetic activation during walking shapes rest activity

We next investigated how PFR activity during rest relates to activity during walking, and whether rest activity is shaped by prior walking activity. To monitor and perturb activity at the same time, we combined two-photon imaging with optogenetics (Extended Data Fig. 5a, see Methods)^33,52^. We co-expressed the calcium indicator GCaMP8m^47^ and the optogenetic actuator CsChrimson^53^ using the GAL4 line R27G06 (R27G06-GAL4 > UAS-GCaMP8m; UAS-CsChrimson). We used this line because reliable calcium recordings could not be obtained when expressing both constructs in the split-GAL4 line used in Fig. 1. For control flies, we only expressed GCaMP8m (R27G06-GAL4 > UAS-GCaMP8m).

We used a red laser and a spatial light modulator for targeted neuronal activation (Extended Data Fig. 5a, see Methods)^52^. This approach ensured that the red laser used for optogenetics did not directly excite GCaMP8m. Therefore, fluorescence changes during activation reflected only calcium activity, not optical artifacts.

In these experiments, flies freely explored a closed-loop arena with a single bright stripe. Walking and resting periods were detected in real time. During walking, optogenetic activation of a subset of PFR columns (Extended Data Fig. 5b) increased calcium signals at the stimulated location, with activity centered around it (Extended Data Fig. 5b, second row and c). Optogenetic activation was automatically turned off when the fly stopped walking. The optogenetics laser did not significantly alter the overall balance of walking and resting behavior compared to navigation without laser (Extended Data Fig. 6a).

Across flies, activation during walking produced low correlation between PFR phase and heading, and often induced continuous turning (see Supplementary Figs. S13–S15), indicating that the heading signal was disrupted in the FB during stimulation: the activity bump remained fixed at the stimulation site rather than tracking heading (Extended Data Fig. 5d).

## Optogenetic activation during rest drives walking behavior

Neural activity during rest is important for decision making, planning, memory consolidation, and learning in mammalian navigation circuits^19–24^ and could also play a similar role in the fly. We therefore asked whether PFR activity during rest was influencing subsequent activity during walking and behavior.

We again used the GAL4 line R27G06, co-expressing GCaMP8m and CsChrimson (R27G06-GAL4 > UAS-GCaMP8m; UAS-CsChrimson); control flies expressed GCaMP8m alone. In these experiments, flies could again freely explore the arena (closed loop) with a single bright stripe, and activation of a subset of columns of PFR neurons during rest (Fig. 2a, top and second) resulted in an increase in calcium at the activated site (Fig. 2b, green). Activation ended as soon as the fly resumed walking and did not affect the rest-to-walk balance compared to navigation without laser (Extended Data Fig. 6b). For all flies, the arbitrary heading offset during walking was removed (Extended Data Fig. 3a-e), and the trajectory in the arena was correspondingly transformed by this offset (Extended Data Fig. 3h, see Methods).

**Figure 2.**
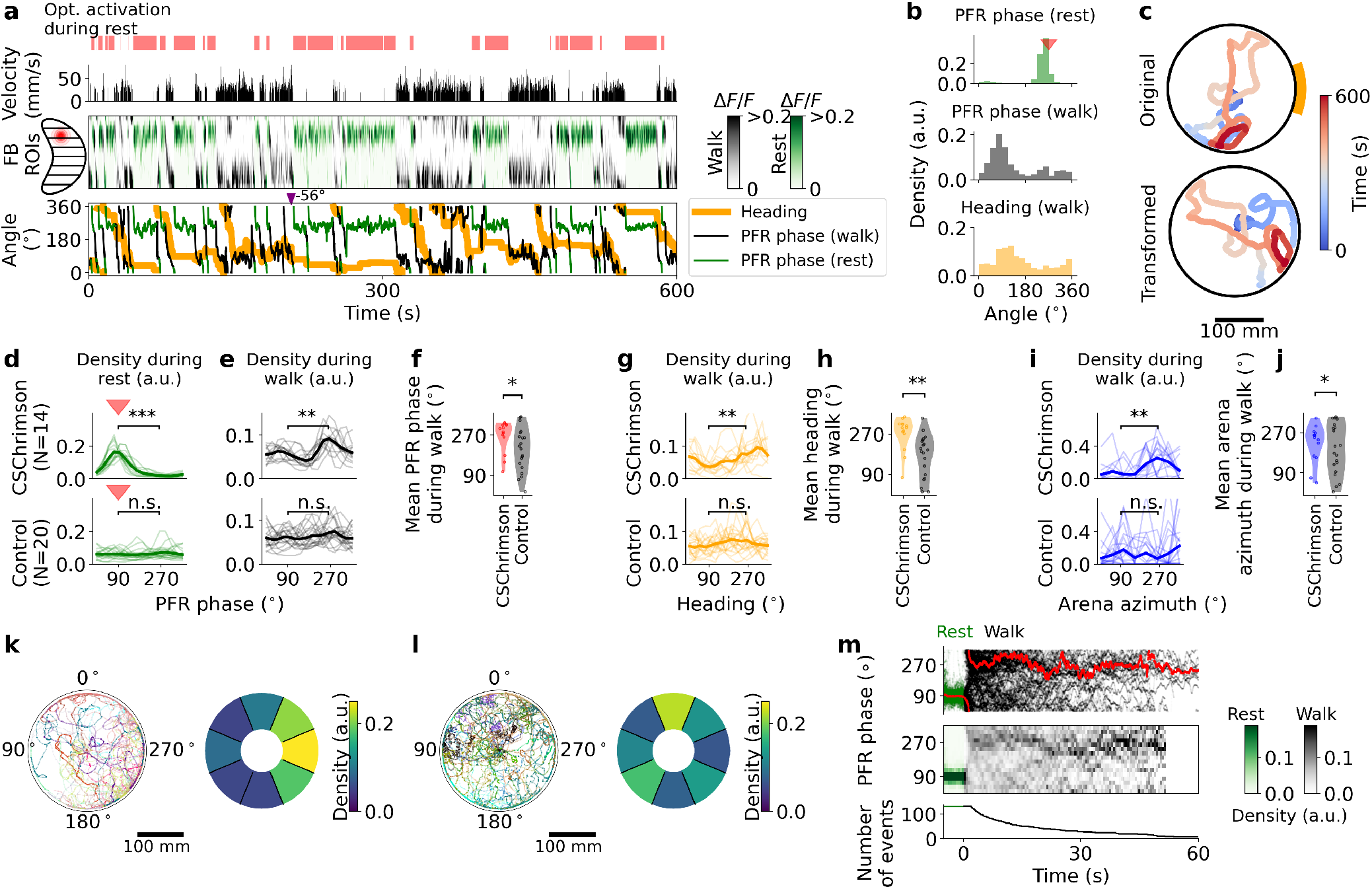
Optogenetic activation during rest drives subsequent behavior. **a** Optogenetic activation of PFR columns during rest in navigating flies. Top: laser for optogenetics is only switched on when the fly rests (walking velocity is zero). First row: walking velocity of the fly. Second row: fluorescence from 16 ROIs defined in the FB. Optogenetic activation was performed in columns on one side of the FB (shown on the left). Third row: phase of PFR neurons (see Methods), during walking (black) and rest (green). Orange: fly orientation in the mechanical virtual reality. Inverted purple triangle indicates switching offset correction (see Methods). **b** Top: PFR phase distribution during rest. Red triangle indicates corresponding optogenetic activation site. Center and bottom: PFR phase and heading distributions across 16 bins during walking, respectively. **c** Trajectory of the fly in the VR. Top: original trajectory; Bottom: transformed trajectory according to PFR phase and heading offset (see Methods). Color encodes time across trial. **d** PFR phase distribution during rest for CsChrimson flies (top) and controls (bottom). **e** Same as **d** during walking. **f** Mean PFR phase during walking: CsChrimson (red) vs. controls (black). Each dot represents a single fly. **g** Heading distributions (16 bins) during walking: CsChrimson (top) vs. controls (bottom). **h** Mean heading during walking: CsChrimson (orange) vs. controls (black). **i** Transformed arena azimuth distributions (8 bins, radial distance *≥* 50 mm): CsChrimson (top) vs. controls (bottom). **j** Mean transformed arena azimuth direction: CsChrimson (blue) vs. controls (black). **k** CsChrimson: single-fly transformed trajectories (left); azimuth density (8 bins, radial distance *≥* 50 mm, right). **l** Same as **k** for controls. **m** Rest (green) to walking (black) transitions and PFR phase: PFR phase trajectories (top, red line shows the circular mean), density from top panel (middle), and transition events count (bottom). Transitions defined as *≥* 5 seconds rest followed by *≥* 2 seconds walking. Thick lines in panels d, e, g and i represent mean across flies. Asterisks in density panels (d, g, and i) indicate significant differences between 90° and 270° (Kolmogorov-Smirnov test: p < 0.005 (**), p < 0.0005 (***)). Asterisks in panels f, h, and j represent statistical significance between Chrimson and control flies (Two-Sample Kuiper Test: p < 0.05 (*), p < 0.005 (**)). Data are shown in Supplementary Figs. S19-S21 for all Chrimson flies, and in Supplementary Figs. S22-S25 for all control flies.

Across flies (Supplementary Figs. S19–S21), optogenetic activation during rest increased activity at the stimulation site (Fig. 2d, top), an effect not observed in controls (Fig. 2d, bottom; Supplementary Figs. S22–S25). Following activation, the PFR phase during walking was shifted in the direction opposite to the stimulated direction (Fig. 2e and f), along with heading (Fig. 2g and h) and transformed walking direction (Fig. 2i and j), whereas no such effect was observed in controls. As a result, flies occupied the opposite direction in the arena relative to the activation site (Fig. 2k). These effects were not observed in control flies (Fig. 2l). The PFR phase shift occurred immediately after the transition to walking and persisted for several seconds (Fig. 2m). Thus, optogenetic activation of PFR neurons during rest is sufficient to drive subsequent behavior in the opposite direction of the activation site.

## Goal memory emerges during learning

We next asked whether directional learning would lead to changes in PFR drift activity during rest. To investigate this, we combined an assay for tethered place learning (Fig. 3a,b)^41^, similar to place learning in freely walking flies^28^, with two-photon calcium imaging. In these experiments, flies learned to find a cool area in the mechanical virtual reality with two visual landmarks, one bright (yellow in Fig. 3a) and one dim (gray in Fig. 3a, see Methods). To simulate the cool area, the fly was heated with an IR laser (Fig. 3b) unless it entered the cool area (green area in Fig. 3a, left), where the IR laser switched off. Flies were trained for 10 trials and each trial lasted until the fly stayed inside the cool area for at least 10 s. The position of the bright and dim stripes were then swapped, the cool area staying with the same stripe, thus repositioning the fly and the goal to opposite sides of the arena. The next trial then started with the fly again navigating in closed loop to the cool area (Fig. 3a, right). If the fly did not find the cool area within 15 minutes during a trial, a new trial started (again after swapping dim and bright stripe positions as well as the cool area, see Methods).

**Figure 3.**
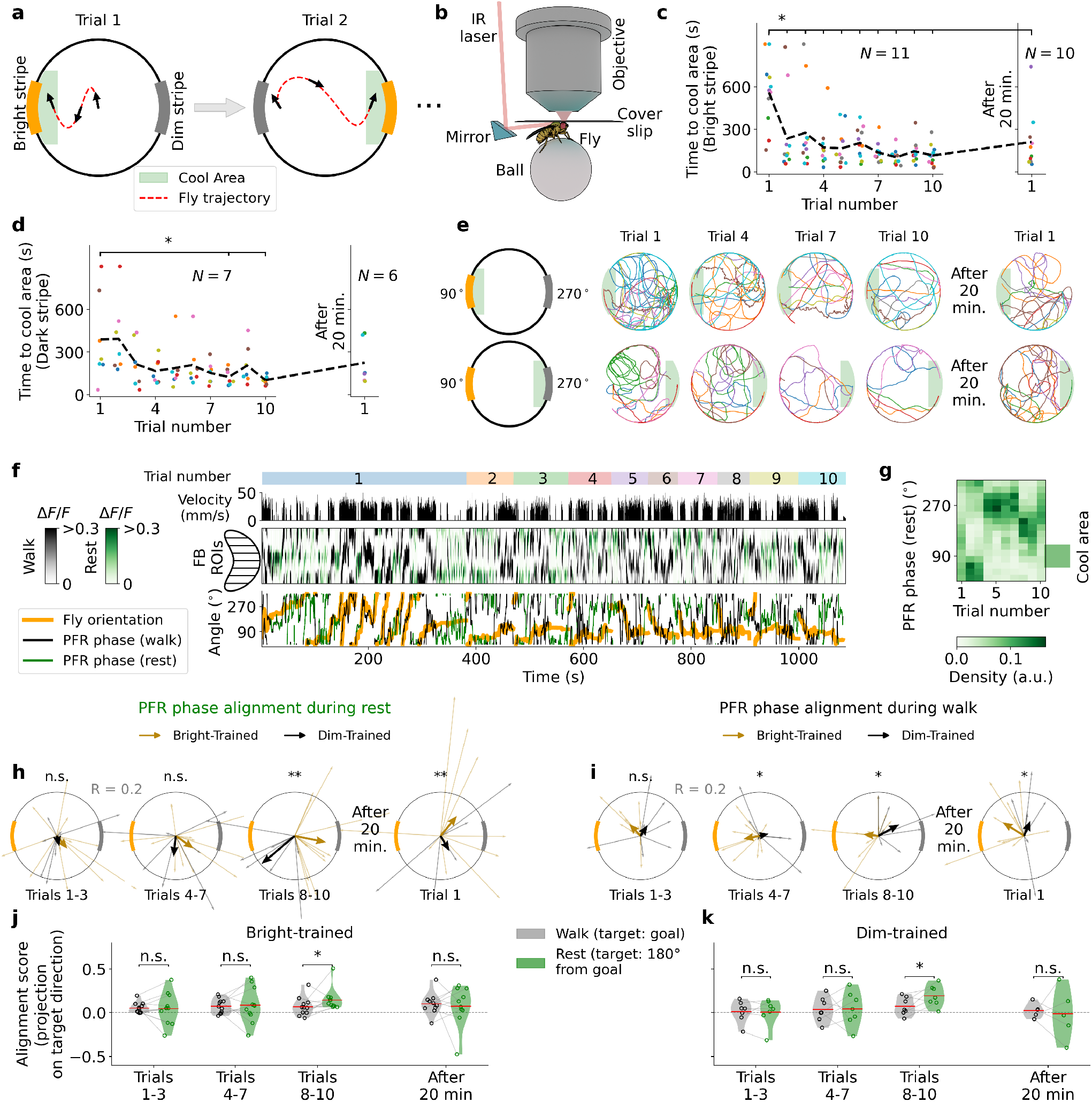
PFR activity during learning. **a** Schematic of behavior protocol for place learning experiment: flies are trained over 10 trials in a mechanical arena with two stripes (one dim, one bright, see Methods). First trial: fly starts at center of arena, has to reach cool area, and stay there for 10 seconds (red line). The two stripes switch positions, and the cool area moves to the other side. The fly has to reorient and find the cool area again. This pattern repeats for all trials, with the cool area and stripes switching upon trial completion. **b** The aversive heat stimulus is delivered with an IR laser, focused on the thorax of the fly using a mirror. The laser is switched off in the cool area. **c** Left: time taken for all flies to reach the cool area, positioned in front of the bright stripe for 10 trials. Right: time to reach the cool area after a 20-minute break. **d**: Same as c, but with the cool area in front of the dim stripe. **e** First row: fly trajectories over trials 1, 4, 7, and 10 when trained with cool area in front of the bright stripe (protocol schematic in first column). Second row: same as previous, but with the cool area in front of the dim stripe. **f** Example of place learning during imaging for a single fly. Top row: trial number. First row: walking velocity. Second row: fluorescence in the FB during walking (black) and resting (green). Third row: fly orientation (orange) and PFR phase during walking (black) and rest (green). The offset between PFR phase and arena orientation is corrected so that fluorescence and PFR phase during walking align with fly orientation (see Methods). **g** PFR phase density during rest across trials for the fly in panel f. The green rectangle indicates the direction of the cool area (90° for the bright stripe). **h** Mean resultant vector (aligning vector, see Methods) of the PFR phase distribution during rest across trials for bright-stripe (orange) and dim-stripe (gray) training. Light traces represent individual flies; thick traces represent the group mean. The vector points toward the direction where the PFR phase distribution is centered during rest. **i** Same as h, but for the PFR phase during walking. **j** PFR alignment to the stimulus-defined target direction (goal during walking, goal-opposite, i.e. 180°, during rest) across trials in bright-trained flies. Violins: distributions; circles: individual flies; red lines: means. **k** Same as j for dim-trained flies. Asterisks in panels h and i indicate statistical significance between the two groups using Hotelling’s *T* ^2^ test. Asterisks in panels j and k indicate one-sided paired Wilcoxon test (rest *>* walk). All learning data are in Supplementary Figs. S26–S29 (bright-stripe) and S30–S32 (dim-stripe).

To analyze neural activity during successful goal-directed learning, we restricted the analysis to flies that demonstrated reliable learning based on a learning index (see Methods). This behavioral selection was conducted independently of the imaging data. For these experiments, we again used the R27G06-GAL4 driver, which supports reliable learning in tethered flies navigating the mechanical VR^41^. The selected flies, which successfully learned to find the cool area relative to either bright or dim (Figs. 3c-e) stripes, showed a corresponding decrease in the time required to reach the goal, as well as decrease in the distance (Extended Data Fig. 7a and b) while walking velocity remained constant across trials (Extended Data Fig. 7c and d). Flies also showed less rest time in later trials (Extended Data Fig. 7e and f).

An example of a fly that learns to locate the cool area positioned in front of the bright stripe across 10 trials is shown in Fig. 3f. Unlike in our previous experiments where we aligned the heading to the PFR phase, we now aligned the imaging data to the fly’s orientation by shifting the FB columns by the heading-phase offset (see Methods). Learning led to a shift in the PFR phase distribution during rest across the 10 trials in the direction opposite to that of the cool area (Fig. 3g, Supplementary Figs. S26-S29 for all bright stripe-trained flies, and Supplementary Figs. S30-S32 for all dim stripe-trained flies). Specifically, across all flies, the aligned PFR phase (see Methods) during rest shifted to point away from the goal location during the final trials, for both the bright and dim stripe (Fig. 3h, see Extended Data Fig. 7g for PFR phase distributions). During walking, the PFR phase was aligned toward the goal direction (Fig. 3i; see Extended Data Fig. 7h for PFR phase distributions). However, this alignment was less pronounced than the PFR phase alignment observed during rest (Fig. 3j and k), which pointed toward the opposite direction of the goal. We attribute the broader PFR phase distribution during walking to active searching behavior, whereas the sharper distribution during rest, especially in the 8-10 trials, reflects a stable memory representation of the learned goal direction. Thus, a memory signature of the learned direction is visible in the drift distribution during rest periods.

After training, we tested if flies retained the memory of the cool area over longer timescales. After 10 trials, flies rested for 20 minutes, after which their memory for the location of the cool area was tested with an additional trial. Performance after the resting period, however, showed weak improvement compared to the very first trial (Fig. 3c, e, right side and Extended Data Fig. 7a, b, right side). PFR phase alignment was also weak during rest (Fig. 3h, right side, and Extended Data Fig. 7g, right side) and walking (Fig. 3i, right side and Extended Data Fig. 7h, right side). The transition from walking to rest shows a memory trace of the walking direction that persists for several seconds (Extended Data Fig. 5g), which, together with the learning experiments, suggests that PFR drift during rest supports short-term memory of goal direction.

Together, these experiments show that direction learning changes the distribution of the PFR phase during rest. Additionally, since rest activity drives subsequent behavior (Fig. 2), drift activity during rest may play a role in guiding fly behavior toward the memorized direction.

## FB layers detect temperature changes

The FB is thought to encode attributes for directions in tangential neurons^4,6,16,54,55^. We identified a population of tangential neurons in dorsal FB layers which respond to the change in temperature of the cool area (see Extended Data Fig. 8 and Supplementary Information). This shows that in addition to direction information, also changes in temperature are encoded in the FB network which could serve to reinforce preferred directions.

## Downstream hΔ neurons show a 180° –shifted connectivity pattern

To gain insights into the computational role of the 180° shift between walking and rest, we investigated the propagation of PFR activity through downstream circuits in the connectome. The major directly connected downstream populations are the hΔA and hΔH neurons (Supplementary Table S2), columnar neurons in the fan-shaped body, which we simply refer to as hΔ neurons. Interestingly, a single hΔ neuron can receive input from two PFRa neurons in different columns (Fig.4a).

When sorted by output column (see Methods), the connectivity reveals two input patterns: one along the main diagonal of the connectivity matrix (same column) and another along an off-diagonal shifted by four columns (180°, Fig. 4b and c for hΔA; Supplementary Fig. S4c for hΔH). PFR neurons are the main columnar population showing this dual connectivity pattern; other populations target hΔ neurons through only the main diagonal (Supplementary Figs. S33 and S34), through the off-diagonal (Supplementary Figs. S35 and S36), or through both diagonals (Supplementary Figs. S37 and S38) .

**Figure 4.**
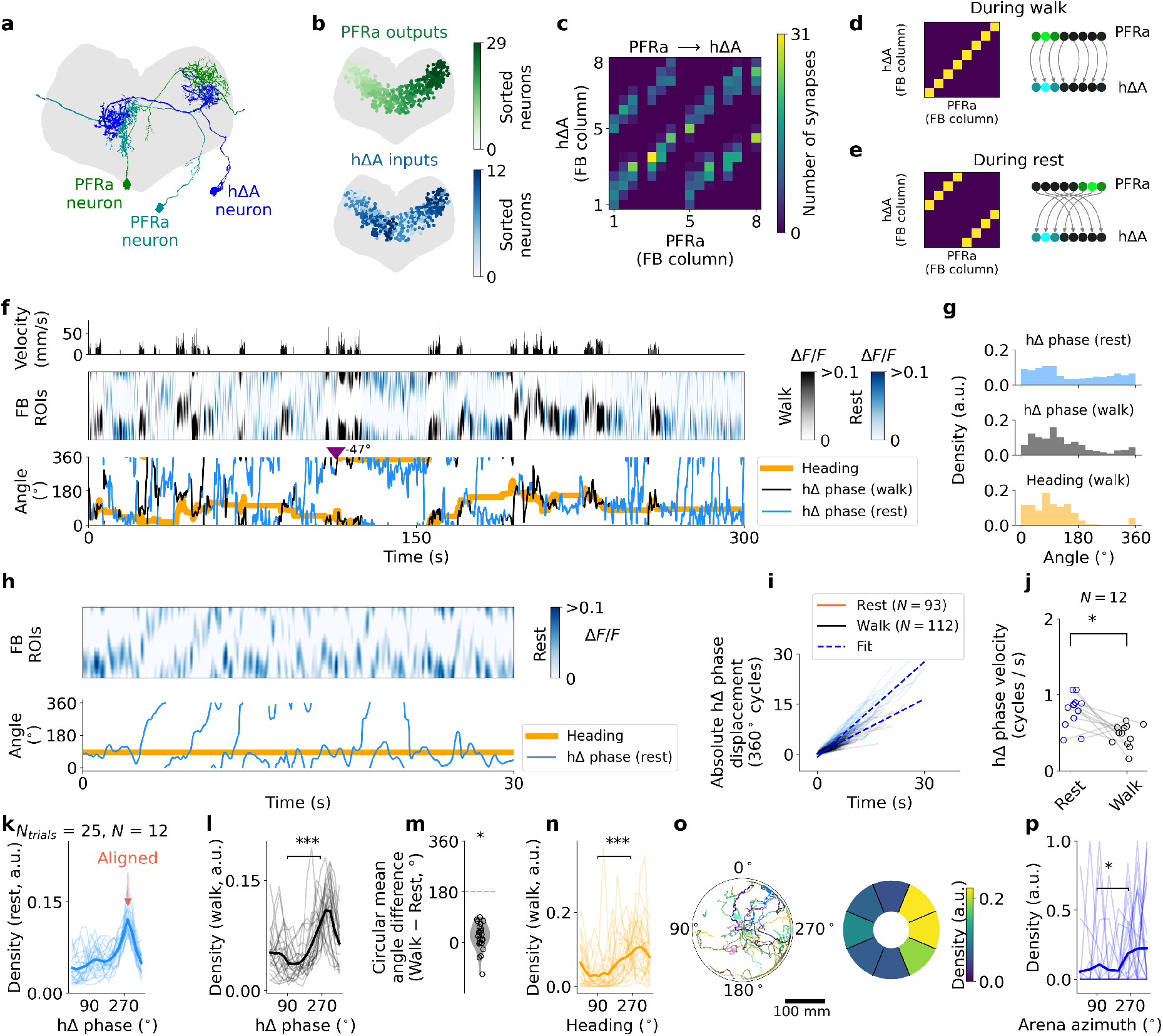
hΔ drift and shift correction. **a** A single hΔA neuron (red) receives input from two PFRa neurons in two different FB columns. Data from^77^. **b** Top: synapses in FB colored by presynaptic PFRa neurons, arranged by axonal arborization. Bottom: same synapses colored by postsynaptic hΔA neurons, sorted similarly. **c** Synapse counts between PFRa and hΔA neurons across FB columns. **d** During walking, PFR outputs propagate bump along the main diagonal (no shift). **e** During rest, PFR outputs propagate along the off-diagonal, shifting the bump by four columns or 180°. **f** Example hΔ calcium activity: Top: fly velocity. Middle: hΔ activity across 16 ROIs during rest (blue) and walking (black). Bottom: hΔ phase during rest (blue), walking (black), and fly heading (orange). Inverted purple triangle indicates switching offset correction (see Methods). **g** hΔ phase distributions during rest (top), during walking (middle), and heading distribution (16 bins, bottom) for the fly in f. **h** hΔ activity drifts during 30 s of rest. Top: activity across 16 ROIs. Bottom: hΔ phase (blue) with heading (orange). **i** Absolute hΔ phase displacement for 12 flies during *≥* 5 seconds of continuous resting (blue) and walking (black) epochs, with linear fits (red, see Methods). **j** hΔ phase velocity during rest (blue) and walking (black) in cycles per second for 12 flies. **k** hΔ phase density during rest for all flies, maximum aligned to 270°. **l** hΔ phase density during walking after alignment. **m** Difference in circular mean hΔ phase between walking and rest. **n** Heading distribution during walking across 16 bins after alignment. **o** Fly trajectories (left) and trajectory density (8 bins, radial distance *≥* 50 mm) after alignment. **p** Arena azimuth density (8 bins, radial distance *≥* 50 mm) for all flies after alignment. Asterisks in panels j and m show statistical significance based on a paired two-sided t-test across flies (p < 0.05 (*), p < 0.0005 (***)). Asterisks in density panels (**l**, **n**, and **p**) indicate significant differences between 90° and 270° (Kolmogorov-Smirnov test: p < 0.05 (*), and p < 0.0005 (***)). All data are shown in Supplementary Figs. S41-S45.

The PFRa-hΔ connectivity pattern suggests that PFR neurons might route the activity bump through hΔ neurons in a state-dependent manner: along the main diagonal during walking (Fig. 4d), and along the off-diagonal (four columns shifted) during rest (Fig. 4e). Such pathway selection could arise from state-dependent co-activation of populations that selectively enhance the main diagonal route during walking (Supplementary Figs. S33 and S34), or the off-diagonal route during rest (see Methods, Supplementary Information and Supplementary Figs. S35 and S36). This mechanism could compensate the 180° phase shift, maintaining a coherent hΔ activity across behavioral states.

## Downstream hΔ neurons do not show a 180° phase shift between rest and walking

To test whether hΔ neurons exhibit drift during rest and compensate for the 180° phase shift, we performed calcium imaging in hΔ neurons during navigational behavior in the mechanical VR. To label hΔ neurons, we selected the line R67A04-GAL4 using NeuronBridge^56^ (see Methods and Supplementary Fig. S39) and crossed it with UAS-sytjGCaMP7f to record localized activity in hΔ axons. We hypothesized that the 180° shift in bump position would be corrected in axons, since dendrites receive the output of PFR neurons with the observed 180° phase shift. In these experiments, flies were allowed to explore the mechanical arena for 5 minutes per trial, across three trials, as in Fig. 1.

hΔ neurons also exhibited a cosine-like activity bump (Supplementary Fig. S40), and its phase closely tracked heading direction during walking (Fig. 4f; Supplementary Figs. S41–S45), although with lower correlation than EPG or PFR neurons (Extended Data Fig. 2d). We again observed persistent drift during rest, however with similar phase distributions between rest and walking (Fig. 4g). The hΔ phase drifted over long rest periods, as observed in PFR neurons (Fig. 4h). Although hΔ phase velocity was slower than that of PFR neurons (compare Fig. 4i–j with Fig. 1h–i), likely due to the larger size of the filter required for noisier hΔ calcium signals (see Methods), the drift during rest was still nearly twice as fast as during walking (Fig. 4j), as observed for PFR neurons (Fig. 1i).

To compare hΔ activity during rest and walking across trials and flies, we applied the same analysis used for PFR neurons (Fig. 1) and removed arbitrary phase offsets, setting the rest-phase peak distribution to 270° (Fig. 4k). After alignment, hΔ phase distributions during walking closely matched those during rest (centered around 270°, Fig. 4l). The difference between rest and walking was around 0° (Fig. 4m), while the headings aligned by the rest-phase offset (Fig. 4n) and the equally aligned trajectories (Figs. 4o and p) were centered also at 270°. Thus, after aligning trajectories by the rest-phase offset, flies walked largely in the same direction as the peak of the hΔ phase during rest.

hΔ axonal activity shows similar distributions during rest and walking, despite the 180° shift in presynaptic PFR neurons. This shows that drift during rest is not a mere epiphenomenon, but could be utilized through the PFR-hΔ connectivity to maintain coherent activity distributions across behavioral states. This mechanism therefore could reactivate the FB columns active during walking for memory consolidation during rest (see Methods and Supplementary Information).

## Ring attractor network produces drift and 180° phase shift

Ring attractor networks naturally sustain a localized activity bump, which can drift due to connectivity asymmetries or noise^57–59^. To model the bidirectional 180° shift and drift during rest in PFR neurons, we implemented a ring attractor network with short-term synaptic depression. During walking, sustained external input stabilizes the activity bump but weakens the recurrent excitatory connections of the active neurons via synaptic depression. During rest, in the absence of external input, these weakened connections impair the ability of previously active neurons to maintain the bump. Consequently, the activity drifts toward the less recently active population (180° opposite on the ring), whose connections remain strong, while the previously active neurons recover (Fig. 5a and Supplementary Information). Consequently, this network tracks external input during walking but drifts during rest, resulting in 180°-shifted activity distributions between these two states (Fig. 5b and c).

**Figure 5.**
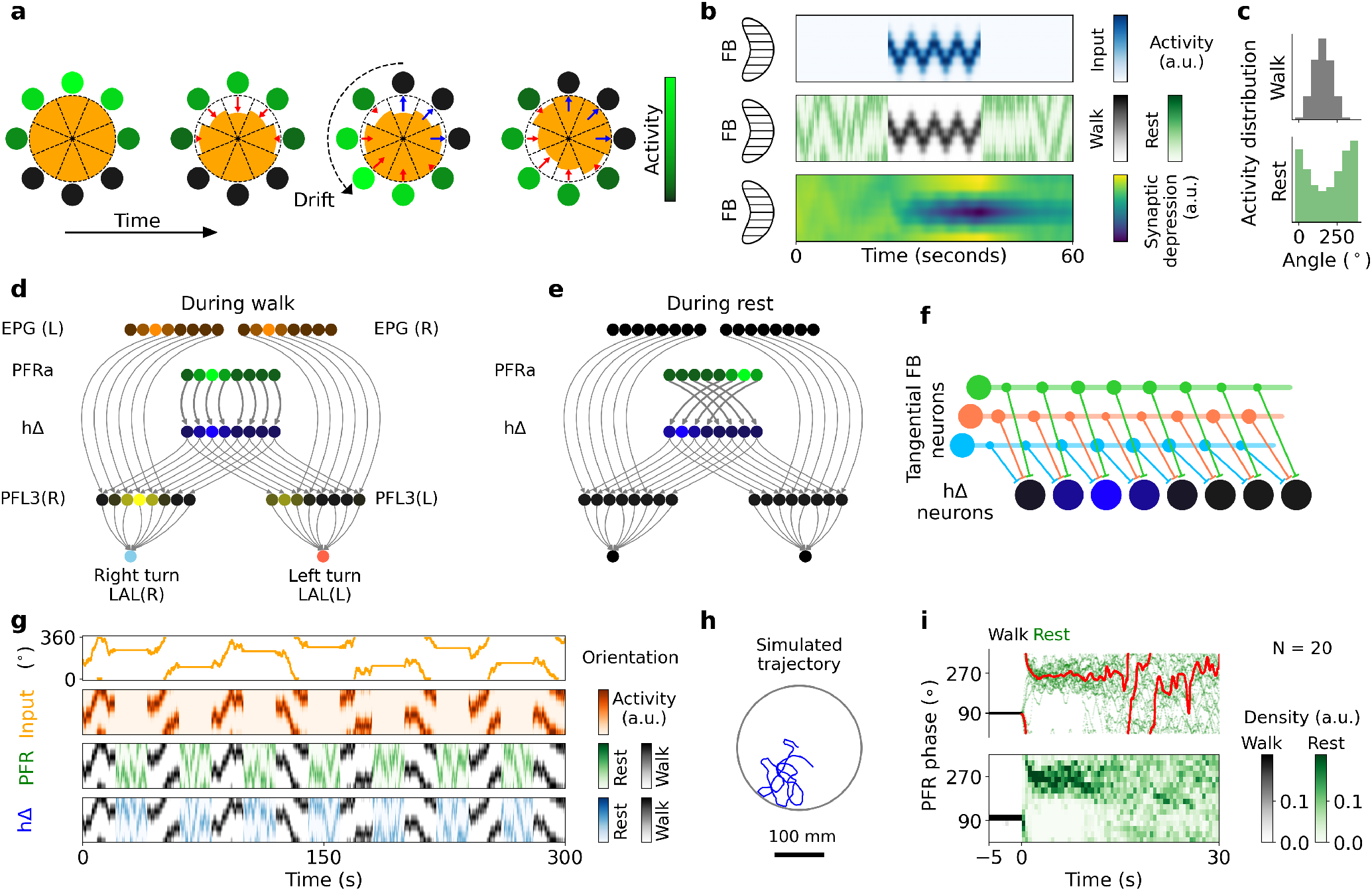
Ring-attractor model produces drift and 180° shifts in activity. **a** Neurons form a ring with activity (green) and neurotransmitter availability (orange wedges). During rest, sustained firing induces synaptic depression, shifting the activity peak to neurons with more available transmitter. **b** During walking, PFR neurons (middle, black) track heading input (EPG via hΔB; top, blue); during rest without input, activity drifts (green) due to synaptic depression (bottom). **c** Activity distributions during walking (top) and rest (bottom). **d** FB network architecture during walking: PFR neurons project column-to-column onto hΔ neurons. EPG and hΔ inputs drive PFL3 neurons, which control turning. **e** FB architecture during rest without input or output: PFR neurons project onto hΔ neurons with a four-column shift. **f** Tangential FB neurons selectively inhibit hΔ columns, encoding different goal directions. **g** Simulated FB activity: fly orientation (top), PFR input (second row), PFR activity during walking (black) and rest (green), and hΔ activity during walking (black) and rest (red). **h** Simulated trajectory of a fly in a circular arena. **i** Simulated PFR phase transitions from walking fixed at 90 ° (black) to rest (green). Top: simulated PFR phase trajectories (red line denotes circular average). Bottom: simulated PFR trajectory density.

## Connectome-based model reproduces experiments

We integrated the ring attractor model into a connectome-based FB circuit model to reproduce our experimental results. In this circuit, EPG neurons encode heading and transmit this signal to PFR neurons during walking. During rest, heading input is assumed to be zero—consistent with weakened EPG activity during rest periods^30^ (Supplementary Figs. S1-S3)—and PFR activity drifts with a 180° phase shift due to synaptic depression. PFR neurons are predicted to be dopaminergic^60^; we therefore assumed them to be excitatory, a choice supported by a qualitative connectivity analysis (Supplementary Information and Supplementary Fig. S46). PFR neurons transmit heading signals to hΔ neurons through the main diagonal during walking (Fig. 5d) and through the off-diagonal during rest (Fig. 5e), effectively correcting the 180° phase shift across behavioral states. hΔ neurons provide strong output to PFL3 neurons, which compute the error between the current heading (from EPG neurons) and the goal^15^. Therefore, in our model, hΔ neurons encode the goal direction, while tangential neurons modulate this signal through inhibition to drive the fly toward preferred directions^16^ (Fig. 5f). PFL3 neurons then compute the heading-goal error to drive corrective turns during walking (Fig. 5d; Supplementary Information).

We simulated the circuit in a closed-loop 2D arena, similar to our experiments, with alternating walking and rest periods (Fig. 5g, h). In this unified model, PFR activity tracks heading input (EPG, via hΔB) during walking and transmits it to hΔ via the main diagonal. We simulated the firing of FB tangential neurons as random inputs that modulate hΔ activity (Fig. 5f, see Supplementary Information), effectively driving the simulated fly toward random directions. In this closed-loop simulation, the fly turns to correct deviations between the instantaneous goal encoded in hΔ and the heading, and hΔ activity closely follows the heading direction during walking (Fig. 5g, last row). During rest, synaptic depression causes PFR activity to drift toward less recently active neurons, while hΔ receives this signal via the off-diagonal, effectively compensating for the PFR phase shift.

We simulated 20 flies navigating randomly for 5 minutes and applied the same analysis as to our experimental data. Aligning the simulated PFR phase distribution during rest to 270° yielded peaks in PFR phase during walking at 90°, as well as in hΔ phase during both rest and walking (Extended Data Fig. 9a-d). Heading distributions and aligned trajectories also showed peaks biased toward 90° (Extended Data Fig. 9e–g), demonstrating that the simulation replicates our experimental findings (Fig. 1, 4; see Supplementary Information).

We also simulated optogenetic perturbations to probe walking–rest transitions in the ring attractor model. Sustained input at 90° for 10 seconds during walking induced an immediate drift in the model toward 180°, maintained for several seconds — again matching experimental observations (Extended Data Fig. 5g). The duration of rest-phase activity was determined by time constant of synaptic depression (see Supplementary Information).

## Discussion

The importance of the CX for navigation learning and memory has long been recognized^25–29,61^ and corresponding circuit mechanisms have been proposed based on computational models as well as behavior, physiology, and connectivity data^4,8,16,54,55^. We found that a population of PFR neurons is sufficient for driving direction selection. Direction memory emerges during learning over time through changes in the rest-state distribution. The 180° shifted distributions between rest and walking point to a dedicated computational role of rest states for navigation. Supporting this, downstream hΔ neurons do not show the 180° shifted distributions, suggesting that they correct the inverted drift through their connectivity pattern with PFR neurons. Thus, the CX offers an architecture for the close interaction of head direction, direction selection, and memory consolidation during rest and walking.

Optogenetic activation during either rest or walking drives PFR activity toward a 180°-shifted direction (Extended Data Fig. 5, Fig. 2m). This bidirectional effect indicates that neural dynamics are shaped by activation history, consistent with a synaptic depression mechanism, as also modeled in the ring attractor network, which equally displays drift during rest^57–59^ (Fig. 5).

Neural representations underlying visual memory behavior have been investigated using two-photon imaging in tethered flies during behavior^14,35,36,62^. These experiments show that neurons across CX substructures are important for learning, such as ring neurons that detect visual features^35,63–65^, or EPG neurons^14,36^ which represent the fly’s heading angle^30^. Goal-related columnar neurons in the fan-shaped body have been identified using optogenetics^14^. However, the connectome shows only weak direct interactions between these and PFR neurons, which could point to the integration of multiple neural populations for goal-directed navigation.

hΔ neurons provide strong input back to PFR, which could support bump maintenance and ring attractor dynamics (Supplementary Fig. S4d and Supplementary Information), and to PFL neurons (Tables S3 and S4), which compute the error between current heading and goal to drive turning^14,15^. hΔ neurons are therefore well positioned to encode instantaneous goal direction^16^, consistent with their weaker heading tuning relative to EPG or PFR neurons during walking (Extended Data Fig. 2d).

Inputs to hΔ neurons from FB neurons are either restricted to the main diagonal, the off-diagonal, or both, suggesting distinct routing of bump transmission (Supplementary Figs. S33–S38). We identified tangential FB neurons that respond to cooler temperatures, although they likely do not connect directly to hΔ neurons; other tangential neurons signal hunger^52^. Models propose that distinct tangential FB populations modulate hΔ neurons according to behavioral needs^4,16,54,55^. Our data suggest that PFR drift during rest reactivates recently used hΔ columns, potentially reinforcing preferred walking directions through coactivation with tangential FB neurons. Consistent with this idea, temperature-sensitive tangential FB neurons also display spontaneous activity during rest (Extended Data Fig. 8g,h).

In mammals, grid and place cells move towards goal locations following learning^5,6,8,18^. Such distortions could originate from local modifications of “Mexican hat” connectivity patterns^18^. Similarly, local modification of connectivity could result in the shaping of drift distributions observed here. In mice, changes in head direction dynamics have been observed during walking in darkness – although not during learning – revealing a memory of the direction of a previous visual stimulus^66^. This suggests that potentially similar representations could be used across species to remember salient or behaviorally important directions.

Activity during rest has been described in other *Drosophila* neurons^67^ and glia^52^. Reactivation of neurons in the FB after learning has also been associated with memory consolidation^37^. In mammals, head direction representations drift during sleep^68,69^. Neural populations involved in particular in navigation show autonomous activity during rest and sleep, which plays a role in planning, decision-making, and learning^19–21,23,24,70^.

In the proposed model PFR drift during rest is corrected by downstream hΔ neurons through specific PFR-hΔ connectivity. During walking, hΔ neurons integrate PFR and tangential FB inputs to encode goal direction. During rest, drifting PFR activity reactivates the same hΔ columns active during walking, potentially supporting offline consolidation of walking history^71,72^.

Although hypothetical, the proposed mechanisms have supporting evidence. First, while synaptic depression within PFR neurons was not measured directly, activating PFR neurons at rest or during walking was followed by immediate inhibition of the stimulated columns (Fig. 2m, Extended Data Fig. 5g), consistent with depression at the activated synapses. Attractor dynamics may arise from Δ7 neurons, the only PB inputs of PFR neurons (see Supplementary Information), which support ring attractor dynamics in EPG neurons through sinusoidal inhibition. Finally, the connectome predicts state-dependent routing of PFR output to hΔ neurons, consistent with the 0° shift in hΔ phase between rest and walking. Stimulating PFR columns while imaging hΔ axonal activity would directly test the proposed routing switch.

Together, our data suggest that drift activity is leveraged by a fan-shaped body circuit to shape navigation and memory consolidation during rest, analogous to replay in reinforcement learning^36,73,74^. This offline reactivation could strengthen walking heading directions, suggesting similarities between mammalian and insect navigation systems beyond basic ring attractor dynamics^72,75,76^, including drift, changes in spatial representations following goal learning, and changes in dynamics between rest and walking^5,18,66^.

## Acknowledgements

We thank Anna Kaatz for fly crosses and dissections for imaging experiments. We thank Ivan Vishniakou for technical support. We thank Gülin Öykü Tabel for fly crosses, Jasmin Beinert, Anne Buecker, and her team for help with maintaining flies, and Vivek Jayaraman for Chrimson flies. We thank Jason Kerr for the Mai Tai Laser as well as other equipment, and Anne Buecker, Christoph Geisen, Alexandr Klioutchnikov, and Yazid Rachad for helpful discussions. We thank the MPINB mechanical and electronics workshops for support with building instrumentation.

## Author contributions

AFV and JDS designed the study and wrote the manuscript. All authors designed and built the setup, RH built mechanical arena, AFV built automation and control. AFV performed experiments, data analysis, figures, and computational modeling.

## Methods

### Drosophila preparation

All flies used for experiments were female and were reared in an incubator at a temperature of 22 degrees Celsius and under a 12-hour light/dark cycle. Flies between 3 and 8 days old were used for imaging. The GAL4 line R60D05 (Bloomington #39247) was used to express GCaMP8m (Bloomington #92591) in EPG neurons. The split line SS02270 (Bloomington #75925), or the GAL4 line R27G06 (Bloomington #54779) was used to express GCaMP8m in PFR neurons. The GAL4 line R67A04 (Bloomington #39396) was used to express sytjGCaMP7f (Bloomington #605095) in hΔ neurons. The GAL4 line R72G07 was used to express GCaMP8m in heat-sensitive neurons (Bloomington #49622). The dissection of the flies for inserting a transparent window into the cuticle was performed as described^79^. First, a window was cut in the head using a laser. Then the cuticle and air sacs were removed under a dissection microscope using a microrobotic arm^79^ or manually. The opening was then sealed with a drop of transparent UV glue (Freeform, UV fixgel composite, purchased before 2023, or NOA68 optical glue). Usually, between 3 and 5 flies were dissected at a time. prior to any experiment was performed. Flies were allowed to recover after the dissection in isolation, in vials with food for 2 to 3 days before imaging. The flies were then glued to a glass slide with UV glue and transferred to the mechanical VR setup and placed under the two-photon microscope^42^. Immediately after positioning the flies under the microscope and in the VR setup, visual inspection was performed for the behavior of the fly and the imaging quality. Flies with low imaging quality, for example where the FB was only partially visible or too dim due to residual connective tissue not fully removed from the brain during surgery, were discarded. Similarly, flies exhibiting poor walking behavior on the air suspended ball, for instance pulling up and down the ball without coordinated walking patterns, were also excluded. This filtering occurred before any experiment was performed. Since flies are sealed after dissections with glue, and are irreversibly mounted with glue to a cover slide, adjustments to the dissection or posture of the fly with respect to holder are not possible, different from approaches with saline-based methods^43^.

**Extended Data Fig. 1.**
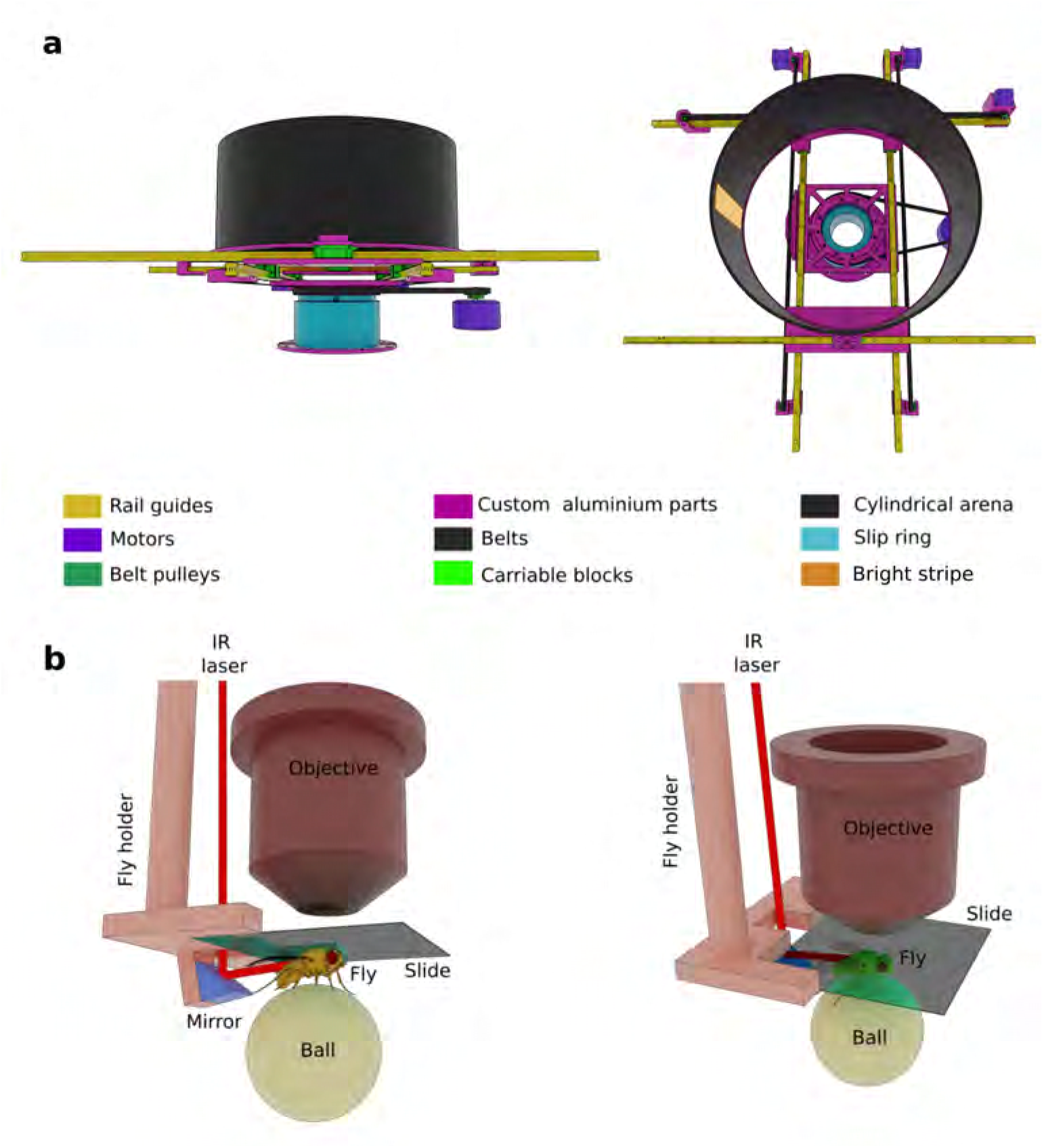
**a** Mechanical virtual reality setup from two different viewpoints, colors are for visualization. The setup is described in detail in^41^. **b** A fly glued to a cover slide is attached to the fly holder and placed between the air-supported ball and the objective. The IR laser used as an aversive stimulus is focused on the back of the fly through an opening in the fly holder (visible on the left side) and with the help of a small mirror.

**Extended Data Fig. 2.**
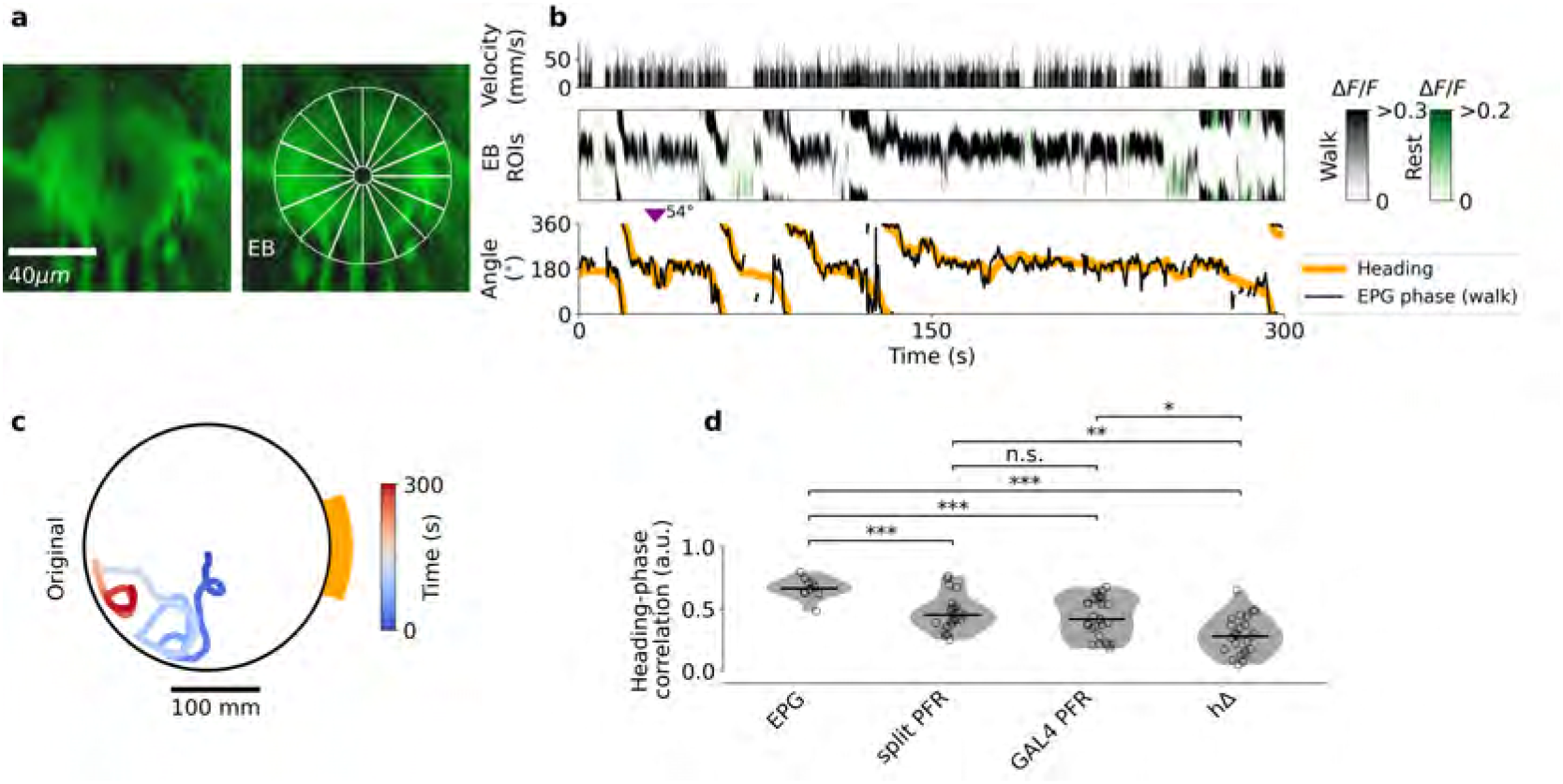
**a** Expression of calcium indicator in EPG neurons (left) and defined ROIs across the ellipsoid body (EB, right). **b** Example of EPG calcium activity during navigation. Top: fly velocity. Middle: EPG activity across 16 ROIs (panel a) during rest (green) and walking (black). Bottom: EPG phase during walking (black), and fly heading (orange). Inverted purple triangle indicates switching offset correction (see Methods and Extended Data Fig. 3). **c** Fly original trajectory in the mechanical VR during 300 seconds. **d** Heading-phase correlation during navigation in EPG (Supplementary Figs. S1–S3), PFR (using the split line SS02270; Supplementary Figs. S7–S11, and the GAL4 R27G06 line), and hΔ neurons (Supplementary Figs. S41–S45). Violin plots show the distribution of the heading–phase correlation per group, with each dot representing a single fly; black lines indicate the group median. Group distributions were compared pairwise using the two-sample Kolmogorov–Smirnov test (two-sided); p-values were corrected for multiple comparisons across the six pairwise tests using the Holm–Bonferroni method. Significance is indicated above each pair: *^∗^ p <* 0.05, *^∗∗^ p <* 0.01, *^∗∗∗^ p <* 0.001 (Holm-adjusted); n.s., not significant (adjusted *p ≥* 0.05).

### Mechanical virtual reality

We combined a mechanical virtual reality (VR) setup similar to the one described in^41^ with a custom two-photon microscope (as described in^79,80^) for recording neural activity during learning.

The mechanical VR allows flies to navigate within a cylindrical arena that both translates and rotates around the fly. It consists of two main components: a rotational stage with a slip ring for continuous rotation and a translation stage that offsets the arena along two dimensions. We used a cylindrical arena twice the size of that in^41^, with a diameter of 500 mm and a height of 250 mm (Fig. 1a and b) . For translation, we employed the same Nema 17 stepper motors with encoders as in^41^. However, for rotation, we used a T-Motor AK10-9 V2.0 (60kv Brushless Motor) instead, to accommodate the increased inertia of the larger arena. Additionally, the translation stage was extended in both dimensions to support the required larger translations. The setup is shown in Fig. 1a, b, and Extended Data Fig. 1a. To prevent vibrations from the motors to interfere with fly behavior or two-photon imaging, the mechanical VR was mechanically isolated by placing it on a table that was mechanically decoupled from the optical table, as described in^41^.

The cylindrical arena contained either a single vertical stripe (Figs. 1, 2, 4 and Extended Data Figs. 4 and 5), measuring 40 mm in width and 200 mm in height, or two vertical stripes (one bright, one dim) positioned 180*^◦^* apart for learning experiments (Fig. 3a). The stripes could be illuminated from the back of the cylinder using white LEDs. The LED light was diffused through white paper and filtered using colored long-pass filters (Thorlabs, FGL550S) to prevent interference with the photomultiplier tube (PMT) used for fluorescence detection. During the learning experiment (Fig. 3a), the LEDs illuminating each stripe were independently controlled via an Arduino UNO.

**Extended Data Fig. 3.**
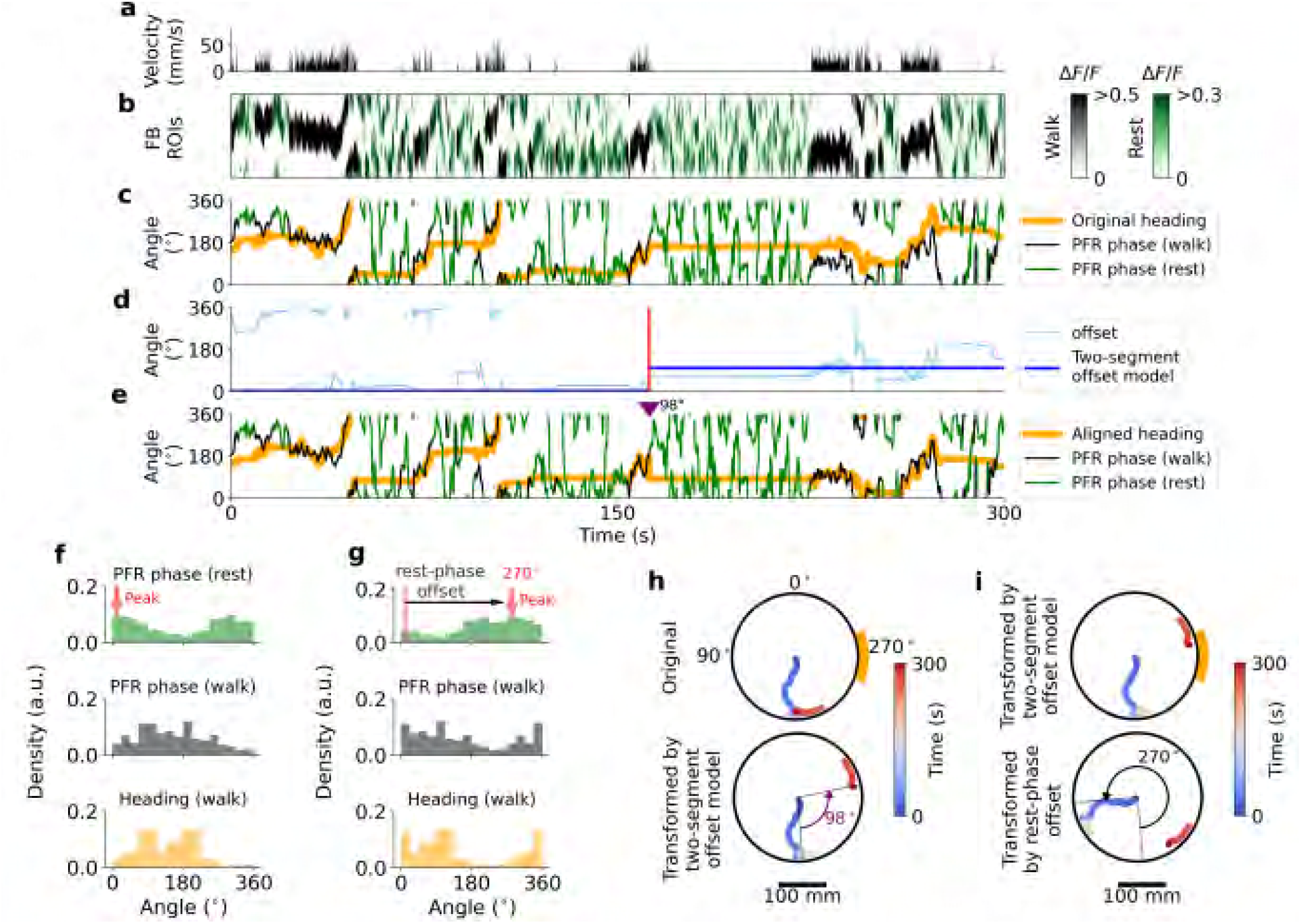
Analysis pipeline for PFR phase and behavioral data. **a** Fly velocity during a 5-minute mechanical VR trial. **b** PFR neuron fluorescence during walking (black) and resting (green) epochs. **c** Original fly heading (orange) aligned with the extracted PFR phase during walking (black) and resting (green). **d** The offset between the walking PFR phase and fly heading (light blue), modeled as a two-segment constant offset (dark blue) to account for phase jumps. **e** Heading direction corrected using the two-segment offset model (orange), overlaid with the PFR phase (walking: black; resting: green). **f** Phase distributions: PFR phase during rest (top), PFR phase during walking (middle), and heading direction (bottom). The peak of the resting PFR phase is identified (red arrow; e.g., at 0*^◦^*). **g** Data realigned using the rest-phase offset. The resting PFR phase peak is shifted to 270*^◦^* (top), and this same offset is applied to the walking PFR phase (middle) and heading (bottom) distributions. **h** Trajectory transformation. Top: Original fly trajectory. Bottom: Trajectory transformed using the two-segment offset from d. This rotation aligns the mechanical VR azimuth origin (0*^◦^*) with the PFR phase zero-point. The offset jump at *t*_switch_ effectively rotates the arena by 98*^◦^*, shifting the fly’s apparent position. **i** Final transformed trajectory. The trajectory from h is further rotated by the rest-phase offset (bottom), resulting in the final aligned trajectory shown in Supplementary Figs. S7-S11.

For imaging experiments, flies were glued to a cover slip and positioned on an air-suspended ball using a custom aluminum fly holder (Extended Data Fig. 1b). The ball was placed 150 mm above the bottom of the cylindrical arena (Fig. 1b). Fly movement was tracked using an optic flow-based algorithm^81^, which analyzed ball motion via a small mirror from the back and a camera from below, as described in^41^. To prevent collisions between the objective and the cylinder walls, a walking limit with a 270 mm diameter was set in the mechanical VR, as in^41^. Thus, the effective walking area for the flies was confined to a circular region of 270 mm diameter. The rotational gain of the arena was set to 0.3 to prevent fast turns in the mechanical virtual reality which could lead to excessive strain on the mechanical parts due to the large diameter and corresponding rotational path; the translational gain was set to 1. We validated the mechanical VR and closed-loop imaging using EPG neurons, which track the heading of the fly^30^(Supplementary Figs. S1–S3). The rotational gain affects only the visual motion experienced during walking; during rest the scene is stationary, so the drift, the 180*^◦^* offset, and the effect of rest stimulation on subsequent walking do not depend on the gain setting. Despite the low rotational gain, most flies were able to track the position of the stripe (Extended Data Fig. 2d). Flies that were not able to track the stripe, showing low correlation between neural tracking and heading in the arena, were discarded (see Table S1 and sections below for details).

**Extended Data Fig. 4.**
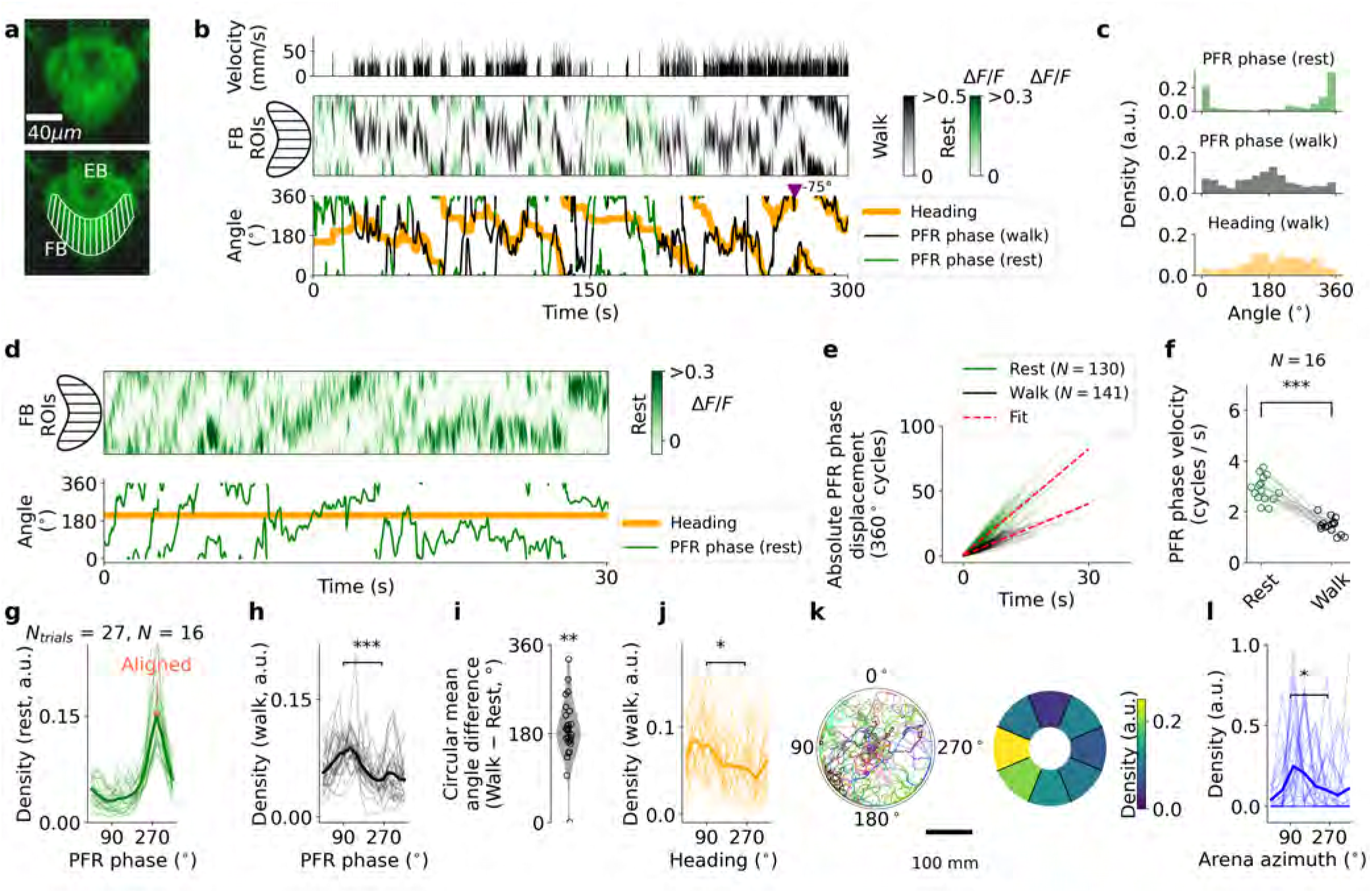
PFR drift in the R27G06-GAL4 line. **a** Expression of calcium indicator in PFR neurons with R27G06-GAL4 (average of 500 frames), which also labels other populations (in the ellipsoid body, EB). Right: ROIs across the FB. Panels **b-l** are similar to Fig. 1e-o, respectively. Asterisks in panels f and i show statistical significance based on a paired two-sided t-test across flies (p < 0.005 (**), p < 0.0005 (***)). Asterisks in density panels (h, j, and l) indicate significant differences between 90*^◦^* and 270*^◦^* (Kolmogorov-Smirnov test: p < 0.005 (**), p < 0.0005 (***)).

**Extended Data Fig. 5.**
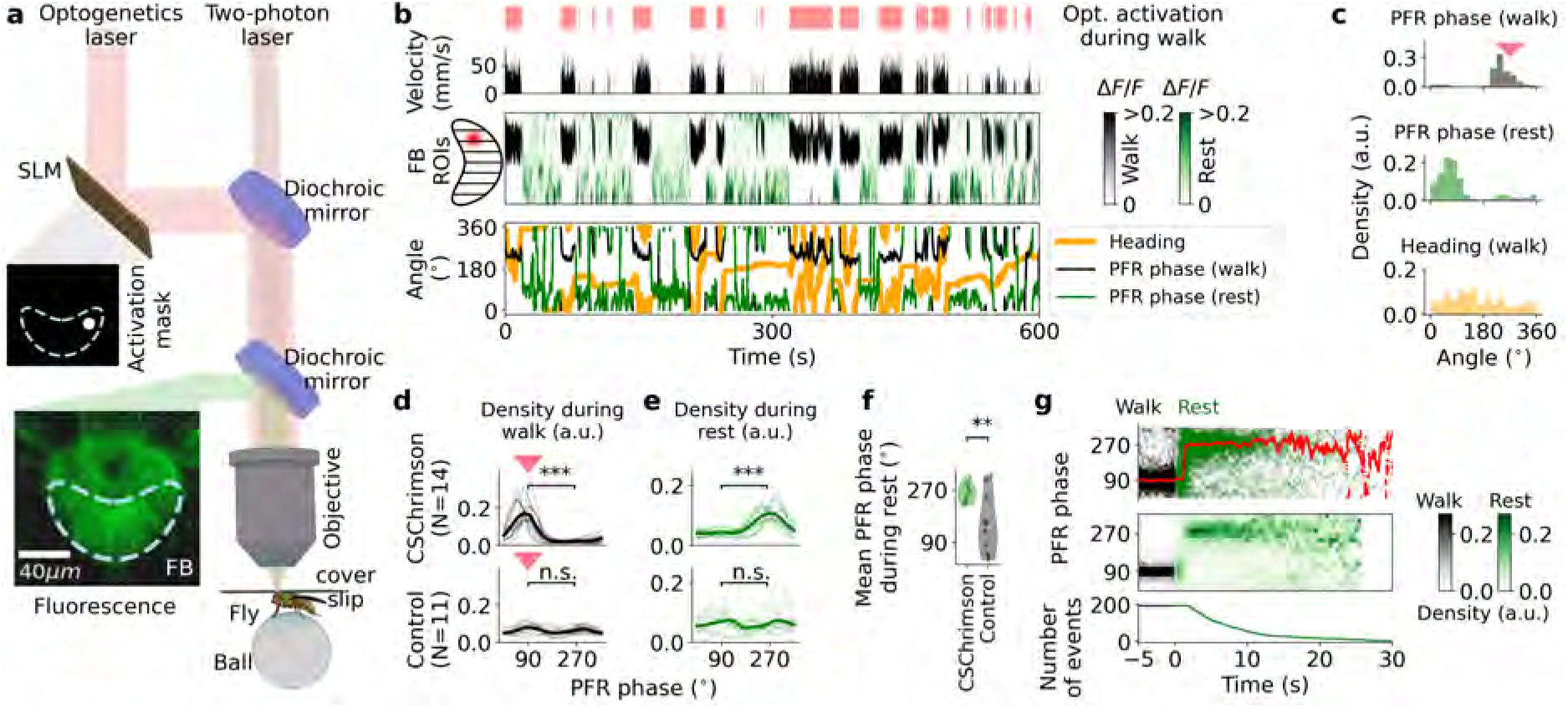
Optogenetic activation during walking drives subsequent activity during rest. **a** Setup for combined local optogenetic activation and two-photon calcium imaging in behaving flies. A red laser is reflected off an SLM and activation patterns as well as a laser for two-photon imaging are targeted to the fly brain. **b** Optogenetic activation of PFR columns during walking in navigating flies. Top: laser for optogenetics is only switched on when the fly walks (walking velocity *≥* 0). First row: walking velocity of the fly. Second row: fluorescence from 16 ROIs defined in the FB. Optogenetic activation was performed in columns on one side of the FB (shown on the left). Third row: phase of PFR neurons (see Methods), during walking (black) and rest (green). Orange: fly heading **c** Top: PFR phase distribution during walking. Red triangle indicates the corresponding optogenetic activation site. Center: PFR phase distribution during rest. Bottom: Fly heading distributions across 16 bins during walking. **d** PFR phase distribution during walking for CsChrimson flies (top) and controls (bottom). **e** Same as d, but during rest. Asterisks in density panels (d and e) indicate significant differences between 90*^◦^* and 270*^◦^* (Kolmogorov-Smirnov test: p < 0.005 (**), p < 0.0005 (***)). **f** Mean PFR phase during rest: CsChrimson (green) vs. controls (black). Each dot represents a single fly. Asterisks represent statistical significance between Chrimson and control flies (Two-Sample Kuiper Test: p < 0.005 (**)). **g** PFR phase transitions from walking (black) to rest (green): PFR phase trajectories (top), density (middle), and transition events count (bottom). Transitions are defined as *≥* 5 seconds of walking followed by *≥* 2 seconds resting. Data are shown in Supplementary Figs. S13-S15 for all Chrimson flies, and in Supplementary Figs. S16-S18 for all control flies.

### Two-photon microscope

We used a custom two-photon microscope setup similar to the one described in^79,80^. This setup contains two axially offset beams with independently controlled focal planes that were offset in time by 6 ns to additionally detect them independently using temporal multiplexing. This setup configuration allows estimating and correcting axial motion^80^. Here we considered both beams as a combined excitation volume with an effective extended focal length, allowing us to record volumetric calcium activity while decreasing two-photon laser power for imaging. Axial motion estimation and correction were therefore not considered here, as axial motion during imaging produced little fluorescence changes in the recorded neurons.

### Data acquisition

Behavioral data from the mechanical VR was obtained at 100 Hz, providing the time stamps of the experiment, the ball tracking data, and the position and orientation of the mechanical arena. The time stamps were sent through ROS 1 (Robot Operating System Version 1) and through the local network to the computer running Scanimage that controlled the two-photon microscope. These time stamps were stored together with each recorded frame from the microscope and were later used to synchronize the behavior and imaging data. The imaging data were obtained at a resolution of 256 *×* 256 pixels at approximately 60 Hz continuously during the whole experiment.

**Extended Data Fig. 6.**
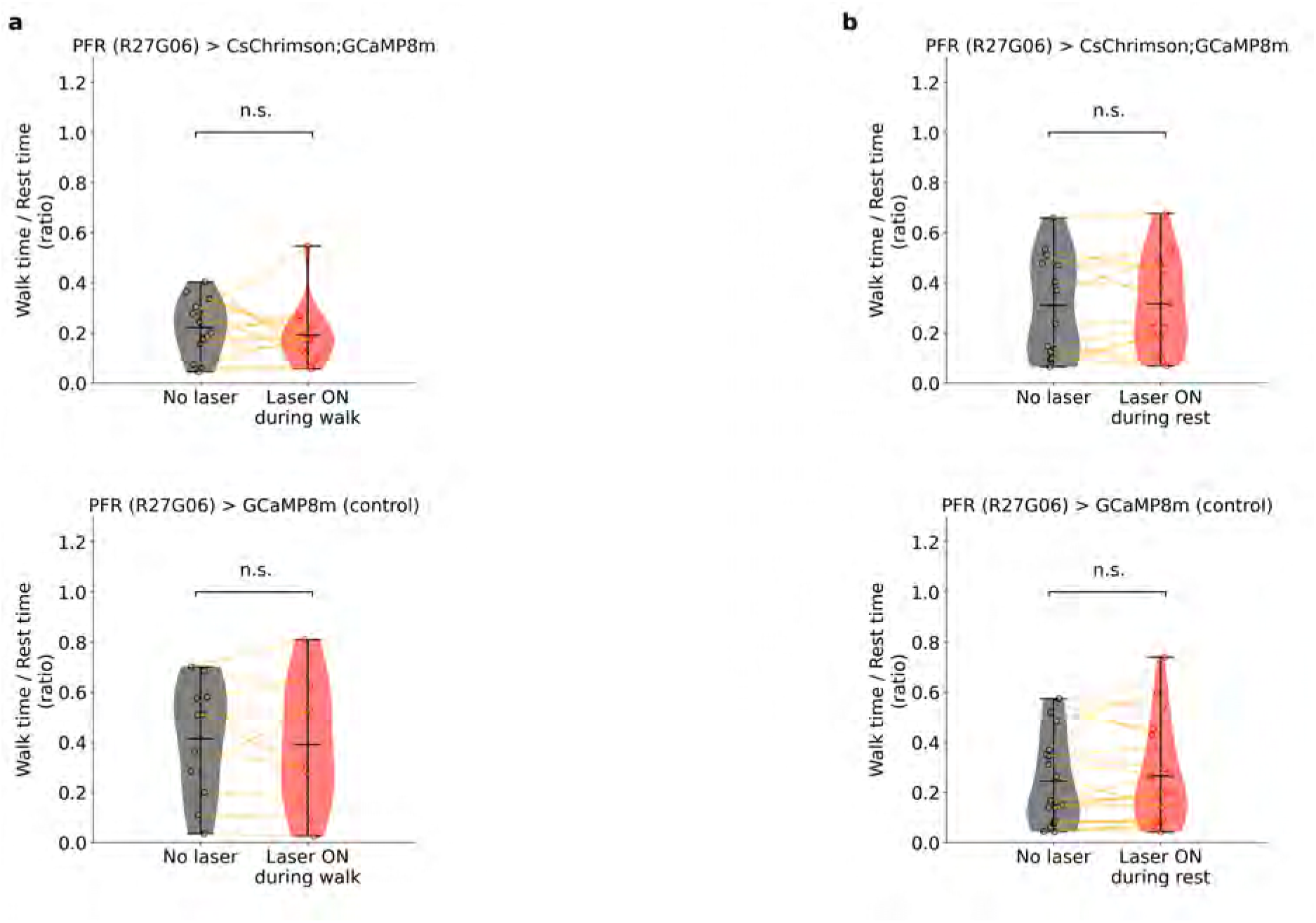
Effect of optogenetic red laser on fly behavior. **a** Top: Distribution of the rest-to-walk time ratio for flies expressing CsChrimson, comparing behavior without laser (laser off) and with laser activation during walking periods (laser on). Each dot represents an individual fly; lines connect the same fly across conditions. Bottom: Same comparison for control flies expressing only GCaMP8m. **b** Same as (a) but for laser activation triggered during rest periods. No significant difference between laser-on and laser-off was observed (paired two-sided t-test).

### Analysis of behavior data

The linear velocity of the fly was first computed from the recorded ball velocity. We then thresholded the linear velocity to define bouts of rest (with zero linear velocity) or walking for each fly. This threshold was set to 0.125 mm*/*sec, which was adjusted to be right above the linear velocity noise level.

### Analysis of imaging data

For each fly, all recorded imaging frames were aligned to a common template to correct for lateral motion using cross-correlation. Once every frame was aligned, we defined a total of 16 regions of interest (ROIs) along the expression pattern in the FB (Fig. 1d) or in the ellipsoid body (EB, Extended Data Fig. 2a), and computed the total intensity *I_i_*(*t*) within each ROI *i* and for each frame, associated with a time stamp *t*. Then, the fluorescence over the frames for each ROI was computed as

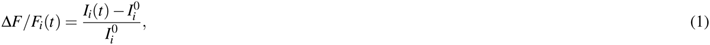

where *I_i_*^0^ is the 0.1 quantile of the intensity *I_i_*(*t*), calculated using a sliding window containing intensity data over 300 seconds. The window was continuously shifted along the timeline to compute a dynamic baseline, enhancing the signal of the activity bump compared to using a fixed baseline over the entire experiment. To better visualize local activity in the FB (e.g., the activity bump) independent of global fluctuations, we subtracted the mean fluorescence across all 16 ROIs from each ROI signal at each time point. This normalization accounts for global intensity increase in the whole FB or EB, often caused by locomotion, effectively holding the average fluorescence constant over time.

**Extended Data Fig. 7.**
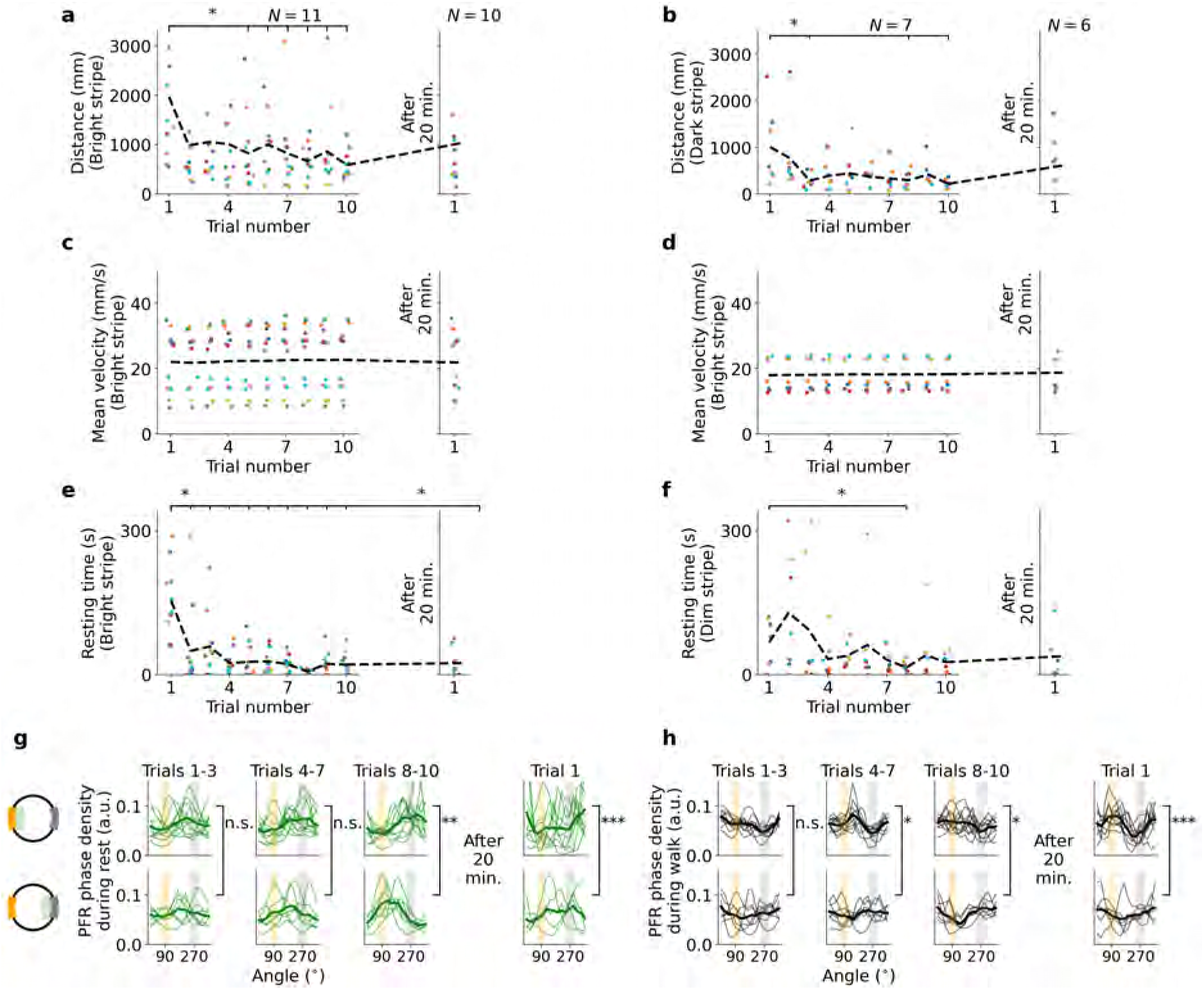
Behavior during learning tasks and PFR phase distributions. **a,b** Walked distance per trial for flies during the learning task with the goal located at the bright stripe (a) or dark stripe (b). Each colored point represents an individual fly. **c,d** Mean velocity across trials for each fly when the goal was at the bright stripe (c) or dark stripe (d). **e,f** Resting time across trials for each fly when the goal was at the bright stripe (e) or dark stripe (f). **g** Top: PFR phase density during rest across trials 1–3 (left), 4–7 (center), and 8–10 (right) for all flies trained with the cool area in front of the bright stripe (at a 90*^◦^* direction, schematic on left). On the right, PFR phase density during rest over 1 trial after the 20-minute break. Bottom: same as top, but with cool area in front of dim stripe (at 270*^◦^* direction). Light green traces represent individual flies; thick green lines show averages. **h** Same as panel e but for PFR phase density during walking. Asterisks and connecting lines in **a–f** denote statistical significance (p < 0.05, two-sided t-test) relative to trial 1; absent lines indicate no significance. Trials with non-connecting lines to trial 1 are not statistically significant. Asterisks in panels g and h indicate statistical significance between distribution sets, using a two-sided t-test on the reduced distribution for each fly (projected to one dimension via PCA, see Methods). Significance levels: p < 0.05 (*), p < 0.005 (**), p < 0.0005 (***).

**Extended Data Fig. 8.**
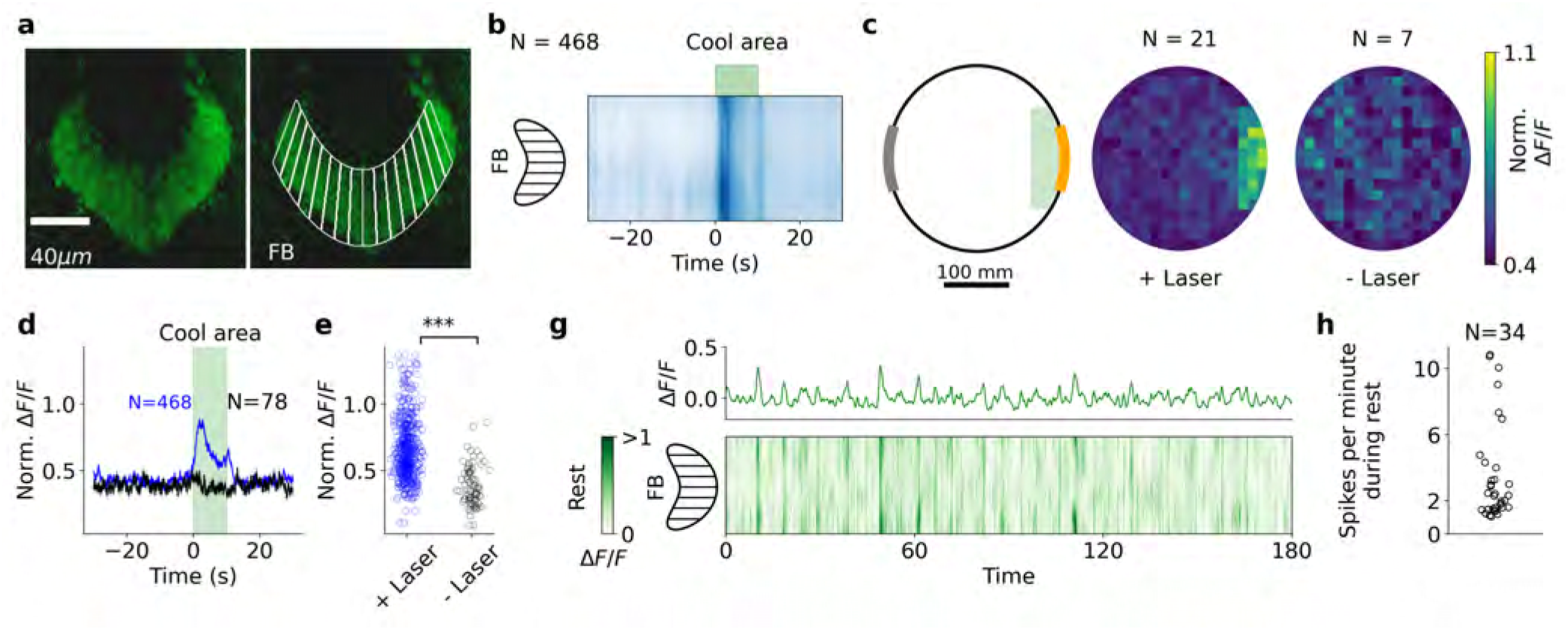
Temperature sensitive neurons in FB. **a** Neurons labeled with 72G07-GAL4 innervate the dorsal layers of the FB (top). Regions of interest in the FB (bottom). **b** Average fluorescence FB from 21 flies crossing the cool area 242 times, aligned to the time point they enter the cool area. Events where flies entered the cool area but did not remain for 10 seconds during training were truncated to the time they exited the cool area. Events where flies stayed in the cool area for 10 seconds triggered the start of a new trial, during which the cool area switched. As a result, flies were in the cool area for a maximum of 10 seconds. **c** Left: arena with cool area. Center: fluorescence of dorsal FB layer neurons at each position in the arena (arena position binned in 15 mm increments). Right: same as center panel but for flies where the IR laser was not used. **d** Amplitude of dorsal FB layer fluorescence when crossing the cool area, comparing conditions where the IR laser was used (blue line) versus when it was not used (control, black line). Thick lines represent average, while shaded areas indicate the standard error of the mean (SEM). Events where flies entered the cool area but stayed for less than 10 seconds during training were truncated to the time they exited the cool area. **e** Fluorescence levels of dorsal FB layer neurons in the cool area for flies with laser activation (blue, 21 flies) and without laser activation (black, 7 flies).

The resulting signal, denoted as 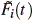, is calculated as:

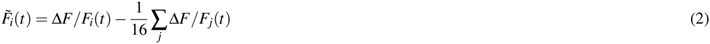

Once normalized fluorescence data was obtained for each ROI, we created a fluorescence matrix with dimensions *T ×* 16, where *T* is the total number of recorded frames for one fly. The matrix was filtered using a Gaussian filter. To do this, we first extended the fluorescence matrix along the second dimension, resulting in a matrix of size *T ×* 48, and a Gaussian filter with a size of 1 *×* 1 was applied for recordings of PFR neurons, whereas a larger filter of size 5 *×* 1 was used for recordings of hΔ neurons to reduce the higher noise levels associated with their dimmer fluorescence signals. Finally, we extracted the 16 ROIs from the center slices (16 to 32) of the extended fluorescence matrix (*T ×* 16: 32). This approach ensured that the Gaussian filtering was applied in a circular manner, effectively filtering together ROI 1 and ROI 16.

### Phase calculation from calcium activity

We calculated the phase for EPG, PFR and hΔ neurons from the fluorescence signals of 16 defined ROIs in the EB or FB (Fig. 1d). This was done by sliding a cosine function across the 16 ROIs, shifting one ROI at a time, and computing the Pearson correlation between the cosine function and the fluorescence profile of the ROIs at each frame. The EPG, PFR and hΔ phase was determined as the cosine offset with the highest correlation. Finally, the phase was converted to degrees by multiplying by 360/16. The EPG or PFR phase was further smoothed across time using a mean circular filter, calculating the average value over a sliding window of 0.5 seconds.

**Extended Data Fig. 9.**
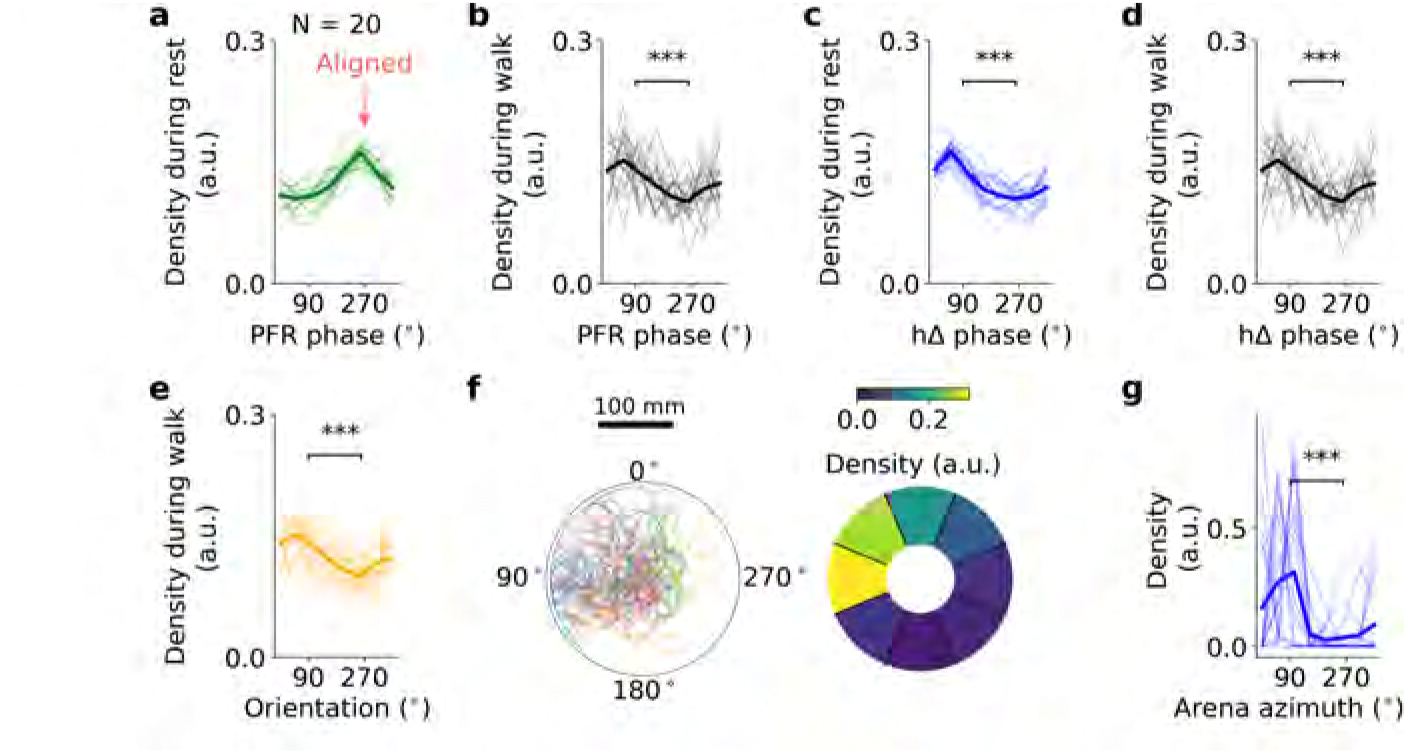
**a** PFR phase distributions aligned to 270*^◦^* across 20 simulated flies. **b** Distributions of PFR during walking. **c** and **d** Distributions of hΔ phases during rest and walking after PFR rest-phase offset alignment. **e** Fly orientation distributions, after rest-phase offset alignment. **f** Left: Trajectories of 20 simulated flies after rest-phase offset alignment (left) and trajectory density (right, 8 bins, radial distance *≥* 50 mm). **g** Arena azimuth across 8 bins for all flies aligned by PFR rest-phase offset. Asterisks denote significant differences between 90 and 270*^◦^* (Kolmogorov–Smirnov test, *p <* 0.0005, ***).

### Offset correction between phase and heading

The PFR and hΔ phase during walking reflects the fly’s traveling direction relative to visual landmarks but exhibits an arbitrary offset across individual flies^30^. To align the neural data with the orientation of the fly, we calculated the offset by first focusing on the phase during walking epochs, since the PFR and hΔ neurons drift during rest while the orientation of the fly remains constant. Therefore, we only considered the phase of PFR and hΔ neurons during active walking for offset computation.

The offset at any given moment is calculated as the difference between the phase from PFR or hΔ neurons and the fly heading in the mechanical VR:

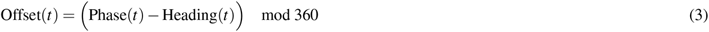

This offset is filtered using a circular mean filter with a sliding window of 15 seconds to smooth out noise and reduce variability. Extended Data Figure 3d (light blue line) shows an example of the calculated offset over time.

In some flies, we observed that the offset remained relatively constant for extended periods, but switched during certain portions of the experiment (see for example Extended Data Figure 3d). This phenomenon of offset switching has been observed when different visual cues are presented in virtual reality^30^, or in real-world virtual environments^49^. Given that our mechanical virtual reality system has a low rotational gain of 0.3 (i.e., the VR rotates 0.3 revolutions for every full rotation of the fly), it is possible that this low gain contributed to the offset switching observed in our experiments.

To model this switching offset, we allowed for a maximum of one switch during each experiment and aimed to divide the filtered offset into two distinct constant segments. The optimization approach we used for this task is based on minimizing the error between the filtered offset and the two-segment model of the offset. Specifically, we optimized the position of one breakpoint in time, *t*_switch_, that divides the experiment into two segments:

- The first segment spans from the start of the experiment (*t_min_*) to time *t*_switch_.
- The second segment spans from time *t*_switch_ to the end of the experiment (*t_max_*).

We modeled the offset in each segment as a constant value, so that the offset at any time *t* is given by:

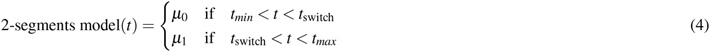

where *µ_i_*is the mean offset in the *i*-th segment. For each segment, the mean offset *µ_i_* is computed as the circular average of the filtered offset between the defined times.

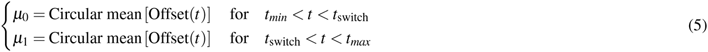

The optimization objective is to minimize the total error between the original filtered offset and the two-segment offset model (*E*(*t*_switch_)), defined as:

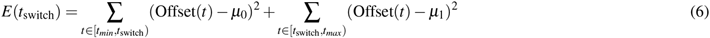

The breakpoint *t*_switch_ is therefore found by minimizing the error:

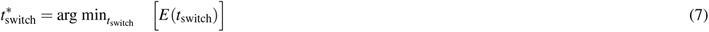

This minimization was performed using the Powell method via the *minimize* function from the Scipy package. Once the optimal breakpoint is identified, the offset is segmented into two parts, each with its respective constant mean offset. In experiments where no offset switching occurs, this method results in two segments with nearly identical offset values, reflecting the stability of the offset throughout the trial. Extended Data Fig. 3d shows the two-segment offset model (dark blue) fitted to the filtered offset (light blue), with the identified breakpoint (vertical red line). By subtracting the two-segment offset model to the original heading of the fly, we obtained the aligned heading direction of the fly (Extended Data Fig. 3e). This aligned heading is what we show in all our figures throughout the manuscript.

To quantify and visualize the magnitude of the offset switch, we calculated the angular shift Δ*µ* as the circular difference between the post-switch mean offset (*µ*_1_) and the pre-switch mean offset (*µ*_0_):

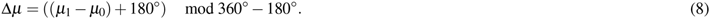

This value (in the range of [*−*180*^◦^,* 180*^◦^*]) represents the sudden phase jump of the PFR or hΔ phase relative to the fly’s heading direction. We visualized these events in the bottom panels of our figures (e.g., Figs. 1e, 2a, 4f, and Extended Data Fig. 4b) by plotting an inverted purple triangle at the time of the switch (*t*_switch_) with the value of the offset jump *µ* (see also Extended Data Fig. 3e). Similarly, this jump is visualized as a purple inverted triangle in all recordings in Supplementary Figs. S1–S3, S7-S11, S19-S21, S22-S25, and S41-S45.

### Correlation between neural phase and heading in the mechanical VR

In some flies, the PFR neurons were unable to reliably track the orientation of the arena. This could be due to various factors, such as the inability of flies to track the low-gain mechanical VR, or variations in fly surgery. To ensure accurate tracking, we discarded flies that did not show good alignment between the PFR phase and the mechanical arena orientation. This was determined by computing the correlation between the PFR phase and the arena orientation during walking.

For this, we first aligned the PFR phase with the mechanical arena orientation using the two-segment offset fit described in the previous section. We then calculated the Pearson correlation coefficient between the arena orientation and the offset PFR phase. Flies with a correlation coefficient lower than a minimum threshold were excluded. The specific threshold values for each experiment are listed in Table S1. While flies walking exclusively forward exhibit higher correlation, flies that walk backward or sideways introduce phase jumps of 180*^◦^* or 90*^◦^*, respectively^45,46^. These movements reduce the overall correlation. We therefore set a threshold of 0.1 (or 0.2 for rest-stimulation and learning experiments) to retain data from flies with diverse walking patterns that do not strictly align their heading with the arena orientation.

### Neural activity recording during mechanical VR navigation and fly selection criteria

For experiments in Fig. 1, Extended Data Fig. 4, and Fig. 4, flies were allowed to navigate in the mechanical VR for 3 trials lasting 5 minutes. At the start of each trial, the fly’s position in the arena was reset to the center.

Because we aimed to quantify neural activity during both walking and resting epochs in PFR and hΔ neurons, we selected trials in which flies exhibited a total resting and walking duration of at least 60 seconds within each 5-minute trial to ensure sufficient data in each behavioral state. After applying the offset correction between the PFR or hΔ phase and fly heading (see Extended Data Fig. 3), we computed the correlation between the PFR or hΔ phase during walking and fly heading (see previous sections). Only flies with a correlation > 0.1 were included. Additionally, to compare the shape of the PFR or hΔ phase distributions during walking with those during rest (Fig. 1j–o; Extended Data Fig. 4g–l, and Fig. 4k–p; see next sections for this analysis), we restricted the analysis to trials where the circular standard deviation of the phase during rest was < 120*^◦^*. Applying these selection criteria to the 20 flies recorded driving expression in PFR neurons with the SS02270 split line (3 trials each) yielded *N* = 13 flies and *N* = 23 trials (Fig. 1j). The same criteria were applied to experiments using the R27G06-GAL4 line (Extended Data Fig. 4), giving *N* = 16 flies and *N* = 27 trials out of 18 recorded flies, and to hΔ recordings (Fig. 4), giving *N* = 12 flies and *N* = 25 trials out of 16 recorded flies. All selection criteria are summarized in Table S1.

Low correlation between PFR phase and heading in the arena was likely due to several technical limitations. First, our mechanical VR has a low rotational gain (0.3). While effective for the majority of the cohort, this resulted in suboptimal heading tracking for some flies, consistent with previous studies where orientation tracking in heading neurons was impaired at lower rotational gains^30^. Second, the delay between the arena response and ball is larger than in projector-based systems. Third, two-photon imaging can lead to photobleaching or toxicity especially during long uninterrupted recordings. Together, these challenges necessitate selection of recordings with consistent heading tracking in the mechanical VR.

### PFR activity profile during walking and rest

We expressed GCaMP8m in PFR neurons using the split line SS02270 (SS02270 > UAS-GCaMP8m) and recorded calcium activity while flies navigated in the mechanical VR for three trials. Only trials that met the selection criteria described above were included (see previous section and Table S1).

PFR neurons show spontaneous activity fluctuations during periods when the fly is immobile or grooming (defined as rest; Fig. 1g). Because the PFR activity profile during walking exhibits a bump-like pattern that shifts across the FB columns, we asked whether a similar bump-like activity profile also exists when the fly is at rest.

To test this, we combined PFR activity from all 13 flies (23 trials) across the 16 ROIs spanning the FB. For each time point, we *z*-scored the activity across the 16 ROIs by subtracting the mean and dividing by the standard deviation, which isolates the shape of the activity profile at each time point. We then projected the 16-dimensional data into two dimensions using principal component analysis (PCA). The resulting points were colored according to the fly’s behavioral state: walking (Supplementary Fig. S5a, black) or rest (Supplementary Fig. S5a, green). In both states, the data formed a ring-like structure in PCA space, reflecting the fact that each time point was normalized independently. The ring geometry allowed us to define a circular trajectory through PCA space (Supplementary Fig. S5a, red line). We then mapped this circular trajectory back to the original *z*-scored (normalized) fluorescence profiles across the 16 FB ROIs. The trajectory contained 500 points for both walking (Supplementary Fig. S5b, black) and rest (Supplementary Fig. S5b, green). This circular trajectory in PCA space corresponds to a continuous activity bump that shifts across the 16 ROIs in the FB. We also mapped the trajectory back to the raw fluorescence data (without normalization) for both states (Supplementary Fig. S5c), yielding similar results. Finally, we aligned the 500 points along the trajectory by their circular phase so that the peak of the activity profile was centered across all points. This alignment revealed a cosine-like activity profile across the ROIs for both walking (Supplementary Fig. S5d, black) and rest (Supplementary Fig. S5d, green), although the amplitude was lower during rest than during walking.

**Table S1.**
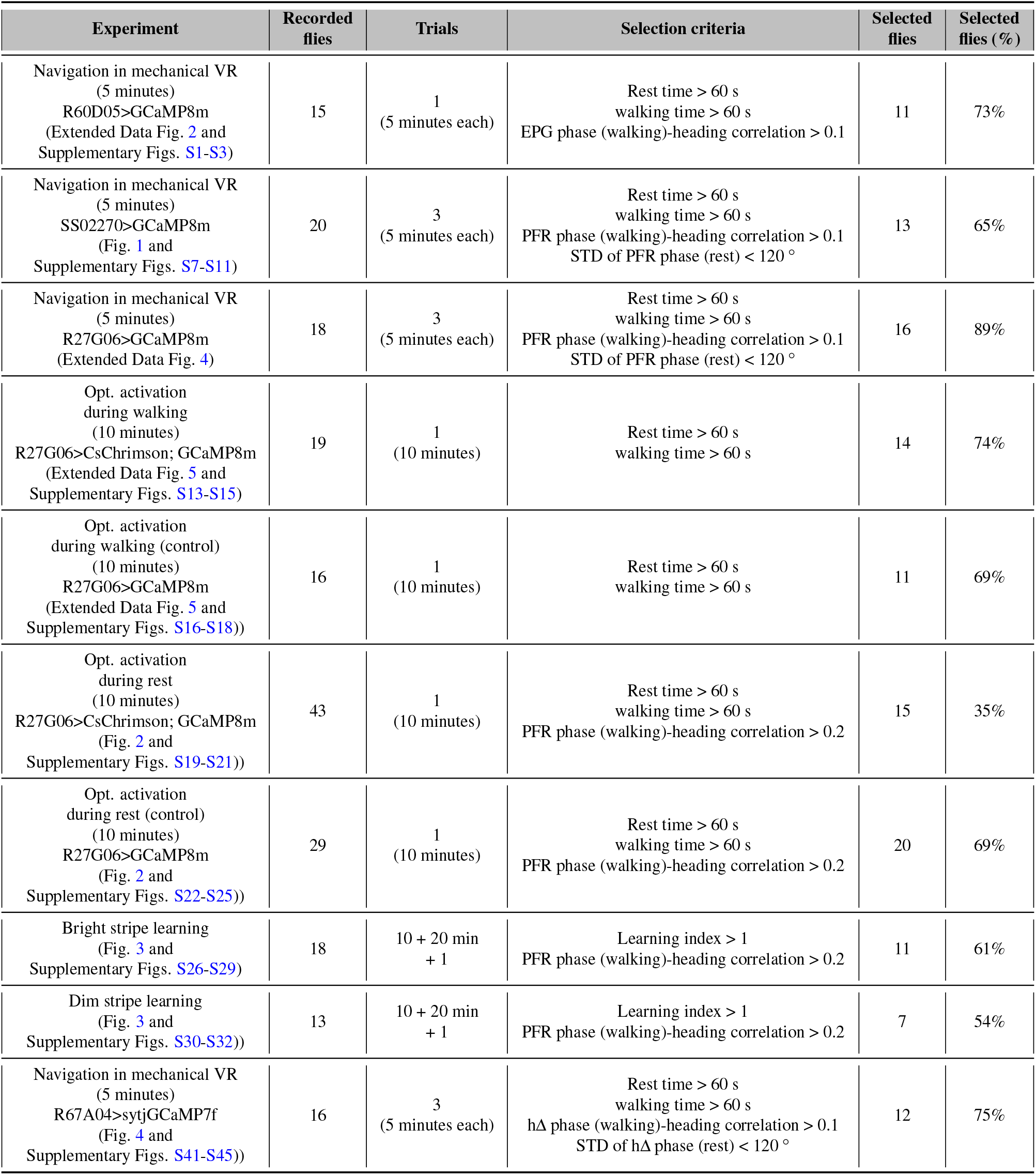
Selection criteria across all experimental groups and number of selected flies.

| Experiment | Recorded flies | Trials | Selection criteria | Selected flies | Selected flies (%) |
|---|---|---|---|---|---|
| Navigation in mechanical VR<br>(5 minutes)<br>R60D05>GCaMP8m<br>(Extended Data Fig. 2 and<br>Supplementary Figs. S1-S3) | 15 | 1<br>(5 minutes each) | Rest time > 60 s<br>walking time > 60 s<br>EPG phase (walking)-heading correlation > 0.1 | 11 | 73% |
| Navigation in mechanical VR<br>(5 minutes)<br>SS02270>GCaMP8m<br>(Fig. 1 and<br>Supplementary Figs. S7-S11) | 20 | 3<br>(5 minutes each) | Rest time > 60 s<br>walking time > 60 s<br>PFR phase (walking)-heading correlation > 0.1<br>STD of PFR phase (rest) < 120 ° | 13 | 65% |
| Navigation in mechanical VR<br>(5 minutes)<br>R27G06>GCaMP8m<br>(Extended Data Fig. 4) | 18 | 3<br>(5 minutes each) | Rest time > 60 s<br>walking time > 60 s<br>PFR phase (walking)-heading correlation > 0.1<br>STD of PFR phase (rest) < 120 ° | 16 | 89% |
| Opt. activation<br>during walking<br>(10 minutes)<br>R27G06>CsChrimson; GCaMP8m<br>(Extended Data Fig. 5 and<br>Supplementary Figs. S13-S15) | 19 | 1<br>(10 minutes) | Rest time > 60 s<br>walking time > 60 s | 14 | 74% |
| Opt. activation<br>during walking (control)<br>(10 minutes)<br>R27G06>GCaMP8m<br>(Extended Data Fig. 5 and<br>Supplementary Figs. S16-S18)) | 16 | 1<br>(10 minutes) | Rest time > 60 s<br>walking time > 60 s | 11 | 69% |
| Opt. activation<br>during rest<br>(10 minutes)<br>R27G06>CsChrimson; GCaMP8m<br>(Fig. 2 and<br>Supplementary Figs. S19-S21)) | 43 | 1<br>(10 minutes) | Rest time > 60 s<br>walking time > 60 s<br>PFR phase (walking)-heading correlation > 0.2 | 15 | 35% |
| Opt. activation<br>during rest (control)<br>(10 minutes)<br>R27G06>GCaMP8m<br>(Fig. 2 and<br>Supplementary Figs. S22-S25)) | 29 | 1<br>(10 minutes) | Rest time > 60 s<br>walking time > 60 s<br>PFR phase (walking)-heading correlation > 0.2 | 20 | 69% |
| Bright stripe learning<br>(Fig. 3 and<br>Supplementary Figs. S26-S29) | 18 | 10 + 20 min<br>+ 1 | Learning index > 1<br>PFR phase (walking)-heading correlation > 0.2 | 11 | 61% |
| Dim stripe learning<br>(Fig. 3 and<br>Supplementary Figs. S30-S32)) | 13 | 10 + 20 min<br>+ 1 | Learning index > 1<br>PFR phase (walking)-heading correlation > 0.2 | 7 | 54% |
| Navigation in mechanical VR<br>(5 minutes)<br>R67A04>sytjGCaMP7f<br>(Fig. 4 and<br>Supplementary Figs. S41-S45)) | 16 | 3<br>(5 minutes each) | Rest time > 60 s<br>walking time > 60 s<br>hΔ phase (walking)-heading correlation > 0.1<br>STD of hΔ phase (rest) < 120 ° | 12 | 75% |

Because the PCA representation also contained points closer to the center of the ring, we repeated the analysis using a circular trajectory with a smaller radius (Supplementary Fig. S5e). Mapping this smaller-radius trajectory back to the normalized (Supplementary Fig. S5f) and non-normalized fluorescence data (Supplementary Fig. S5g) revealed a cosine-like profile (Supplementary Fig. S5h) but with a lower and more diluted amplitude. Together, these results show that the PFR activity profile follows a cosine-like pattern both during rest and walking, with amplitude varying across time.

We repeated the same analysis for recordings where PFR neurons were labeled with the R27G06-GAL4 line, which also exhibits autonomous activity during rest (Extended Data Fig. 4d). The PCA analysis revealed the same cosine-like profile during both walking and rest (Supplementary Fig. S12). This confirms that the activity bump across the FB is present not only during walking but also during rest; for example, the profile does not show multiple bumps, unlike other FB cell types.

### PFR phase velocity during walking and rest

To quantify the PFR drift during rest (Fig. 1g) and to compare it to the PFR phase during walking, we computed the absolute PFR phase displacement for both states. The displacement was calculated as the absolute difference between consecutive PFR phase time points during epochs lasting 5 to 30 seconds in both rest and walking periods. We used the same 13 selected flies as in the previous section (see Table S1 for selection criteria and Supplementary Figs. S7-S11 for all selected flies). Figure 1h shows the combined data from all 13 flies, with PFR phase displacement during rest (green) and walking (black), along with a linear fit (red). Figure 1i presents the slope of the fit (PFR phase velocity) during rest and walking for each fly, with statistical significance assessed via a paired two-sided t-test. On average, the PFR phase velocity during rest was about twice as fast as during walking. Although the mechanical VR had a low rotational gain (0.3), rapid PFR phase shifts were still observed during walking epochs, particularly when flies walked backwards or sideways. These results suggest that the PFR phase velocity during walking is more influenced by the fly’s walking patterns than by the mechanical VR gain (e.g., PFR phase jumps during walking in Fig. 1e and Supplementary Figs. S7-S11).

We also quantified the PFR phase velocity using the GAL4 line R27G06. Using the same selection criteria (see Table S1), we verified in 16 flies that PFR phase velocity during rest was approximately double that during walking (Extended Data Fig. 4e and f). The PFR phase velocity using the GAL4 line R27G06 is slightly faster than in the split line SS02270 (compare Extended Data Fig. 4f and Fig. 1i). This could be due to spontaneous activity from other neural populations labeled in the R27G06 line, since the split line provides cleaner, more specific labeling of the PFR neural population.

### PFR phase distribution during walking and rest

We analyzed the PFR phase distributions during rest and walking for the previously selected 13 flies navigating a mechanical VR for 5-minute trials (selection criteria is shown in Table S1). To enable comparison across flies, we first aligned the heading direction using a two-segment offset model (example pipeline: Extended Data Fig. 3a-e). For each fly, we computed the PFR phase distributions during rest and walking, as well as the heading direction distribution (Supplementary Figs. S7-S11).

To compare these distributions across flies, we identified the peak of the PFR phase distribution during rest (example: Extended Data Fig. 3f) and calculated a “rest-phase offset,” defined as the difference between the peak position and 270*^◦^*. Using this offset, we aligned the rest peak to 270*^◦^* and consistently shifted the walking PFR phase and heading distributions accordingly (example shown in Extended Data Fig. 3g). Following this alignment, the average PFR phase during rest peaked at 270*^◦^* (Fig. 1j), while the PFR phase during walking peaked at approximately 90*^◦^* across all flies (Fig. 1k). A two-sided t-test confirmed a significant difference between the distribution densities at 90*^◦^* and 270*^◦^*. Furthermore, the difference between the circular average of the PFR phase during walking and the circular average during rest was centered around 180*^◦^* across flies, a result that was statistically significant via a two-sided t-test (Fig. 1l), indicating a 180*^◦^* shift in the PFR phase between rest and walking.

After applying the rest-phase offset, the heading direction distribution for all flies also peaked around 90*^◦^* (Fig. 1m), consistent with the PFR phase tracking the heading during walking. The statistical significance of the difference between the 90*^◦^* and 270*^◦^* values was again verified using a two-sided t-test.

We further examined the fly trajectories in the mechanical VR, first aligning them according to the two-segment offset model (see example in Extended Data Fig. 3h). Because this model can assign two distinct rotation angles (*µ*_0_ and *µ*_1_ in Eq. 4), some trajectories exhibit discontinuities. These “jumps” occur because a change in the offset between heading and PFR phase effectively shifts the origin of the fly’s coordinate system. For example, in Extended Data Fig. 3h (top panel), a fly appears to walk first toward the 270*^◦^* direction of the arena and then toward 90*^◦^*. Applying the two-segment model reveals a phase offset jump of approximately 98*^◦^*, which transforms this turn into a consistent downward trajectory toward 270*^◦^* (bottom panel). This suggests that the observed turn was driven by a jump in the fly’s heading representation. This transformation also aligns the arena coordinates with the PFR phase coordinates, ensuring a direct correspondence between arena directions and PFR phase values (e.g., the 0*^◦^* direction in the arena matches the 0*^◦^* PFR phase). Finally, we applied the rest-phase offset to further rotate the trajectories (see example in Extended Data Fig. 3i). These transformed trajectories are shown for individual flies in Supplementary Figs. S7-S11 (right panels) and for all flies in Fig. 1n (left), demonstrating that the transformed trajectories align toward the 90*^◦^* direction of the arena.

To quantify the alignment of transformed trajectories, we computed the “arena azimuth”: the angular density of all trajectories across eight cardinal directions. To exclude initial movement from the center and focus on periods where flies walked sufficiently in a preferred direction, we only included trajectory points located *≥* 50 mm from the arena center (Fig. 1n, right). The individual and average azimuth distributions for all flies are shown in Fig. 1o. A two-sided t-test revealed that these densities at 90*^◦^* was significantly higher from that at 270*^◦^*. These results demonstrate that, when aligned to the rest-phase peak, fly behavior is shifted by approximately 180*^◦^* relative to the PFR phase distribution observed during rest, consistent with the 180*^◦^* offset observed in the PFR phase distribution during walking.

### Optogenetics setup

Optogenetic activation was performed by illuminating specific FB regions with computer-generated holograms projected using a red laser (640 nm, Toptica, iBEAM-SMART-640-S-HP) and a phase-SLM (Meadowlark Optics, HSP1920-1064-HSP8). The setup is described in^52^. A laser beam was expanded and modulated by a phase-SLM before being combined with the fluorescence illumination using a dichroic mirror. The phase-SLM, which generated the holograms, was positioned at a conjugate plane with the microscope’s back focal plane (see Extended Data Fig. 5a). The Gerchberg-Saxton algorithm^82^ was used to generate phase modulation for the desired activation pattern. A custom GUI integrated with Scanimage^83^ allowed users to select activation regions in the two-photon field of view. Calibration for precise alignment between the FOV and the SLM-illuminated area was done using a camera (Basler acA640-750um) imaging reflected and fluorescent light.

### Optogenetic activation during walking

Optogenetic activation was triggered by an Arduino UNO, which controlled a red laser based on real-time ball velocity tracking of the fly. The power of the laser was set to 30*µW* . When the velocity of the fly increased above a defined threshold, determined by baseline noise of the ball tracking algorithm, the laser was turned on after a 0.5-second delay. When the fly rested and the velocity dropped below the threshold, the laser was switched off. During walking periods, the laser alternated between 0.1-second ON and OFF cycles.

We used the GAL4 line R27G06 (GAL4-R27G06 > UAS-CsChrimson; UAS-GCaMP8m), which shows the same 180*^◦^* dynamics between walk and rest periods (Extended Data Fig. 4), since SS02270 was very dim and therefore required high excitation power which could also activate CsChrimson. When the animals were fed retinal-supplemented food, we did not observe robust calcium dynamics, as CsChrimson was strongly activated by the two-photon laser. In the absence of retinal food, calcium activity showed a clear bump, while the red laser was still capable of increasing fluorescence, indicating that red-light activation of CsChrimson can occur without retinal while calcium imaging remains reliable. Therefore, we used flies co-expressing CsChrimson and GCaMP8m without retinal food, while we used flies lacking CsChrimson (GAL4-R27G06 > UAS-GCaMP8m) as controls.

Following recovery from surgery, flies were placed in the mechanical VR and allowed to explore the arena for 5 minutes. Their positions were then reset to the center, followed by a 10-minute trial in which optogenetic activation was applied while the flies navigated. For each fly, a small circular activation mask was defined using the SLM to target a randomly chosen side of the FB (left or right). This side was activated for the duration of the 10-minute trial in both CsChrimson and control flies (Extended Data Fig. 5a). These 10-minute trials of FB activation during walking are shown in Extended Data Fig. 5b and Supplementary Figs. S13–S15 (CsChrimson flies) and S16–S18 (control flies). We compared the rest-to-walk ratio between the first 5 minutes of navigation without laser and the subsequent 10-minute trial with optogenetic laser activation during walking. A paired two-sided t-test revealed no significant effect of laser activation on the rest-to-walk ratio (Extended Data Fig. 6a).

### PFR phase during walking-optogenetic activation

We included flies that walked and rested for at least 60 seconds each, resulting in 14 CsChrimson and 11 control flies (Table S1). For each fly, we computed the PFR phase during rest and walking as previously described. Unlike in previous experiments, we did not filter flies based on the correlation between PFR phase and heading direction. This is because optogenetic activation during walking caused the PFR phase to remain stationary around the activation site, preventing the flies from tracking their heading and occasionally causing CsChrimson flies to turn continuously in the VR (Extended Data Fig. 5b, Supplementary Figs. S13-S15). Because the PFR neurons could not track the heading during activation, we did not quantify walking behavior. Instead, we focused on the dynamics of the PFR phase during activation and the subsequent rest period.

In CsChrimson flies, activating one side of the FB during walking caused the PFR phase during rest to shift 180*^◦^* relative to the activation site (Extended Data Fig. 5c, Supplementary Figs. S13-S15), an effect not seen in control flies (Supplementary Figs. S16-S18). To analyze all flies together, we rotated the PFR phase by 180*^◦^* for flies activated on the right side (at around 270*^◦^*), aligning all activation sites to approximately 90*^◦^*. On average, the PFR distribution during walking in CsChrimson flies was centered at the activation site (90*^◦^*), whereas the distribution in control flies was flat; this difference between 90*^◦^* and 270*^◦^* was statistically significant (Kolmogorov-Smirnov test; Extended Data Fig. 5d). Similarly, the average PFR phase during rest in CsChrimson flies was centered at 270*^◦^* and differed significantly from the flat distribution of control flies (Kolmogorov-Smirnov test; Extended Data Fig. 5e). Additionally, the circular mean of the PFR phase during rest in CsChrimson flies was centered at 270*^◦^* and was significantly different from control flies, which showed a broader distribution across 360*^◦^* (Two-Sample Kuiper test; Extended Data Fig. 5f). These results demonstrate that the PFR phase during rest consistently shifts 180*^◦^* relative to the activation site in CsChrimson flies, but not in controls.

To analyze the PFR phase transition between walking and resting, we extracted traces containing at least 5 seconds of optogenetic activation during walking followed by at least 2 seconds of resting. As before, PFR phases for flies activated on the right side (*∼*270*^◦^*) were rotated by 180*^◦^* to align all activation sites at *∼*90*^◦^*. In Extended Data Fig. 5g, the top panel displays individual PFR traces during walking (black) and resting (green), with the circular mean shown in red. The middle panel shows a 2D histogram of these traces using 16 angle bins and 1-second time bins; bins containing fewer than 10 traces were excluded and appear as white regions. The total number of traces is shown in the bottom panel, which decreases as resting time increases. These data show that upon transitioning to resting, the PFR phase almost immediately shifts to *∼*270*^◦^* and remains there for several seconds. A brief period of approximately 1 second immediately following activation, during which the PFR phase remains at 90*^◦^* despite the fly resting, is likely due to delays in real-time resting-phase detection and latency in turning off the red laser.

### Optogenetic activation during rest

Optogenetic activation during resting was performed using the same setup as described for walking trials. We used flies expressing both GCaMP8m and CsChrimson (GAL4-R27G06 *>* UAS-CsChrimson; UAS-GCaMP8m) without retinal food, with flies expressing only GCaMP8m serving as controls (GAL4-R27G06 *>* UAS-GCaMP8m). After an initial 5-minute exploration period, the flies’ positions were reset to the center of the arena for a 10-minute trial. The red laser power was maintained at 30*µ*W. To trigger activation during resting, the laser was turned on after a 0.5-second delay whenever the fly’s velocity dropped below the threshold defined by the ball-tracking algorithm’s baseline noise. The laser was switched off once the fly resumed walking and exceeded this threshold. During these resting periods, the laser was pulsed in 0.1-second ON/OFF cycles. The activation site for each 10-minute trial was randomly selected (left or right side of the FB), following the same procedure used for walking trials. We also assessed the effect of laser activation during rest periods on the rest-to-walk ratio. Again, no significant effect was observed when comparing the first 5 minutes without laser to the 10-minute trial with laser activation during rest (paired two-sided t-test; Extended Data Fig. 6b).

### PFR phase during rest-optogenetic activation

For the 10-minute optogenetic activation trials, we included flies that exhibited at least 60 seconds of both walking and resting behavior. The PFR phase was computed for both periods as described, and the heading was aligned by calculating the offset via the two-segment model, allowing for at least one offset jump during the trial (Fig. 2a, see also Extended Data Fig. 3). Since the PFR phase during walking was not affected by these experiments, we selected flies capable of reliably tracking their heading direction, imposing a PFR phase (walking)–heading correlation threshold of *≥* 0.2. This is higher than the threshold used in our previous non-optogenetic experiments (set at 0.1) to account for the doubled navigation duration of the trial (10 minutes compared to 5 minutes). Based on these criteria, we included 15 CsChrimson flies and 20 control flies (Table S1). The lower selection percentage (35% of all tested flies) for CsChrimson flies was due to two factors: optogenetic activation during rest occasionally caused CsChrimson flies to resume walking immediately, resulting in total rest periods of less than 60 seconds, and inadvertent CsChrimson activation caused by two-photon imaging likely degraded the PFR phase-heading correlation in many flies. The selection rate differs between experiments mainly because of what each experiment must quantify: the activation-during-rest experiments read out behavior after activation and therefore require reliable phase–heading tracking, whereas the activation-during-walking experiments quantify neural dynamics during subsequent rest and imposed no correlation criterion at all.

To investigate the effects of optogenetic activation during rest, we targeted a section of the FB on the left side (90*^◦^*) or the right side (270*^◦^*). In CsChrimson flies, activation successfully shifted the PFR phase during rest to center around the activation site. Conversely, the PFR phase during walking, as well as the fly’s heading direction, centered approximately 180*^◦^* opposite to the activation site (Fig. 2a and b; Supplementary Figs. S19-S21). No such centering of the PFR phase was observed in control flies (Supplementary Figs. S22-S25).

To allow for a direct comparison across flies, we standardized all data to a 90*^◦^* activation site. For flies activated at 270*^◦^*, we rotated the PFR phase, the heading direction, and the behavioral trajectories by 180*^◦^* to align the activation site to 90*^◦^*.

Following this standardization, the PFR phase distribution during rest in CsChrimson flies was significantly centered at 90*^◦^* (the activation site), whereas control flies showed a flat distribution (Kolmogorov-Smirnov test between 90*^◦^* and 270*^◦^*; Fig. 2d). In contrast, the PFR phase during walking in CsChrimson flies centered at 270*^◦^* (opposite the activation site), a result that was statistically significant (Kolmogorov-Smirnov test between 90*^◦^* and 270*^◦^*; Fig. 2e). This shift was further confirmed by the circular mean of the PFR phase during walking, which differed significantly from the broad distribution seen in control flies (Two-Sample Kuiper test; Fig. 2f).

Consistent with the PFR phase, the heading direction of CsChrimson flies centered at 270*^◦^* (Kolmogorov-Smirnov test between 90*^◦^* and 270*^◦^*; Fig. 2g), with a circular average significantly different from control flies (Two-Sample Kuiper test; Fig. 2h). We observed a similar effect in the spatial trajectories. By transforming the trajectories using the two-segment offset model (see previous sections and Extended Data Fig. 3h) and applying the 180*^◦^* rotation for 270*^◦^* activation flies, the trajectory density (calculated for points *≥* 50 mm from center) peaked at 270*^◦^* in CsChrimson flies, while remaining flat in controls after applying the same trajectory transformation (Kolmogorov-Smirnov test between 90*^◦^* and 270*^◦^*; Fig. 2i). This was further supported by the circular average centered at 270*^◦^* in the arena azimuth position for CsChrimson flies compared to control flies (Two-Sample Kuiper test; Fig. 2j). All transformed trajectories and density distributions are provided in Fig. 2k (CsChrimson) and l (control).

Optogenetic activation during rest did not cause flies to turn toward 270*^◦^* in the physical arena; original 2D trajectories showed a random preference for direction (Supplementary Figs. S19-S21 for CsChrimson flies, and S22-S25 for control flies). However, once transformed by the two-segment offset model, these trajectories aligned consistently in the direction opposite to the activation site. This demonstrates that optogenetic activation in CsChrimson flies consistently biases movement in the direction opposite to the activation site relative to the animal’s internal heading offset.

We further analyzed the PFR phase transition in CsChrimson flies as they transitioned from optogenetic activation during rest to subsequent walking. We extracted trials containing at least 5 seconds of activation during rest followed by at least 2 seconds of walking. As before, PFR phases for flies activated on the right side (*∼*270*^◦^*) were rotated by 180*^◦^* to align all activation sites at *∼*90*^◦^*. As shown in Fig. 2m, the top panel displays individual PFR traces during rest (green) and the subsequent walking period (black), with the circular mean over time indicated in red. The middle panel presents a 2D histogram of these traces (16 angle bins *×* 1-second time bins), where bins containing fewer than 10 traces are omitted (white regions). The bottom panel shows the total number of traces, which decreases as the walking duration increases. These results demonstrate that upon transitioning to walking, the PFR phase shifts to *∼*270*^◦^* and remains stable for several tens of seconds.

### Spatial learning task

We adapted a spatial learning assay from previous studies^28,41^, where flies had to locate a cool area positioned in front of a visual landmark (Fig. 3a). Arena temperature was controlled using an IR laser targeted at the fly’s thorax, which switched off when the fly entered the cool area (Fig. 3b, Extended Data Fig. 1b). Unlike in previous experiments with a single bright stripe, in this task the arena contained two vertical stripes: a self-illuminated “bright” stripe and a “dim” stripe passively lit by the first. Flies were trained in two groups: one with the cool area in front of the bright stripe and another with the cool area in front of the dim stripe (Fig. 3d).

### Aversive stimulus during learning

An infrared (IR) laser (Toptica, ibeam-smart-785-S-HP with pulse option, 785 nm) was used to deliver an aversive heat stimulus, similar to the method described in^41^. The laser beam was focused onto the fly’s thorax using a lens and directed with a small mirror (Thorlabs, MRA05-P01) placed behind the fly holder (Fig. 3b and Extended Data Fig. 1b). Since each fly was attached slightly differently to the slide, a camera was used at the beginning of the experiment to adjust the position of the IR laser after placing the fly on the ball with a kinematic mirror mount. During the learning experiment, the IR laser power was set to 30 mW, and an Arduino UNO was used to pulse the laser at a frequency of 0.2 Hz.

### Analysis of learning behavior

Flies explored the arena freely for 5–20 minutes before training, which consisted of 10 trials. At the start of the first trial, the mechanical VR reset to the starting position (see the last paragraph in this section) a random orientation. An aversive IR laser delivered 0.1-second pulses of “heat” everywhere except the cool area. To prevent prolonged grooming or resting epochs, the air supply to the ball was momentarily turned off for 0.1 seconds if flies rested continuously for over 10 seconds, stimulating them to resume walking. The air valve was triggered by tracking of the velocity of the ball in real time^81^.

Once a fly remained in the cool area for 10 seconds, the next trial began. The bright stripe switched to the opposite side of the arena, and the cool area moved accordingly. This effectively caused an instant 180*^◦^* rotation of the arena from the fly’s perspective, similar to previous place-learning assays in freely moving flies^28^. Each time the fly found the cool area, the stripe and cool area shifted 180*^◦^*, repeating this cycle until all 10 trials were completed. If a fly failed to find the cool area within 15 minutes, a new trial began, and both the cool area and bright stripe were shifted to the opposite side accordingly. In some cases, flies happened to be in the location where the cool area moved, allowing them to immediately “find” it in the next trial by chance. This may have helped some flies navigate back to the cool area in later trials.

We selected flies that met a minimum required PFR phase-orientation correlation of at least 0.2 (see Table S1), as in previous experiments. Additionally, flies that did not demonstrate learning were excluded.

To investigate the correlation of learning behavior with the dynamics of the PFR phase, we selected flies that displayed robust learning behavior. For each recorded fly we calculated a learning index, which quantifies how much faster a fly finds the cool area in later trials (7–10) compared to earlier trials (1–6):

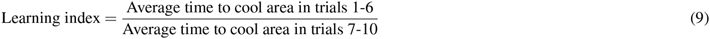

Flies with a learning index below 1, indicating no improvement in performance across trials, were excluded. This criterion also removed flies with a strong initial bias toward the goal location, which would have allowed rapid goal localization in early trials. Based on this selection, 11 of 18 flies were included for the bright stripe condition, and 7 of 13 flies for the dim stripe condition (see Table S1). In our previous methodological study^41^, we did not apply performance-based selection. However, the current experimental setup differs significantly from that study. First, flies underwent surgery for two-photon imaging, which may cause tissue damage. Second, the arena diameter was larger (500 mm vs. 200 mm in^41^). Third, the arena contained two visual cues (bright and dim stripes) instead of one, and the rotational gain was lower (0.3 vs. 0.5). These factors combined to increase the difficulty of the task: flies had to discriminate between two landmarks while undergoing surgery and simultaneous imaging, which can induce phototoxicity and heat stress. Consequently, some recorded flies did not exhibit robust learning. Therefore, we selected only flies showing consistent learning behavior to ensure a reliable correlation between neural activity and learning.

As expected, selected flies in both conditions showed a reduced time to reach the cool area, with a statistically significant difference between the first trial and subsequent trials (two-sided t-test, Fig. 3c and d). Representative trajectories are shown in Fig. 3e. Individual trajectories for all selected flies are provided in Supplementary Figs. S26–S29 (bright stripe condition) and S30-S32 (dim stripe condition).

Selected flies also showed a significant decrease in the walking distance to the cool area across trials (Extended Data Fig. 7a and b; two-sided t-test). Walking velocity remained constant across trials (Extended Data Fig. 7c and d; two-sided t-test), indicating that improved performance was not due to increased locomotion speed. Flies exhibited significantly more resting time in early trials compared to later trials (Extended Data Fig. 7e and f; two-sided t-test). After finishing 10 training trials, the mechanical arena was reset to the starting position, and the flies were given 20 minutes to navigate or rest with the IR laser switched off. Following the 20-minute break, memory was tested in a single trial where the flies had to locate the cool area again. Flies trained with the cool area in front of the bright stripe showed a modest reduction in the time needed to find the cool area on the probe trial compared to the initial trial (Fig. 3c, right side). In contrast, flies that learned to find the cool area in front of the dim stripe did not reliably remember its location, as shown by the lack of significant difference between the times taken in the first trial and the probe trial after the 20-minute break (Fig. 3d, right side).

The starting position in Trial 1 varied across flies: some began at the arena center, while others started opposite the cool area (see Supplementary Figs. S26–S29 and S30–S32). Starting at the center reduces the initial distance to the goal, potentially giving Trial 1 an advantage over subsequent trials, where flies typically start at the opposite side of the arena. To address this, we adjusted the experimental protocol for some flies to begin Trial 1 opposite the cool area, making the initial distance comparable to that of later trials after the stimulus switch. Nevertheless, we included all flies in our analysis as a conservative approach, since starting at the center represents an advantageous baseline (shorter initial distance). Despite this potential bias, we observed statistically significant improvements in performance from trial 1 vs the rest of the trials, confirming that the learning effects are robust and not driven by initial positional advantages.

### PFR phase during learning

We trained flies expressing GCaMP8m in PFR neurons using GAL4-R27G06; UAS-jGCaMP8m, which allowed recording calcium activity during the learning task. To analyze the data, we aligned the fluorescence ROIs and PFR phase to the arena orientation using the two-segment offset model, ensuring that the PFR phase corresponded to the fly’s orientation in the arena. Fig. 3f shows the recorded imaging data from a fly trained with the cool area in front of the bright stripe.

For each fly, we calculated the PFR phase distribution during rest for trials with at least 10 seconds of resting time. Fig. 3g shows the distribution of PFR phase during rest for the fly shown in Fig. 3f across 10 trials, with the cool area located around the 90*^◦^* direction. Individual rest distributions for all flies are shown in Supplementary Figs. S26–S29 (bright stripe condition) and S30–S32 (dim stripe condition). Across flies, we observed that the PFR phase distribution during rest shifted across trials, centering approximately 180*^◦^* opposite to the direction of the cool area.

We grouped the distributions of PFR phase during rest into three sets: trials 1–3, trials 4–7, and trials 8–10 to calculate distributions over extended rest periods. Extended Data Fig. 7g compares the PFR phase distributions during rest for flies trained with the cool area in front of the bright stripe (top) and the dim stripe (bottom). We used PCA to reduce each fly’s distribution to a single dimension and applied a two-sided t-test to assess statistical significance (Extended Data Fig. 7g). Statistically significant differences between bright and dim stripe conditions emerged in the last trials (8-10), with the PFR phase distribution peaking approximately 180*^◦^* from the direction of the cool area.

After the 20-minute break, the distributions of PFR phase during rest remained statistically different (Extended Data Fig. 7g, right), although the distributions were somewhat distorted compared to the last 8–10 trials and the peak of the distributions were no longer centered 180*^◦^* apart from the cool area.

We also calculated the distributions of PFR phase during walking for the three trial sets (Extended Data Fig. 7h). As with the distributions during rest, the PFR phase distributions during walking became statistically significant in the last trials (8-10). After the 20-minute break, these distributions remained significantly different (Fig. 7h on the right).

To quantify the directional alignment of PFR phase distributions, we computed the mean resultant vector for each fly’s activity profile across the grouped trials from above (1–3, 4–7, 8–10, and the post-rest probe trial). For the normalized phase distribution *H_k_*across *N* = 16 bins computed above, we calculated the Cartesian components:

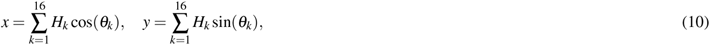

where *θ_k_* is the angle of bin *k*. The resulting vector **v***_d_* = (*x, y*) represents the dominant direction of PFR activity. Individual vectors for each fly are shown in light orange (bright stripe) and gray (dim stripe) in Fig. 3h (rest) and 3i (walking), with group means shown in dark orange and black. In early trials (1–3), vectors were randomly distributed. By trials 8–10, rest-phase vectors shifted to point *∼* 180*^◦^* away from the cool area, while walking-phase vectors aligned toward the cool area. We compared bright– and dim-stripe groups using Hotelling’s *T* ^2^ test. While groups were indistinguishable in early trials, significant differences emerged in trials 8–10 for both rest and walking states (Fig. 3h, i), confirming that PFR dynamics encode learned goal direction.

We also tested whether PFR phase was more strongly aligned during rest than during walking. We computed an alignment score for each fly as the projection of its resultant vector **v***_d_* onto a unit target vector defined by the location of the cool area, **u**^_bright_ for flies trained with the cool area in front of the bright stripe and **u**^_dim_ for flies trained in front of the dim stripe. During walking, the score was *s*_walk_ = **v***_d_ ·* **u**^, and during rest, given the 180*^◦^* offset, *s*_rest_ = *−***v***_d_ ·* **u**^, so that higher scores indicate stronger alignment with the expected direction in each state. For each group of trials (1–3, 4–7, 8–10, and the post-rest probe trial), we compared *s*_rest_ and *s*_walk_ within flies using a one-sided paired Wilcoxon signed-rank test, separately for the bright (Fig. 3j) and dim (Fig. 3k) stripe conditions. We used a one-sided test because our hypothesis was directional (*s*_rest_ *> s*_walk_), and a non-parametric test because of the small number of flies per group. In Fig. 3j,k, violins show the distribution of scores across flies, dots show individual flies (walking in gray, rest in green) connected by gray lines, and the red horizontal lines show the means.

### Connectome and PFR downstream neurons

Using the fly connectome through *neuprint*^78^, we identified hΔA and hΔH neurons as the main targets of PFR neurons (see Table S2), showing a larger number of connections from PFR neurons. The connectome shows that hΔA and hΔH neurons receive input from PFR neurons in two distinct columns, offset by half the width of the fan-shaped body (Fig. 4a), corresponding to a 180*^◦^* phase shift. This is evident in the connectivity matrix between PFR and both hΔA and hΔH neurons (Fig. 4b and Supplementary Fig. S4c), which displays two diagonals: the main diagonal, where no transformation occurs (input from one column produces output in the same column), and an off-diagonal, shifted by 4 columns, representing a 180*^◦^* input-output transformation. By contrast, this does not apply to other columnar neurons, such as PFNd or hΔB neurons, which exclusively connect to hΔA and hΔH neurons along the main diagonal, without any phase shift (Supplementary Figs. S33 and S34). hΔA neurons also project back to PFRa neurons, maintaining their phase without any shift across the FB, as shown by a single diagonal in Supplementary Fig. S4d.

**Table S2.** Mean number of synapses per connection from PFRa to downstream population (FB). Obtained from the hemibrain connectome^78^.

| **PFRa target** | **mean synapses** |
|---|---|
| hΔA | 36.79 |
| hΔH | 27.74 |
| FB4X | 25.97 |
| FB4P_b | 24.33 |
| FB5I | 22.50 |
| hΔB | 21.33 |
| hΔJ | 21.00 |
| FB4Q_b | 20.33 |
| FB4P_a | 20.00 |
| FB4Q_a | 20.00 |
| PFRb | 20.00 |

While some columnar neurons connect to hΔA neurons without causing a phase shift (Supplementary Figs. S33 and S34), several tangential FB neurons establish connections with hΔA and hΔH neurons across FB columns through a 180*^◦^* offset diagonal (Supplementary Figs. S35 and S36). Other tangential neurons and columnar vΔ neurons connect to hΔA and hΔH through both diagonals, similarly as PFR neurons (Supplementary Figs. S37 and S38). This suggests that inputs to hΔA and hΔH neurons can be coordinated, with either the main diagonal (0*^◦^* phase shift) or the off-diagonal (180*^◦^* phase shift) being functionally active depending on the context.

Finally, hΔA and hΔH neurons provide strong input to both PFL2 and PFL3 (Tables S3 and S4) neurons without phase shift (Supplementary Fig. S4g and h, respectively), and these PFL2 and PFL3 neurons drive turning behavior to orient the fly towards a goal^14,15^. Thus, PFR neurons are positioned upstream of the turning PFL neurons, with hΔA and hΔH neurons in between. We therefore hypothesize that the synaptic patterns in hΔA and hΔH neurons could potentially correct the phase shift of PFRa neurons observed during rest.

### Identification of a GAL4 line for hΔ neurons

Given the similarities between hΔA and hΔH in terms of connectivity and upstream and downstream populations, we use hΔ to refer to both populations. To test whether hΔ neurons compensate for the 180*^◦^* shift observed in the PFR phase distribution between walking and rest, we recorded calcium activity from hΔ neuron axons in the FB. Based on the connectome, the correction of the phase distribution is expected to occur at the axonal level. We therefore used the axon-targeted calcium sensor sytjGCaMP7f. To express the sensor in hΔ neurons, we used *neuronbridge*, which matches neuronal morphologies from the connectome to GAL4 expression patterns obtained from confocal imaging^56^. Using this method, we identified the GAL4 line R67A04 as a candidate driver for hΔH neurons, potentially also labeling hΔA neurons (Supplementary Fig. S39). hΔH neurons project across multiple FB layers, which facilitates their identification, as they are easily distinguished from other hΔ neurons (including hΔA) that project within a single FB layer. R67A04 also matches other hΔ types well, such as hΔC (Supplementary Fig. S39a). This is difficult to avoid, because hΔH shares a similar morphology with these other hΔ types, and cleaner split lines would face the same limitation.

**Table S3.** Mean number of synapses per connection from hDeltaA to downstream populations in the FB with at least 20 average synapses. Obtained from the hemibrain connectome^78^.

| **hΔA target** | **Mean synapses** |
|---|---|
| PFL3 | 58.45 |
| PFL2 | 53.00 |
| hΔI | 44.88 |
| PFRa | 43.33 |
| PFRb | 39.27 |
| FB4Z | 35.41 |
| FB5R | 29.33 |
| hΔK | 29.00 |
| FS1A | 28.83 |
| FS1B | 28.29 |
| hΔA | 24.67 |
| FB4P_b | 23.00 |
| FS2 | 20.00 |

### Imaging in hΔ neurons

We recorded axonal calcium activity from hΔ neurons (R67A04 > sytjGCaMP7f) in flies navigating the mechanical VR. Each fly performed three 5-minute trials, and the fly position was reset to the center of the arena at the beginning of each trial, as described for the PFR recordings. Fluorescence signals from hΔ neuron recordings were dimmer and noisier than those obtained from PFR neurons, likely because sytjGCaMP7f is an older calcium sensor and was restricted to axonal compartments. Therefore, after extracting fluorescence signals from the 16 ROIs in the FB, we applied a larger Gaussian filter (5 *×* 1) than the one used for PFR recordings (1 *×* 1), as described above.

### Axonal hΔ activity profile during walking and rest

We applied the same PCA analysis (described in a previous section for PFR neurons) to the axonal calcium activity from hΔ neurons. We again combined the *z*-scored activity from all included trials (25 trials from 12 flies, see Table S1), projected the data into two dimensions using PCA, and defined a circular trajectory through PCA space. Mapping this trajectory back to the 16 FB ROIs revealed the same cosine-like activity profile (Supplementary Fig. S40a–h). Specifically, both during walking and rest, the axonal activity in hΔ neurons exhibited a single, continuous bump that shifted across the FB columns. As with PFR neurons, the amplitude of this cosine-like profile was lower during rest compared to walking, indicating reduced axonal output when the fly is stationary.

### hΔ phase velocity during walking and rest

All included hΔ recordings are shown in Supplementary Figs. S41-S45. As observed for PFR neurons, hΔ activity exhibited a continuous phase drift during rest (Fig. 4h). To quantify this drift, we computed the absolute phase displacement as the absolute difference between consecutive hΔ phase values during epochs of 5-30 seconds, separately for walking and rest periods. Figure 4i shows the combined displacement data from all selected flies during rest (blue) and walking (black), together with a linear fit (red). Figure 4j shows the slope of these fits for each fly, corresponding to the hΔ phase velocity, with significance assessed using a paired two-sided t-test.

**Table S4.**
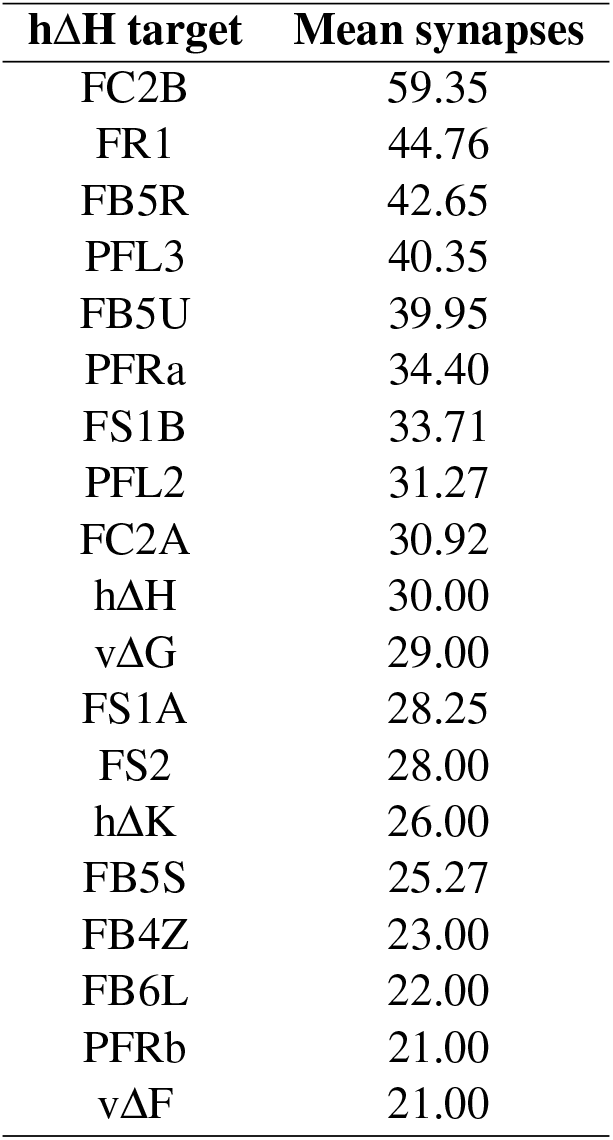
Mean number of synapses per connection from hΔH to downstream populations in the FB with at least 20 average synapses. Obtained from the hemibrain connectome^78^.

On average, the hΔ phase velocity during rest was approximately twice as large as during walking, consistent with the results obtained for PFR neurons (Fig. 1i and Extended Data Fig. 4f). Overall, hΔ phase velocities were slower than those measured in PFR neurons, likely due to the stronger filtering applied to the noisier hΔ fluorescence recordings. Nevertheless, the relative increase in phase velocity during rest compared to walking was preserved, indicating that the drift dynamics of hΔ neurons resemble those observed in PFR neurons.

### hΔ phase distribution during walking and rest

We analyzed hΔ phase distributions during walking and rest using the 12 selected flies described above (selection criteria in Table S1). As for PFR neurons, we first aligned the fly heading using the two-segment offset model described above (example recordings in Supplementary Figs. S41-S45, left). For each fly, we then computed the hΔ phase distributions during rest and walking, together with the corresponding heading direction distributions (Supplementary Figs. S41-S45, middle).

To compare phase distributions across flies, we applied the same analysis used for PFR neurons: we identified the peak of the hΔ phase distribution during rest and computed a rest-phase offset, defined as the angular difference between this peak and 270*^◦^*. This offset was used to align the rest-phase peak to 270*^◦^*, and the same rotation was applied to the walking hΔ phase distribution and heading distribution (see example in Extended Data Fig. 3f-g). Following this alignment, both the rest and walking hΔ phase distributions were centered around 270*^◦^* across flies (Figs. 4k and l). In contrast to PFR neurons, where the phase distributions during rest and walking differed by approximately 180*^◦^*, the circular difference between the average hΔ phase during walking and rest was centered near 0*^◦^* across flies (Fig. 4m). Statistical significance was assessed using a paired two-sided t-test. After applying the same rest-phase offset to the heading distributions, the average heading direction across flies was also centered around 270*^◦^* (Fig. 4n), indicating that hΔ phase activity during walking remained aligned with the heading direction after alignment by the rest-phase distribution.

We next transformed fly trajectories in the mechanical VR using the same alignment procedure applied for PFR neurons. First, trajectories were aligned using the two-segment offset model to compensate for phase-offset switches between heading and hΔ phase. We then applied the rest-phase offset to rotate all trajectories into a common reference frame (see example in Extended Data Fig. 3h-i). Individual transformed trajectories are shown in Supplementary Figs. S41-S45 on the right, and the combined trajectories from all flies are shown in Fig. 4o, left. After alignment, trajectories were predominantly oriented toward the 270*^◦^* direction of the arena.

To quantify this directional bias, we computed the arena azimuth density as described for PFR neurons by measuring the angular density of trajectories across eight cardinal directions. Only trajectory points located at least 50 mm from the arena center were included in the analysis to exclude initial movements near the center of the arena (Fig. 4o, right). Individual and average azimuth distributions are shown in Fig. 4p. In contrast to PFR neurons, where transformed trajectories aligned opposite to the rest-phase peak, hΔ trajectories remained aligned with the rest-phase distribution. These results indicate that hΔ phase activity exhibits similar angular distributions during rest and walking, consistent with a correction of the 180*^◦^* phase shift observed in PFR neurons.

## Data availability

All of the behavior data, raw functional imaging data, and associated behavior data can be made available upon request. The hemibrain dataset is available at https://www.janelia.org/project-team/flyem/hemibrain

## Code availability

All of the code for data processing, analysis and figure generation can be made available upon request.

## Supplementary Information

### FB layers respond to change in temperature

Computational models predict that such learning could occur through plasticity between FB layers and FB columnar neurons^4,16,54^. Based on the anatomical arrangement of CX, computational models typically assume that a reinforc-ing signal should be present in layers of the fan-shaped body. FB neurons respond to taste^84^ feeding^85^, odors^54^, and hunger^52^, and could support the selection of directions associated with these behavioral needs after learning^4,16,54^. We therefore asked whether similarly temperature would be encoded in FB tangential layers.

We found that dorsal tangential neurons respond to temperature changes. These neurons were labeled using the GAL4-72G07 and expressed GCaMP8m (GAL4-72G07; UAS-jGCaMP8m, Extended Data Fig. 8a, left). Using NeuronBridge^56^, we confirmed that they innervate layers 6 and 7 of the FB. We defined 16 regions of interest (ROIs) across the FB, similar to the analysis of PFR neurons (Extended Data Fig. 8a, right). Flies were then trained in the learning task as described in Methods. However, their performance was generally worse than that observed in flies during PFR neuron recordings and we did not analyze their learning behavior in relation to neural activity changes. However, in all flies activity increased whenever they entered the cool area. Extended Data Fig. 8b displays the average fluorescence of 468 traces recorded from defined FB ROIs in 21 flies over 10 training trials, aligned to the moment flies enter the cool area. If flies entered the cool area but stayed for less than 10 seconds during training, the event was cut off at the point they exited the cool area. If flies remained in the cool area for 10 seconds, a new trial began, during which the cool area was switched. Therefore, the maximum time spent by flies in the cool area was limited to 10 seconds. The increase in activity was observed across all FB columns, confirming that the signal spanned the entire FB and corresponded to tangential FB neurons. To further investigate this, we calculated the total fluorescence signal in the FB for each fly, mapping activity across the arena in 15 mm grid bins and averaging across all flies. Extended Data Fig. 8c (left and center panels) shows that the increased activity precisely aligned with the cool area in the arena. To ensure this activity was specific to the cool area, we recorded an additional seven flies navigating the arena for one hour without the IR laser, that is, in the absence of a cool area. In this case, activity remained flat across the arena (Extended Data Fig. 8c, right panel). This confirms that the cool area, that is, the change in temperature, triggered activity in the dorsal FB layers. When fluorescence signals were aligned to the time flies entered the cool area (468 events from 21 flies) and compared to control flies that entered the same location without the IR laser (78 events from 7 flies), we observed a statistically significant increase in fluorescence in the experimental group (Extended Data Fig. 8d, two-sided t-test). Again, if flies entered the cool area but remained for less than 10 seconds, the event was truncated to the time when they exited the cool area.

Additionally, we found that these neurons exhibited spontaneous spiking activity during rest epochs (Extended Data Fig. 8g). Using a threshold to identify spikes, we observed that these events were sparse, typically ranging from 2 to 10 spikes per minute (Extended Data Fig. 8h).

These findings demonstrate that a temperature dependent signal which could serve for reinforcement during the spatial learning task is indeed transmitted to the FB. While these dorsal tangential neurons innervate the upper FB layers, PFR neurons primarily target the central FB, specifically layers 4 and 5. This suggests that direct connectivity between these tangential FB neurons and PFR neurons is unlikely and may rely on intermediate neural populations.

### PFR neurons and connectome analysis

PFR neurons are columnar neurons that project to the PB, FB, and ROB (round body), as illustrated in Fig. 1c. Based on the connectome, there are two types of PFR neurons: PFRa and PFRb^17^. PFRa neurons remain in the same column across the PB and FB, while PFRb neurons shift by one column between these regions. This suggests that PFRb neurons might help shift neural activity in the FB, though their exact function remains unclear. We classify the neurons investigated here as PFRa based on matching analysis using NeuronBridge^56^ (not shown). According to the connectome, PFRa neurons are classified as dopaminergic from electron microscopy images^60^.

Both PFRa and PFRb neurons receive strong input from hΔB neurons, which encode the fly’s traveling direction in allocentric coordinates^45,46^. In fact, PFR activity has been used as a proxy for recording the output of hΔB neurons^46^.

Supplementary Fig. S4a shows the connectivity between hΔB neurons and PFRa neurons, obtained from the hemibrain connectome^17^. In the top-left corner, we display the synapse locations in the FB, with each color representing a distinct presynaptic neuron from the hΔB population. These neurons are sorted based on their output arborization across the FB columns (from left to right). In the bottom-left corner, we show the same synapse locations, but now color-coded according to the postsynaptic neurons, sorted by their output (axonal) arborization in the FB columns. On the right side, we present the connectivity matrix for pre– and postsynaptic neurons in the FB columns, with the number of synapses indicated. As shown in this figure, the hΔB phase is not expected to shift when it is passed to PFRa neurons, with each column connecting to the same corresponding column.

The key outputs of PFRa neurons are to hΔA and hΔH neurons, which are other type of columnar neuron in the FB. The connectome shows that hΔA neurons receive input from PFR neurons in two distinct columns, offset by half the width of the fan-shaped body, corresponding to a 180*^◦^* phase shift. This is evident in the connectivity matrix between PFR and hΔA neurons, which displays two diagonals: the main diagonal, where no transformation occurs (input from one column produces output in the same column), and an off-diagonal, shifted by 4 columns, representing a 180*^◦^* input-output transformation (Fig. 4c). This does not apply to hΔB neurons, which exclusively connect to hΔA neurons along the main diagonal, without any phase shift (Supplementary Fig. S4b). hΔA neurons also project back to PFRa neurons, maintaining their phase without any shift across the FB, as shown by a single diagonal in Supplementary Fig. S4d.

While some neurons, such as hΔB neurons, connect to hΔA neurons without causing a phase shift (Supplementary Fig. S4b), certain tangential FB neurons establish connections with hΔA neurons across FB columns through a 180*^◦^* offset diagonal (Supplementary Fig. S4e and f). This suggests that inputs to hΔA neurons can be coordinated, with either the main diagonal (0*^◦^* phase shift) or the off-diagonal (180*^◦^* phase shift) being functionally active depending on the context (see next section).

Finally, hΔA neurons provide strong input to both PFL2 and PFL3 neurons without phase shift (Supplementary Fig. S4g and h, respectively), and these PFL2 and PFL3 neurons drive turning behavior to orient the fly towards a goal^14,15^. Thus, PFRa neurons are positioned upstream of the turning PFL neurons, with hΔA neurons in between.

### Mechanism for correcting the phase shift of PFR neurons during rest

The 180*^◦^* shift in the PFR phase distribution during rest could be corrected by FB circuits, restoring the original phase distribution observed during walking. This correction would allow the reactivation of the FB columns active during walking to occur during rest, potentially supporting memory consolidation. Such a mechanism is consistent with the connectivity between PFR and hΔA neurons, which includes two diagonals (Fig. 4c), and the way hΔA neurons receive input from both the main diagonal (e.g., hΔB neurons, Supplementary Fig. S4b) and the off-diagonal (e.g., FB4Y and FB5X tangential neurons, Supplementary Fig. S4e and f).

During walking, PFRa and hΔB neurons could provide stronger input to hΔA neurons through the main diagonal, resulting in no phase shift. However, during rest, if hΔB neurons become inactive and tangential neurons are active alongside PFRa neurons, the off-diagonal will receive stronger input than the main diagonal, inducing a 180*^◦^* phase shift between PFRa and hΔA neurons. This shift allows for the correction of the PFRa phase during rest.

### Anatomically constrained model of bump transmission between PFR and hΔ neurons

The synaptic sign of PFR neurons has not been tested experimentally. We therefore asked which combinations of PFR sign (excitatory or inhibitory) and PFR*→*hΔ connectivity can reproduce the hΔ bump during walking and rest. This is a static, qualitative analysis meant to constrain the circuit, not to fit the data.

We modeled *N* = 8 fan-shaped body columns with binary connectivity matrices in a three-population circuit (hΔB, PFR, hΔ; Supplementary Fig. S46a). In FlyWire, hΔB and hΔ neurons (hΔA and hΔH) are predicted to be cholinergic, and thus excitatory^60^. The connections hΔB*→*hΔ and hΔB*→*PFR were set to the main diagonal (Supplementary Fig. S46b). PFR*→*hΔ was tested as main diagonal, off-diagonal (shifted by 4 columns, i.e. 180*^◦^*), or both, with excitatory (Supplementary Fig. S46c) or inhibitory (Supplementary Fig. S46d) PFR. During walking, hΔB activity was a cosine in the range of [0, 1] peaking at column 2 (90*^◦^*) and was transmitted to PFR through the main diagonal. During rest, hΔB activity was zero and PFR activity was a cosine peaking at column 6 (270*^◦^*), reflecting the observed 180*^◦^* offset. The hΔ activity *a* was computed as

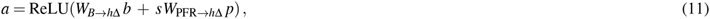

where *b* and *p* are the hΔB and PFR activity profiles, and *s* = +1 (excitatory PFR) or *s* = *−*1 (inhibitory PFR).

The model reproduced the observed hΔ bump (same phase in walking and rest) only with excitatory PFR that transmit the bump through the main diagonal during walking and through the off-diagonal during rest (highlighted in Supplementary Fig. S46c). With inhibitory PFR, hΔ activity during rest was zero after rectification for all configurations (Supplementary Fig. S46d). Inhibitory PFR could also work if the bump were transmitted through the off-diagonal during walking and the main diagonal during rest, but only with an additional constant excitatory input to hΔ across columns at rest, which the inhibitory cosine would then shape into a bump. This requires extra populations or mechanisms not included here. For either sign, transmitting the PFR bump through both diagonals gives PFR inputs that sum to a constant across columns, so PFR carries no phase information to hΔ (Supplementary Fig. S46c,d, third column). Thus, gating between the main and off-diagonal pathways across walking and rest is required regardless of PFR sign.

This analysis supports the switching mechanism and excitatory PFR neurons, but it is not conclusive. The model is a feedforward, static, rate-based simplification of three populations, and other cell types in the FB circuit (e.g. other hΔ types, columnar or tangential neurons) may also contribute to the observed activity.

The connectome also contains a main-diagonal projection from hΔ back to PFR, which we did not include in this analysis. This closes a recurrent loop (hΔ *→*PFR*→*hΔ), and we speculate on its effects without having simulated it. If PFR is excitatory, the loop is a same-phase positive-feedback loop during walking, which could stabilize the bump and support ring-attractor dynamics. During rest, PFR transmit through the off-diagonal, so hΔ would excite PFR 180*^◦^* away from the current PFR bump, which could contribute to the drift observed at rest. If PFR is inhibitory, the same logic applies. During walking (off-diagonal), PFR inhibit hΔ 180*^◦^* away from the bump, while hΔ excite PFR at the bump location, stabilizing it. During rest (main diagonal, with the constant excitatory input), hΔ would again excite PFR 180*^◦^* away from the current PFR bump, which could also contribute to the drift.

**Table S5.**
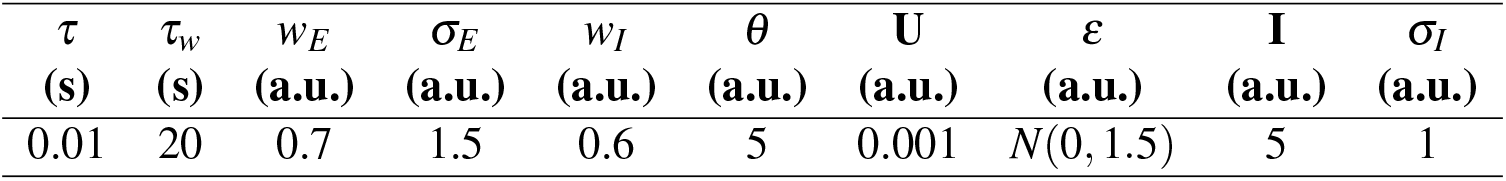
Parameter values used in the simulation of the ring attractor in Fig. 5b.

| *τ*<br>**(s)** | *τ*<sub>*w*</sub><br>**(s)** | *w*<sub>*E*</sub><br>**(a.u.)** | *σ*<sub>*E*</sub><br>**(a.u.)** | *w*<sub>*I*</sub><br>**(a.u.)** | *θ*<br>**(a.u.)** | **U**<br>**(a.u.)** | *ε*<br>**(a.u.)** | **I**<br>**(a.u.)** | *σ*<sub>*I*</sub><br>**(a.u.)** |
|---|---|---|---|---|---|---|---|---|---|
| 0.01 | 20 | 0.7 | 1.5 | 0.6 | 5 | 0.001 | *N*(0, 1.5) | 5 | 1 |

### Ring attractor model

**Table S6.** PFRa input populations in the PB. Obtained from the hemibrain connectome.

| **PFRa upstream populations (PB)** | **Mean synapses** |
|---|---|
| Δ7 | 22.67 |

PFR neurons project to the PB, where their only identified input is from Δ7 neurons (Table S6). Δ7 neurons have sinusoidal connectivity and are hypothesized to provide the inhibition that EPG neurons require for ring attractor dynamics^17^. We therefore propose that PFR neurons may be part of a ring attractor network as well, as also assumed for other FB columnar neurons^16^. In ring attractor networks, activity drift is typically driven by connectivity heterogeneities or noise, which could explain our observations^57–59^.

We employed a rate-based model to simulate a ring attractor network consisting of *N* = 8 neurons, each representing one of the eight columns in the fan-shaped body (FB)^86^. The dynamics of the network are described by the following equation:

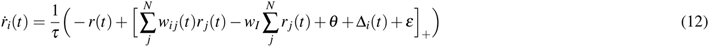

where *r_i_* represents the activity of the *i*-th neuron in the ring attractor, and *τ* is the time constant of the neurons. The recurrent connectivity matrix, *w_i j_*, depends on the distance between neurons *i* and *j* around the ring, and it reflects the interaction between PFR neurons. Alternatively, the reciprocal projection from hΔ neurons back onto PFR neurons (Supplementary Fig. S4) could also contribute to this recurrent connectivity. The term *w_I_* represents global inhibition within the network, which could be potentially mediated by Δ7 neurons; *θ* is a constant background input, Δ*_i_*is the input from hΔB neurons to the *i*-th column, and *ε* represents random noise input sampled from a normal distribution. The term [*·*]_+_ is a threshold-linear function to ensure positive-valued firing rates.

We assume synaptic depression in the *w_i j_* weight matrix, which is described by the following equation:

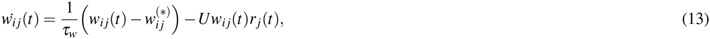

where *τ_w_* represents the time constant of synaptic depression, 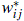 represents the baseline synaptic weight in the absence of activity-dependent depression, *U* is a constant governing the strength of synaptic depression, and *r _j_*(*t*) is the activity of the presynaptic neuron *j*. High presynaptic activity leads to neurotransmitter depletion, which in turn reduces the synaptic strength, making the depression dependent on the presynaptic neuron’s activity.

The baseline synaptic weight, 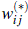, is defined with a Gaussian-like function that depends on the distance between *i* and *j* neurons in the ring:

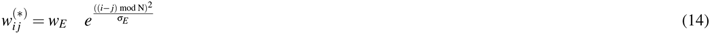

where *w_E_* represents the magnitude of synaptic weights and *σ_E_*determines the width of the synaptic connectivity pattern, influencing how the connectivity strength decreases with distance between neurons in the ring. All parameter values used for the ring attractor simulation are shown in Table S5.

Synaptic depression in the model, along with random input noise, *ε*, induces drift in the ring attractor. When a neuron reaches peak activity, it inhibits other neurons through sustained activity, which leads to synaptic depression. This sustained activity diminishes the active neuron’s influence over time, causing it to lose dominance. Consequently, the peak activity shifts to a different neuron with a non-depressed synaptic weight (i.e., with more available resources). As the depleted neuron becomes fully inhibited and its activity drops to zero, its resources recover, while the newly active neuron weakens, continuing the cycle (Fig. 5a and b, left green side).

In our model, we assume that during walking, the ring attractor network receives strong input from heading direction signals, which are encoded in EPG neurons^30^, and can be conveyed indirectly to PFR neurons through hΔB neurons. We model the heading input with a cosine-like pattern over time, as shown in Fig. 5b (blue), using the following equation:

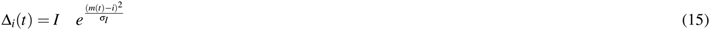

Here, *I* represents the amplitude of the bump, *σ_I_*defines the standard deviation of the activity across the FB columns, and *m*(*t*) represents the sine trajectory across the FB, given by:

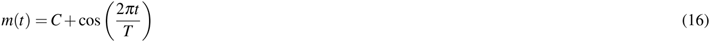

In this equation, *C* is the column along which the sine-like input trajectory oscillates (set to *C* = 4), and *T* is the oscillation period, with *T* = 5 seconds.

This strong input helps the ring attractor counteract drift and accurately track the directional signal (Fig. 5b, black), even in the presence of synaptic depression (Fig. 5b, bottom). We then simulate the fly at rest by setting the input from hΔB neurons to zero. As a result, the peak of activity in the network drifts, governed by the dynamics of synaptic depression (Fig. 5b, bottom). Consequently, the activity shifts toward the neurons that were less active during walking (Fig. 5b, right green side). This results in a 180*^◦^* phase shift in the distribution between rest and walking within the ring attractor network (Fig. 5c). This model suggests that the 180*^◦^* shift in the PFR phase distribution between rest and walking is driven by synaptic depression, while drift is influenced by both random noise and synaptic depression.

### Connectome-based computational model

We developed a biologically inspired computational model to simulate the interaction between neural dynamics in the fan-shaped body (FB) and navigation behavior in a virtual arena. The model consists of three main components: a simulated fly agent navigating a 2D circular arena, a ring attractor network representing PFR neurons with short-term synaptic depression, and a downstream circuit that includes hΔ neurons, tangential neurons, and PFL3 neurons.

### Simulated arena and trajectories

The simulated fly agent navigates a 2D circular arena of radius *R* = 135 mm, as our mechanical VR. The state of the agent at time *t* is defined by its position **p***_t_* = (*x_t_, y_t_*) and heading angle *θ_t_ ∈* [0, 2*π*). The simulated fly only receives the orientation information in the arena discretized in *N* = 8 orientation bins corresponding to the EB and FB columns^30^. The heading angle information that the fly retrieves at a given time

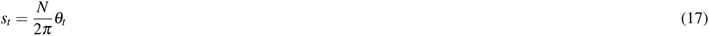

The simulation runs in discrete time steps Δ*t* = 0.001 s. For simplicity we forced the agent to alternate between a walking period of *T_walk_* = 20 seconds, and a rest period of *T_rest_* = 20 seconds. During rest, the agent’s position and heading remain fixed, but neural activity continues to evolve.

During walking, the agent can perform (see below) one of three actions: turn left, turn right, or move forward.

Heading and position are updated as follows:

- Turn left: *θ_t_*_+1_ = *θ_t_ −v_θ_* Δ*t*
- Turn right: *θ_t_*_+1_ = *θ_t_* + *v_θ_* Δ*t*
- Move forward: **p***_t_*_+1_ = **p***_t_* + *v ·* (cos(*θ_t_*), sin(*θ_t_*))Δ*t*

where *v_θ_* = *π/*2 rad/s is the rotation velocity and *v* = 30 mm/s is the forward velocity. If the simulated fly exceeds the arena boundary, its position is projected back onto the edge.

#### PFR neurons

The PFR population is modeled as a ring attractor network with *N* = 8 neurons, as described above. During walking, the network receives the current discretized heading via equation (15), whereas during rest no heading input is provided, allowing the activity bump to drift freely.

#### hΔ neurons

The hΔ neurons receive input from the PFR neurons. To model the experimental finding that hΔ phase distribution remains consistent across behavioral states while PFR activity shifts, we implement a state-dependent connectivity pattern. The hΔ activity *r*^hΔ^ is driven by:

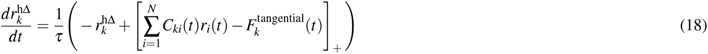

where *τ* is the same time constant as defined for PFR neurons. The connectivity matrix *C_ki_*(*t*) switches between two modes based on the behavioral state:

- **During Walking:** The connectivity preserves the phase, such that

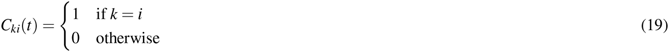 This ensures hΔ tracks the current heading (Fig. 4d).
- **During Rest:** The connectivity shifts by four columns (180*^◦^*), such that

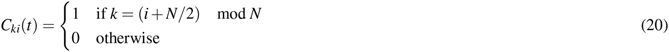 This compensates for the PFR drift, ensuring that the hΔ activity remains aligned with the previous walking direction, effectively correcting the phase shift (Fig. 4e).

This state-dependent routing mimics the structural connectivity of hΔ neurons receiving input from both the main diagonal and the 180*^◦^*-shifted diagonal of the PFR population, as observed in the connectome (Fig. 4c).

We assumed that hΔ neurons encode the fly’s instantaneous goal direction, as reported for other columnar neurons^14^. In our model, goal signals in hΔ are modulated by inhibition from tangential FB neurons, as suggested by other computational models^16^ (Fig. 5f). Each tangential neuron is tuned to a preferred direction (e.g., cool area, food source, or other behavioral needs), and the collective activity of these neurons biases hΔ activity away from the PFR-driven baseline.

For simplicity, tangential input was modeled as a random drive to hΔ neurons. We modeled 8 tangential neurons, each with a distinct preferred direction *m_F_* = 0, 1*, . . .,* 7. At each timestep, a new random tangential neuron was selected uniformly with probability 20% to represent spontaneous switching between different goal directions. Otherwise (80% probability), the previously selected tangential neuron remained active, guiding the fly toward its preferred direction. The selected tangential neuron modulated hΔ activity across columns following a Gaussian profile centered on its preferred direction:

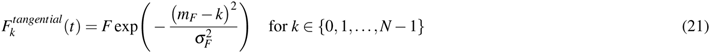

where *F* = 2 is the amplitude and *σ_F_* = 2 is the width of the input profile. Because the preferred direction *m_F_* was chosen randomly, the simulated flies navigated toward random directions over time.

#### Decision Policy and Motor Output

During walking, PFL3 neurons compute the error between the current heading (from EPG) and the goal direction (encoded in hΔ) to generate corrective turns via an offset connectivity pattern (Fig. 5d), as previously described^15^. In our model, the goal direction is modulated by the random tangential input 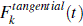 described above.

Steering is computed as the asymmetry in PFL3 activity on the left and right sides of the brain. Specifically, the activity of left PFL3 neurons (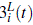) and right PFL3 neurons (PFL3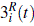) is given by the rectified difference between the heading input and hΔ activity, shifted by two columns (*±*90*^◦^*):

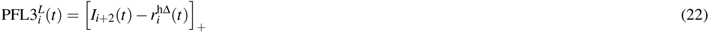

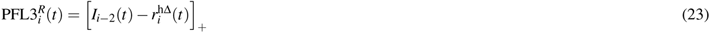

where *I_i±_*_2_(*t*) denotes the heading input shifted by *±*2 columns.

PFL3 neurons project to the lateral accessory lobe (LAL) neuropil, where signals from all PFL3 neurons are summed to generate left turns (D*^L^*(*t*)) or right turns (D*^R^*(*t*))^15,16^ (Fig. 5d):

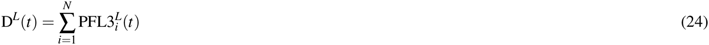

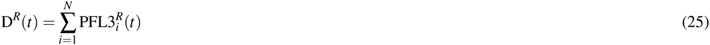

The net steering signal is then:

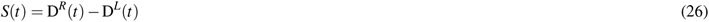

If *S*(*t*) *> θ*_turn_, the fly turns left; if *S*(*t*) *< −θ*_turn_, it turns right; otherwise it moves forward. Here, *θ*_turn_ = 0.1 is the turning activity threshold. This closed-loop coupling ensures that the neural dynamics directly influence navigation behavior.

### Closed-loop navigation simulations

We simulated 20 flies navigating in a circular arena for 300 seconds, starting at the center as in our experiments. As described above, each simulated fly alternated between walking and resting periods of 20 seconds each.

Figure 5g shows the simulated fly orientation (first row) and the corresponding internal heading signal discretized into *N* = 8 columns (second row). During walking, PFR neurons receive input from the heading signal and track it, while during rest—absent this input—PFR activity drifts due to synaptic depression, typically shifting by 180*^◦^* relative to the walking direction. hΔ neurons receive input from PFR neurons and are additionally modulated by tangential FB neurons, generating small deviations from the heading that drive closed-loop navigation. Because the simulation operates in closed loop, hΔ activity closely tracks the heading direction as the fly aligns its goal to its heading. During rest, the state-dependent connectivity shifts PFR signals to the off-diagonal (Fig. 5e), so that the drift in hΔ neurons is no longer shifted relative to the activity during walking. Finally, heading and hΔ signals are integrated in PFL3 neurons during walking to drive navigation (Fig. 5h), as detailed above.

For comparison with our experimental data (Figs. 1, 4, and Extended Data Fig. 4), we applied the same analysis pipeline to the 20 simulated flies. We computed the PFR phase distribution during rest and aligned its peak to 270*^◦^* using the rest-phase offset (Extended Data Fig. 9a). Applying this offset to the PFR phase during walking revealed a peak at 90*^◦^* (Extended Data Fig. 9b), consistent with our experimental observations (Fig. 1). Similarly, the hΔ phase distributions during rest and walking, both shifted by the rest-phase offset, showed aligned peaks at 90*^◦^* (Extended Data Fig. 9c and d), reproducing the alignment observed in our experiments (Fig. 4). Finally, rotating the simulated heading and trajectories by the rest-phase offset resulted in an alignment of heading at 90*^◦^* (Extended Data Fig. 9e) and trajectories toward the 90*^◦^* direction of the arena (Extended Data Fig. 9f and g), as observed in our experiments.

These simulations reproduce our experimental findings, demonstrating that aligning the simulated PFR drift during rest results in aligned activity distributions during walking and corresponding behavioral alignment.

### Simulated optogenetic walk-to-rest transitions

We also simulated the optogenetic activation experiments during walking (Extended Data Fig. 5) to examine the transition from walking to rest in our model. We simulated 20 flies and applied a constant input to a single column corresponding to 90*^◦^* for 10 seconds during walking, simulating targeted optogenetic activation of one PFR column.

This was followed by a rest period during which activity was allowed to drift freely in the ring attractor. We then computed the PFR phase during walking and rest and plotted all simulated flies together (Fig. 5i). The first row shows individual PFR phase traces during walking (black) and rest (green), with the mean PFR phase during rest across flies in red. The second row shows the density of these traces over time (1-second bins) across the 0–360*^◦^* phase range. Our simulated walk-to-rest transition shows that the PFR phase during rest shifts to approximately 180*^◦^* opposite the activation site during walking, centering around 270*^◦^* for roughly 20 seconds before the tuning is lost and the traces drift randomly. This retention of the memory trace for several seconds matches our experimental observations (Extended Data Fig. 5g). The duration of this retention is governed by the time constant of the synaptic depression mechanism, *τ_w_*, in our ring attractor model of PFR neurons.

**Figure S1.**
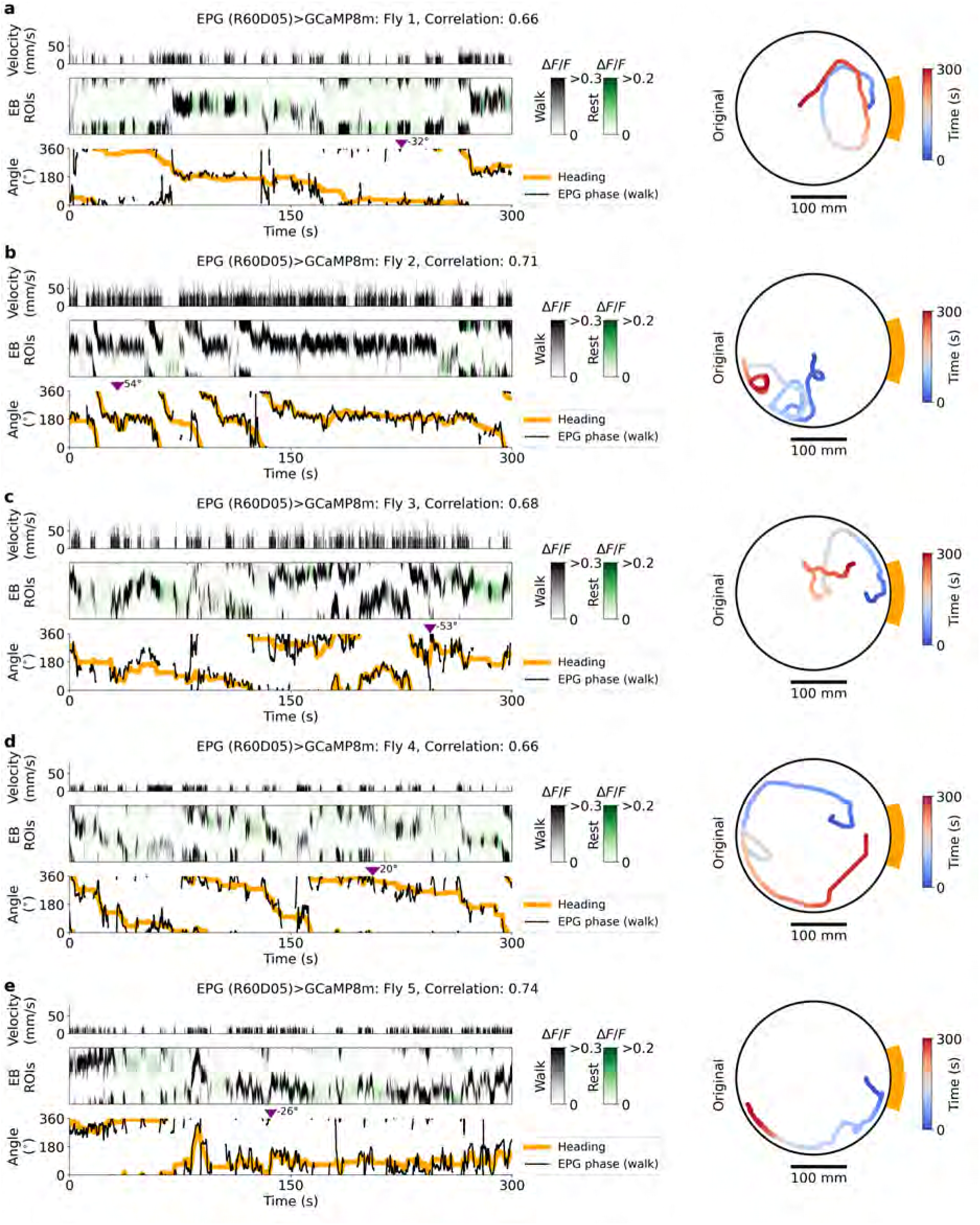
Neural activity and behavior in flies 1-5 expressing GCaMP8m in EPG neurons (GAL4 line R60D05) navigating the mechanical VR. Data shown are from selected flies during 5 min duration of navigation with at least *≥*60 s of walking and resting, and a EPG phase–heading correlation *>* 0.1 (selection criteria in Table S1). **a–e** Left panel: Fly velocity (top); fluorescence in EB ROIs during walking (black) and rest (green, middle); EB phase during walking (black) and fly heading (orange, bottom). Right panel: Original fly trajectory in the mechanical VR.

**Figure S2.**
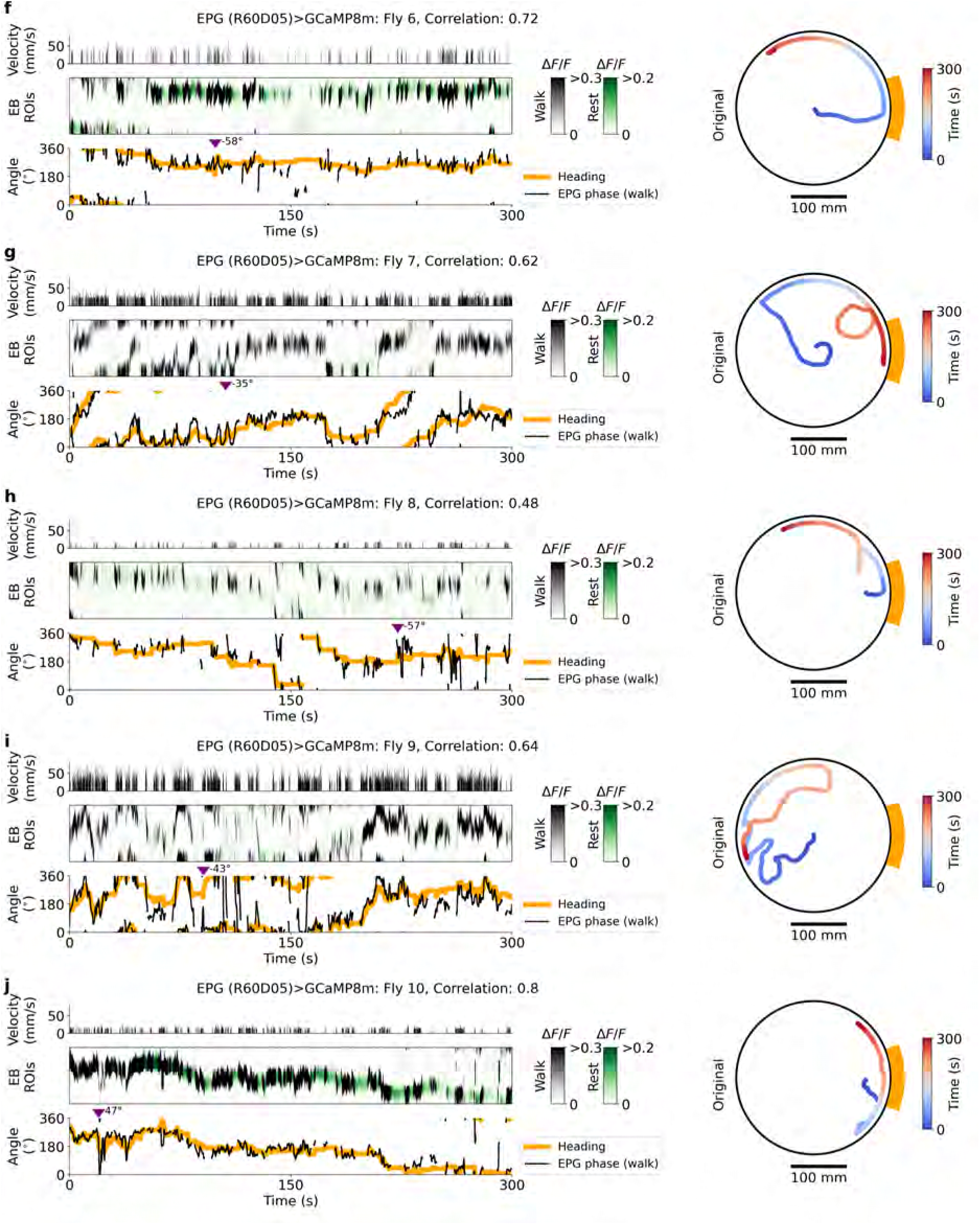
Neural activity and behavior in flies 6-10 expressing GCaMP8m in EPG neurons (GAL4 line 60D05) navigating the mechanical VR. **f-j** Same as Supplementary Fig. S1a-e.

**Figure S3.**
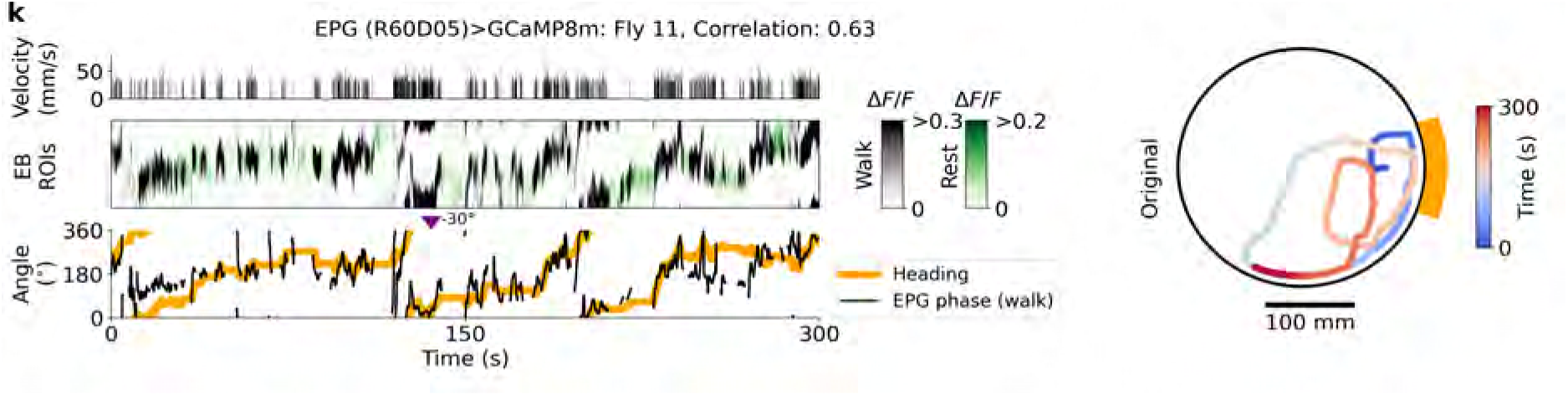
Neural activity and behavior in fly 11 expressing GCaMP8m in EPG neurons (GAL4 line 60D05) navigating the mechanical VR. **k** Same as Supplementary Fig. S1a-e.

**Figure S4.**
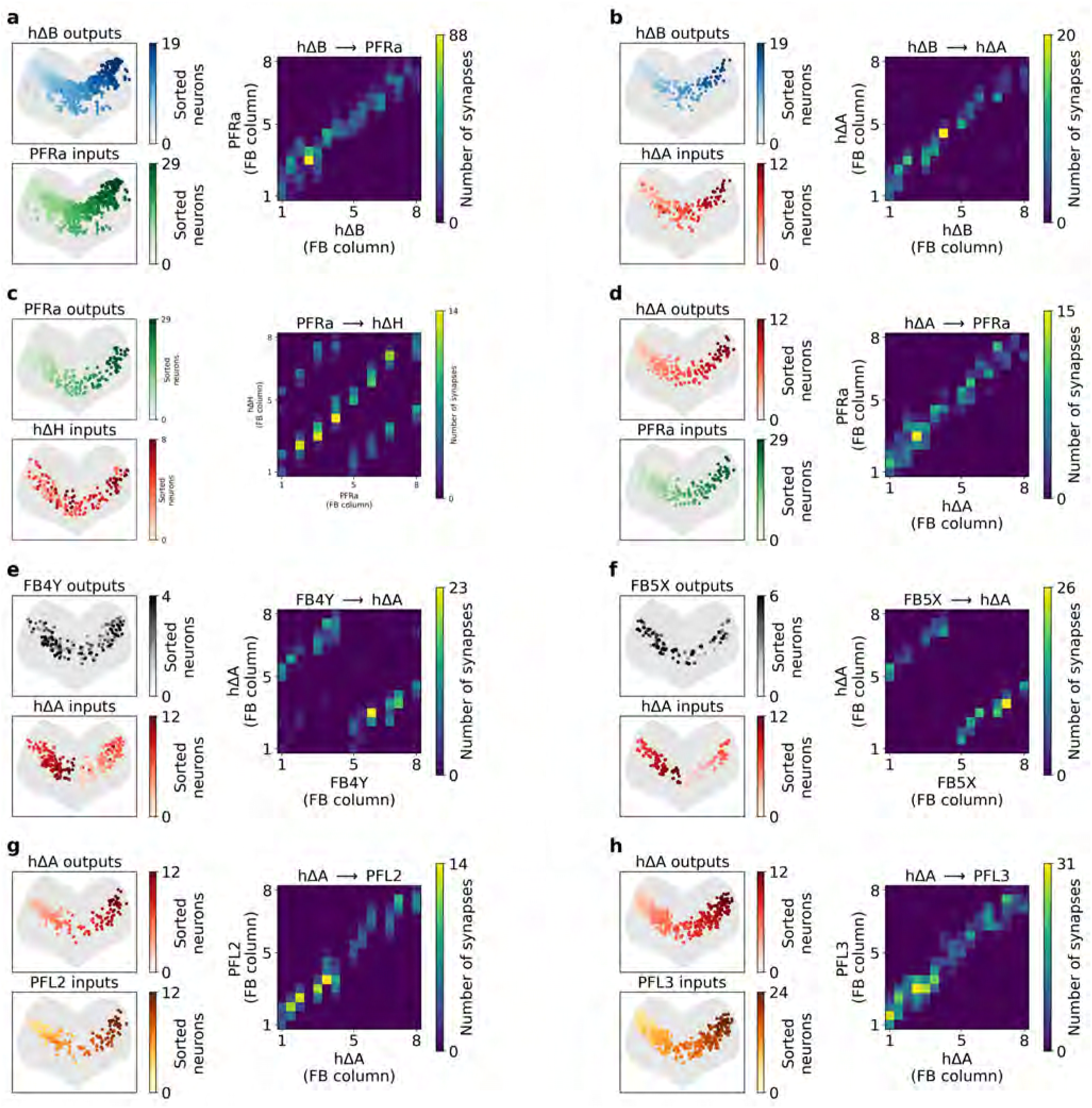
Neural connectivity according to the hemibrain connectome^17,77,78^. **a** Top left: Each point represents a synapse in the FB, with color indicating the presynaptic neuron from hΔB population. Neurons are arranged from left to right based on their axonal (output) arborization in the FB. Bottom right: The same synapses as in the top left panel, now colored according to their corresponding postsynaptic PFRa neurons, which are also sorted by axonal arborization in the FB. Right panel: The number of synapses between hΔB and PFRa neurons across the eight columns of the FB. **b-h** Same as b but for other populations indicated at the top of each panel.

**Figure S5.**
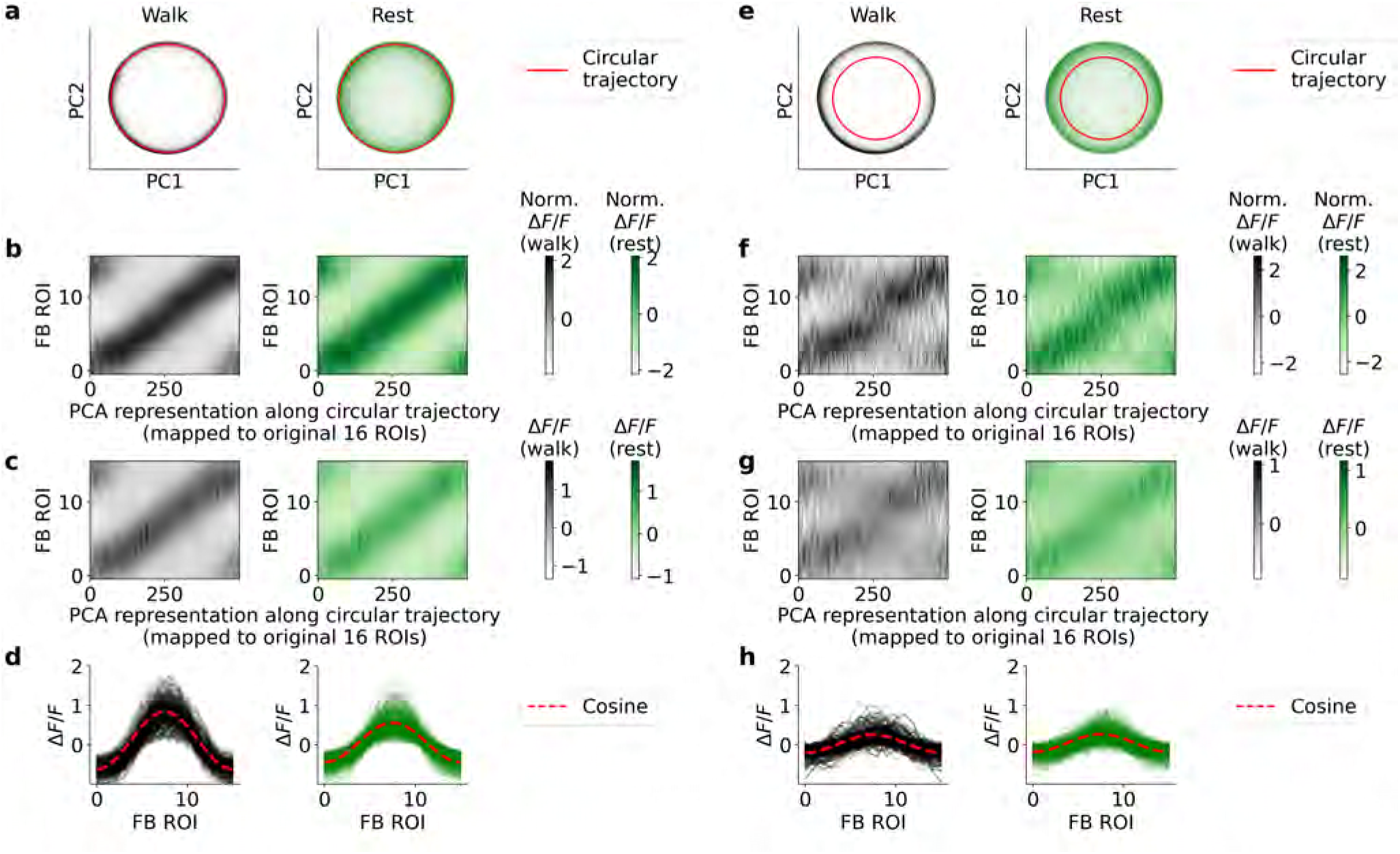
Activity profile of PFR neurons during rest and walking using the split line SS02270. **a** Fluorescence data from all included flies (13 flies total, see Table S1) were combined and normalized (see Methods), resulting in 200,762 fluorescence traces for each of the 16 ROIs in the FB. Principal component analysis (PCA) was applied to reduce the 16-dimensional normalized fluorescence traces (see Methods) to two dimensions, with data points during walking (black, left panel) and resting (green, right panel) displayed along the principal components. **b** A circular trajectory was defined along the reduced data (red line in panel a), with 500 points along the trajectory mapped back to the normalized fluorescence data across the 16 ROIs during walking (black) and rest (green). **c** The same as panel b, but the circular trajectory was mapped to the original (non-normalized) fluorescence data. **d** The 500 data traces from panel c are aligned based on their phase during walking (left, black) and rest (right, green), showing a cosine-like profile (red line) across the ROIs of the FB. **e-h** Same as panels a-d but for a circular trajectory defined in panel e with a smaller radius in the reduced 2D PCA dimensions.

**Figure S6.**
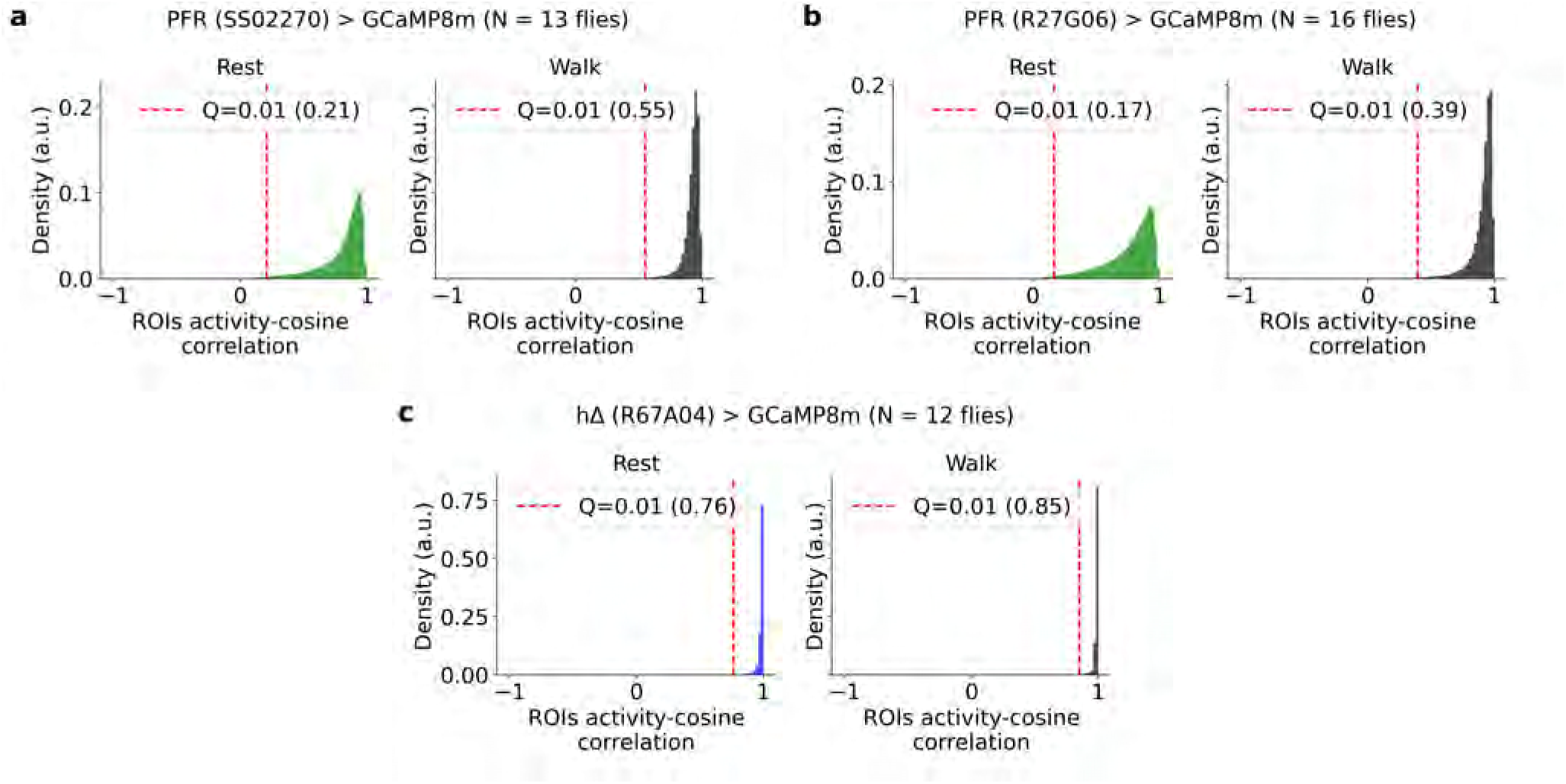
Cosine Pearson correlation of activity profiles. **a** Distribution of the correlation between ROI activity and a cosine function during rest (left) and walking (right) for PFR neurons labeled with SS02270-GAL4. The vertical line indicates the 1% quantile. **b** Same as **a**, but for PFR neurons labeled with R27G06-GAL4. **c** Same as **a**, but for hΔ neurons.

**Figure S7.**
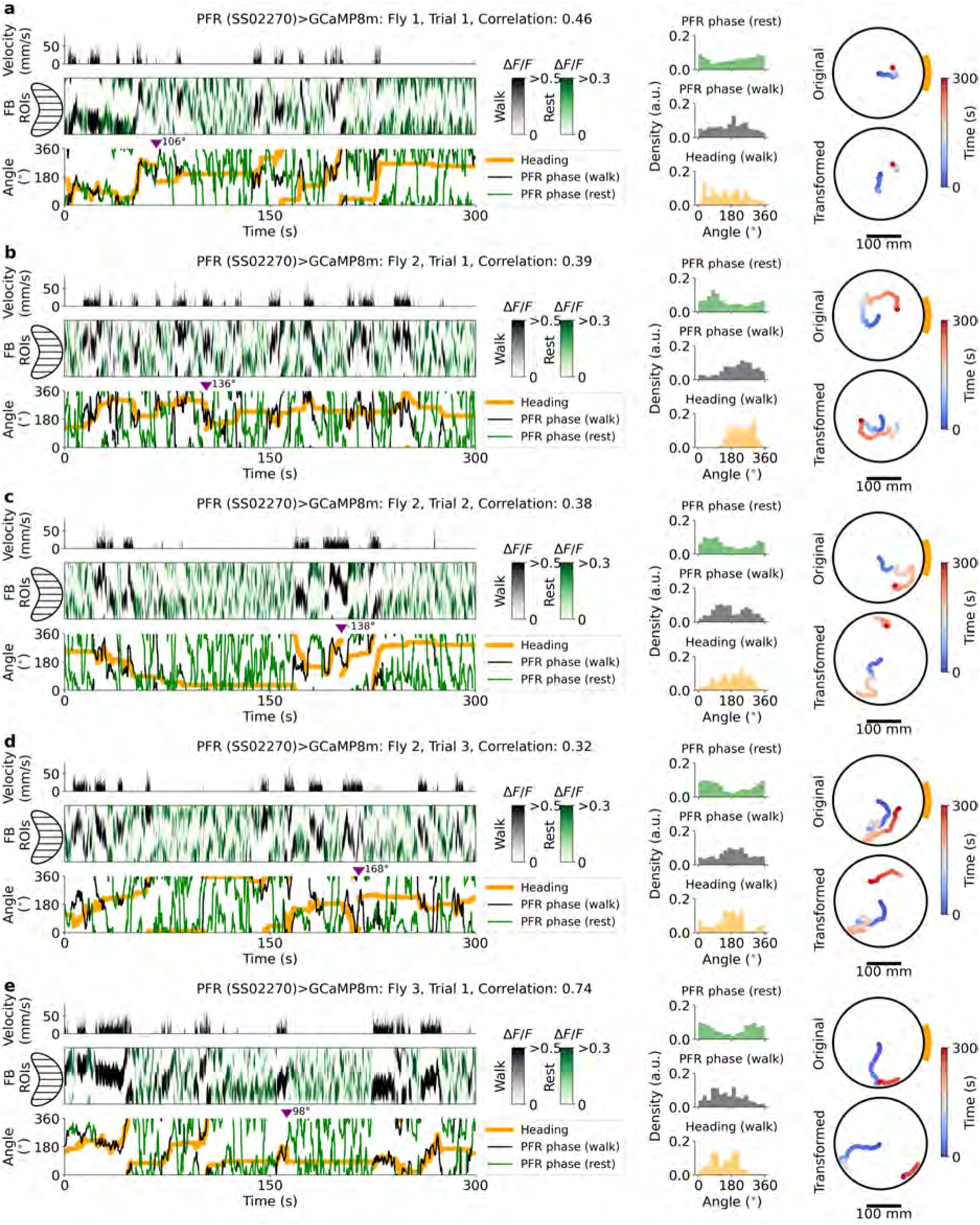
Neural activity and behavior in flies 1-3 expressing GCaMP8m in PFR neurons (split-GAL4 line SS02270) navigating the mechanical VR. Data shown are from selected trials (5 min duration) with *≥*60 s of walking and resting, and a PFR phase–heading correlation *>* 0.1 (selection criteria in Table S1). **a–e** Left panel: Fly velocity (top); fluorescence in FB ROIs during walking (black) and rest (green, middle); PFR phase during walking (black) and rest (green), and fly heading (orange, bottom). Middle panel: PFR phase density during rest (green, top) and walking (black, middle); heading density (orange, bottom). Right panel: Original fly trajectory in the mechanical VR (top); trajectory transformed by the two-segment offset model and rest-phase offset (bottom; see Methods and Extended Data Fig. 3).

**Figure S8.**
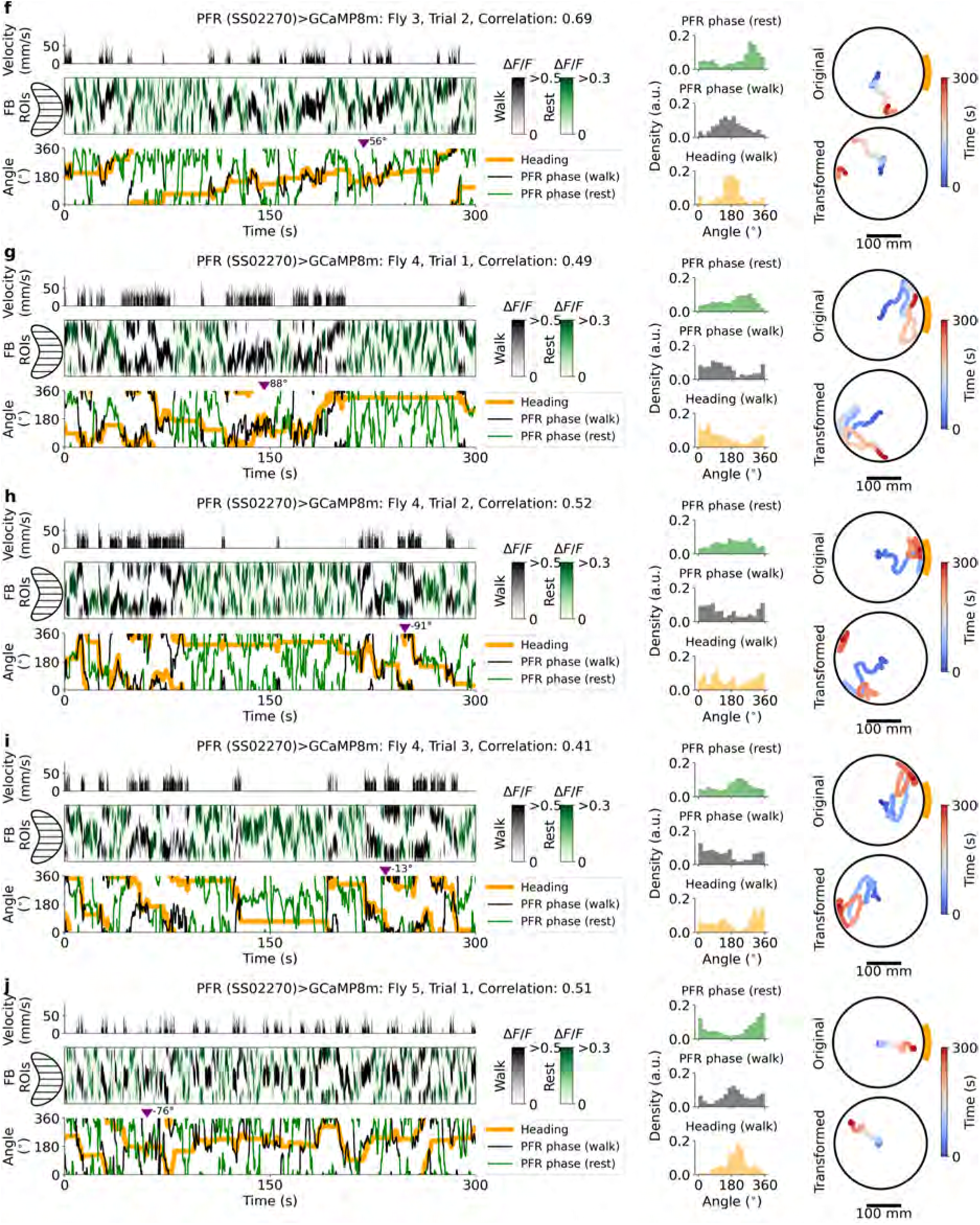
Neural activity and behavior in flies 3-5 expressing GCaMP8m in PFR neurons (split-GAL4 line SS02270) navigating the mechanical VR. **f-j** Same as Supplementary Fig. S7a-e.

**Figure S9.**
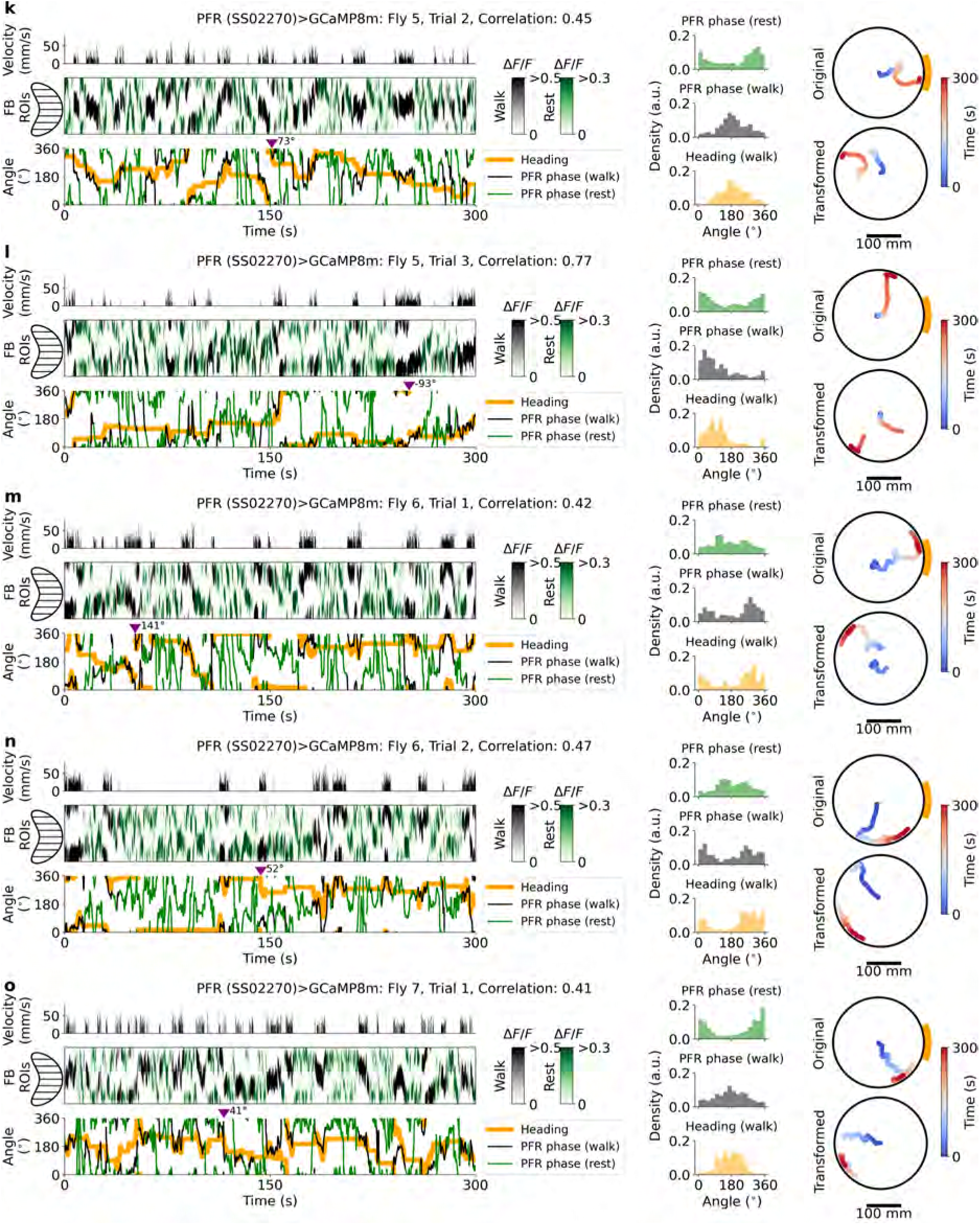
Neural activity and behavior in flies 5-7 expressing GCaMP8m in PFR neurons (split-GAL4 line SS02270) navigating the mechanical VR. **k-o** Same as Supplementary Fig. S7a-e.

**Figure S10.**
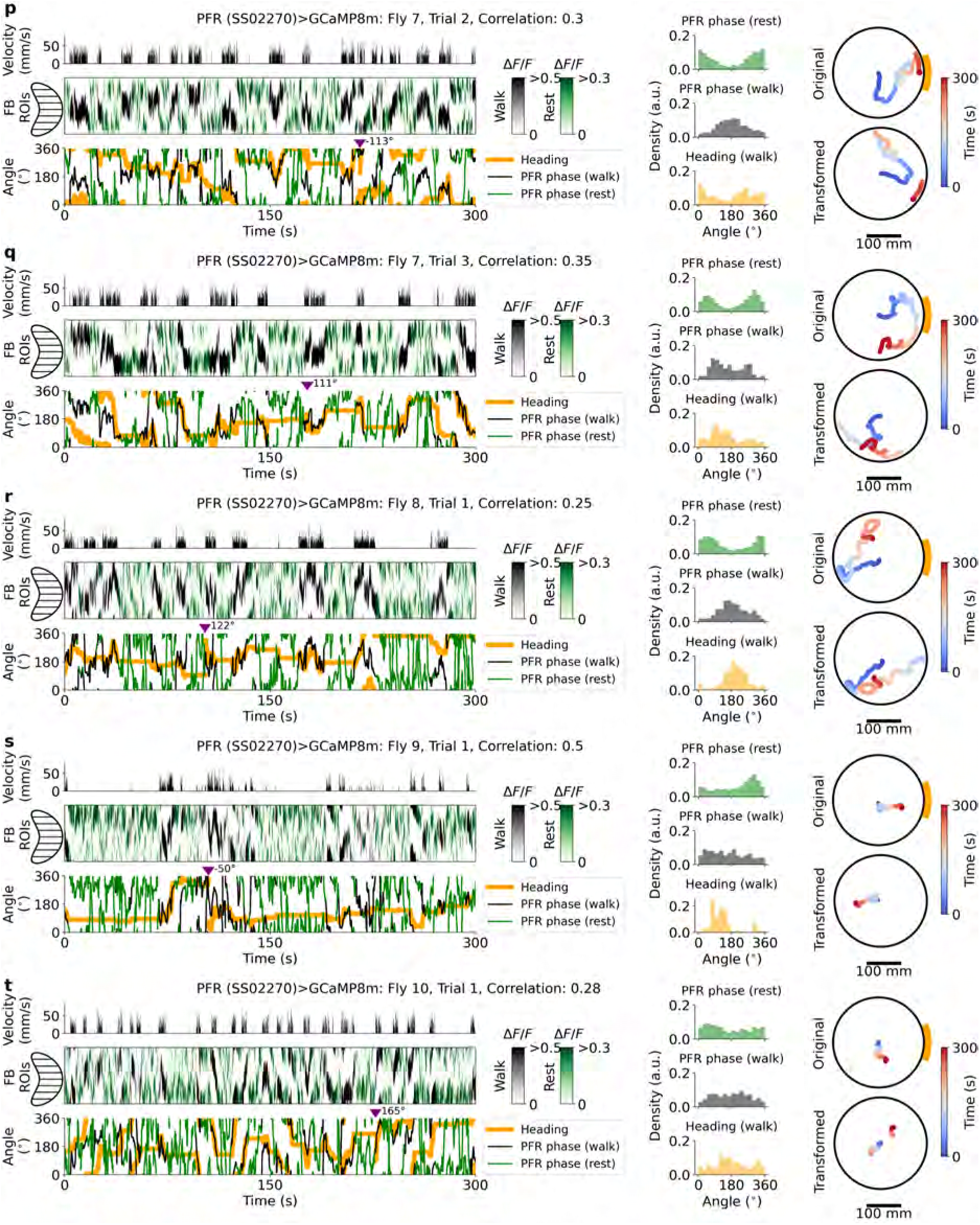
Neural activity and behavior in flies 7-10 expressing GCaMP8m in PFR neurons (split-GAL4 line SS02270) navigating the mechanical VR. **p-t** Same as Supplementary Fig. S7a-e.

**Figure S11.**
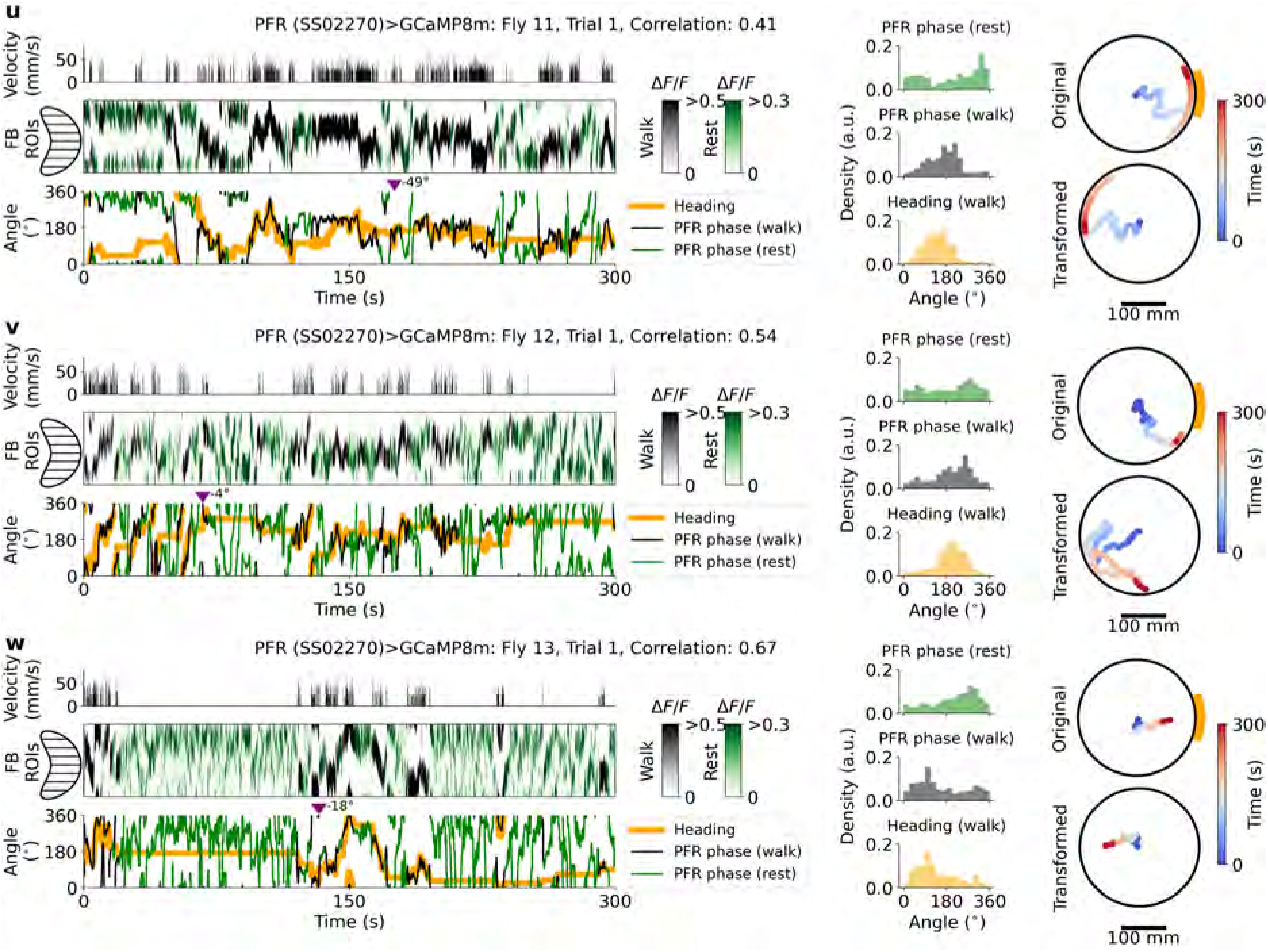
Neural activity and behavior in flies 11-13 expressing GCaMP8m in PFR neurons (split-GAL4 line SS02270) navigating the mechanical VR **u-w** Same as Supplementary Fig. S7a-e.

**Figure S12.**
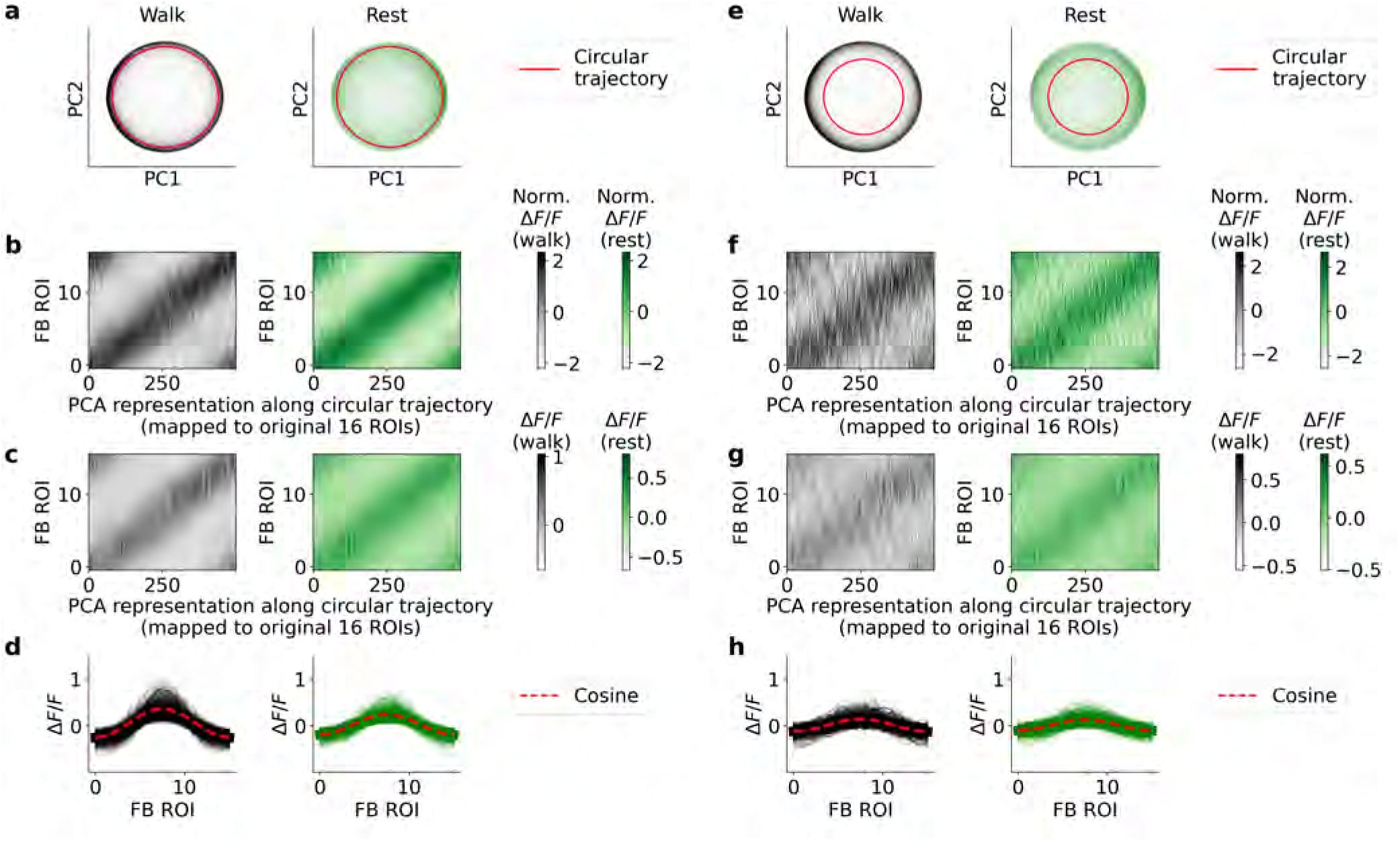
Same as Supplementary Fig. S5 but for PFR neurons using the R27G06-GAL4 line for a total fo 16 flies (see Table S1) resulting in a total of 205,360 fluorescence traces.

**Figure S13.**
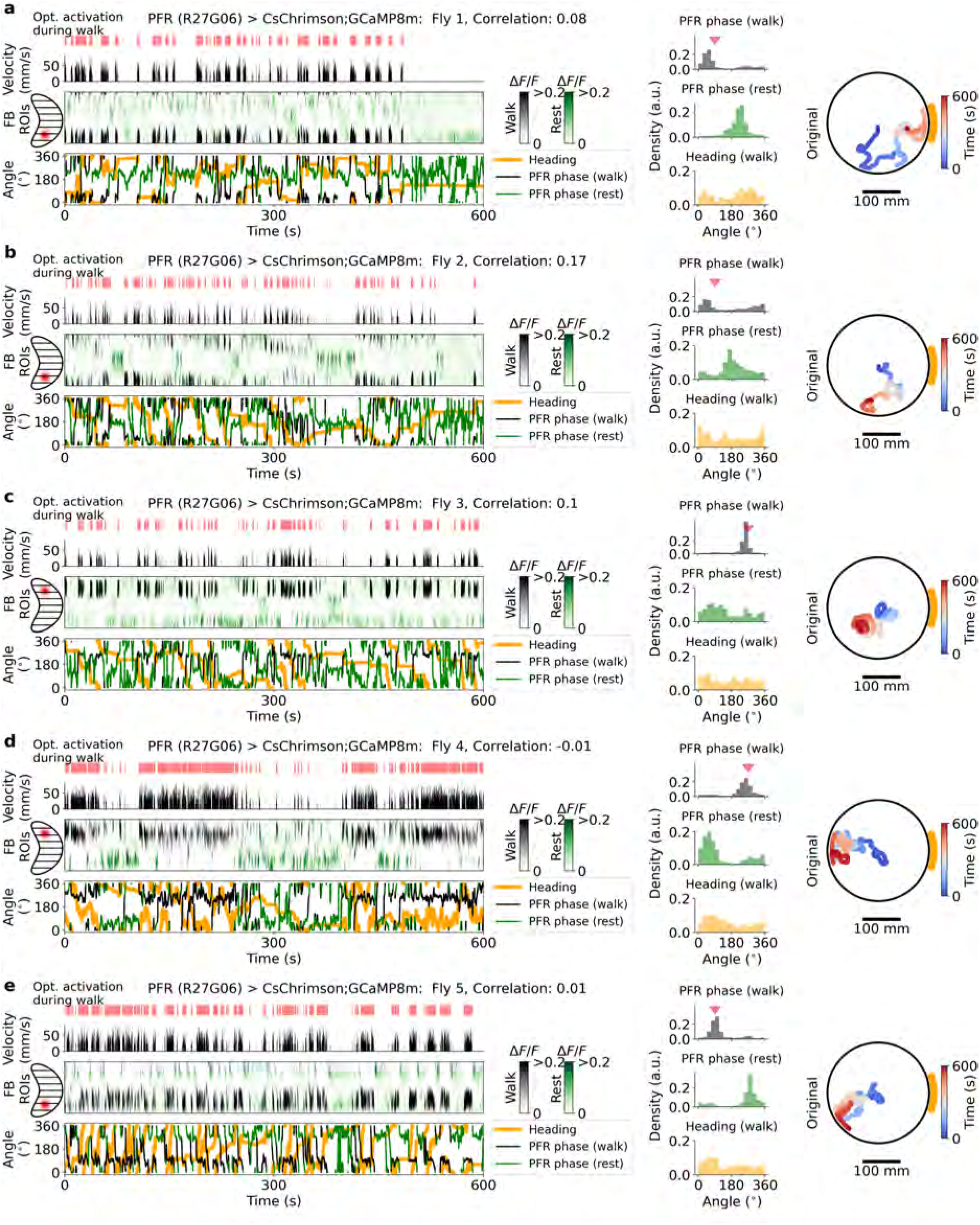
Experiments in flies 1-5 with GAL4 line R27G06 expressing CsChrimson and GCaMP8m in PFR neurons during optogenetic activation during walking periods. Data shown are from selected flies (10 minute trials) with *≥*60 s of walking and resting (selection criteria in Table S1)). **a-e** Left panel: Fly velocity (top); fluorescence in FB ROIs during walking (black) and rest (green) (middle); PFR phase during walking (black) and rest (green), and fly heading (orange) (bottom). Middle panel: PFR phase density during rest (green, top) and walking (black, middle); heading density (orange, bottom). Right panel: Original fly trajectory in the mechanical VR.

**Figure S14.**
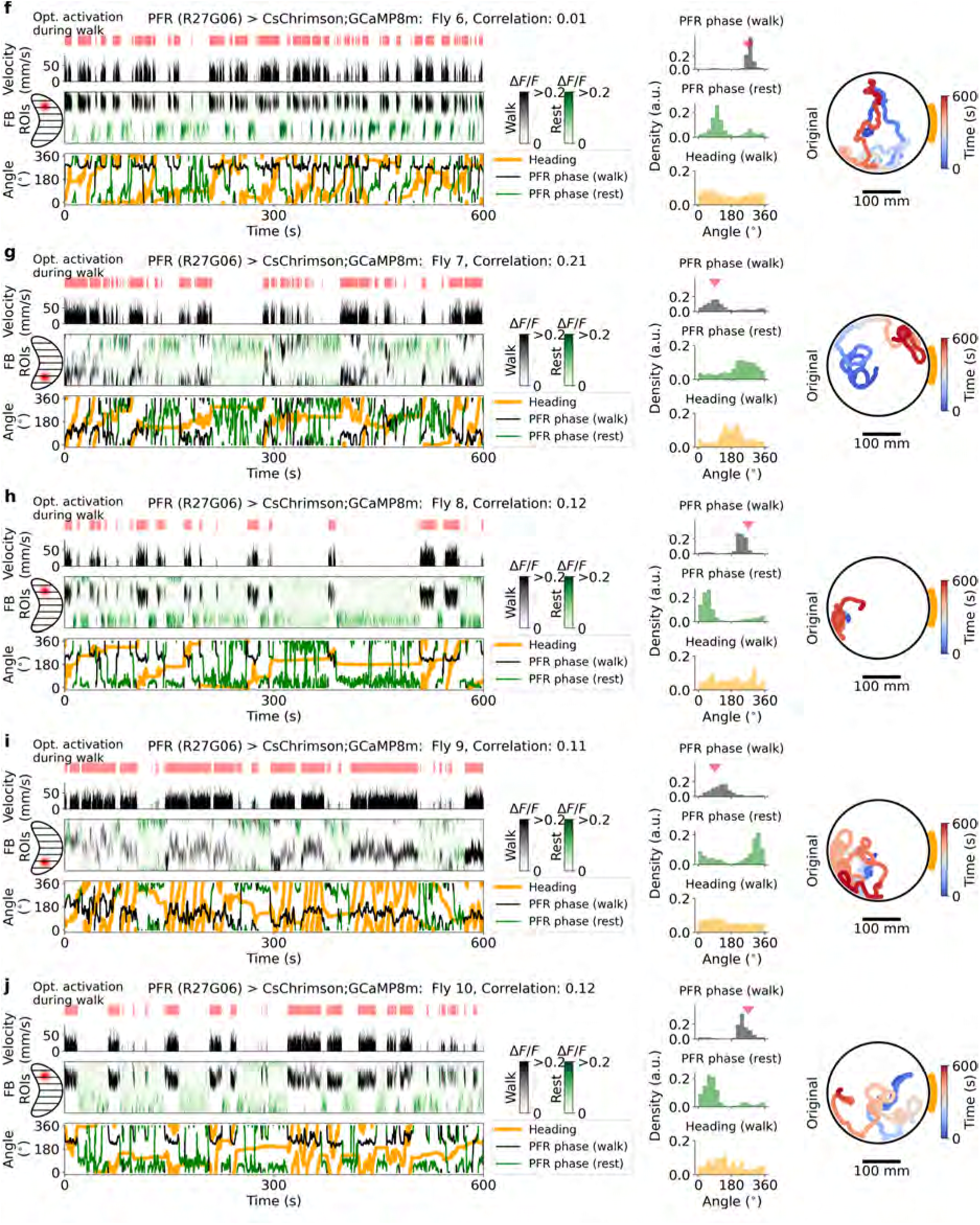
Experiments in flies 6-10 with GAL4 line R27G06 expressing CsChrimson and GCaMP8m in PFR neurons during optogenetic activation during walking periods. **f-j** Same as Supplementary Fig. S13a-e.

**Figure S15.**
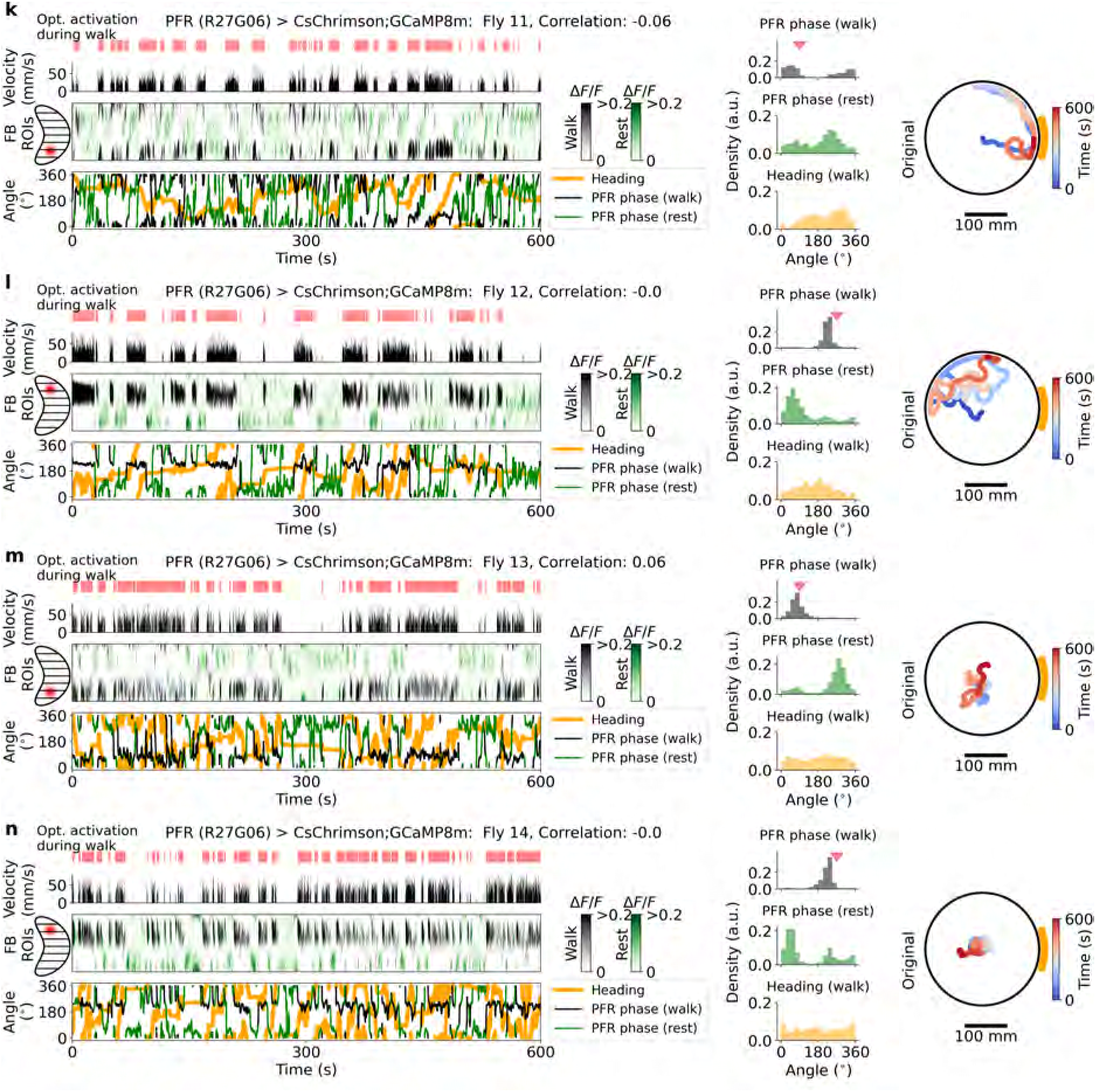
Experiments in flies 11-14 with GAL4 line R27G06 expressing CsChrimson and GCaMP8m in PFR neurons during optogenetic activation during walking periods. **k-n** Same as Supplementary Fig. S13a-e.

**Figure S16.**
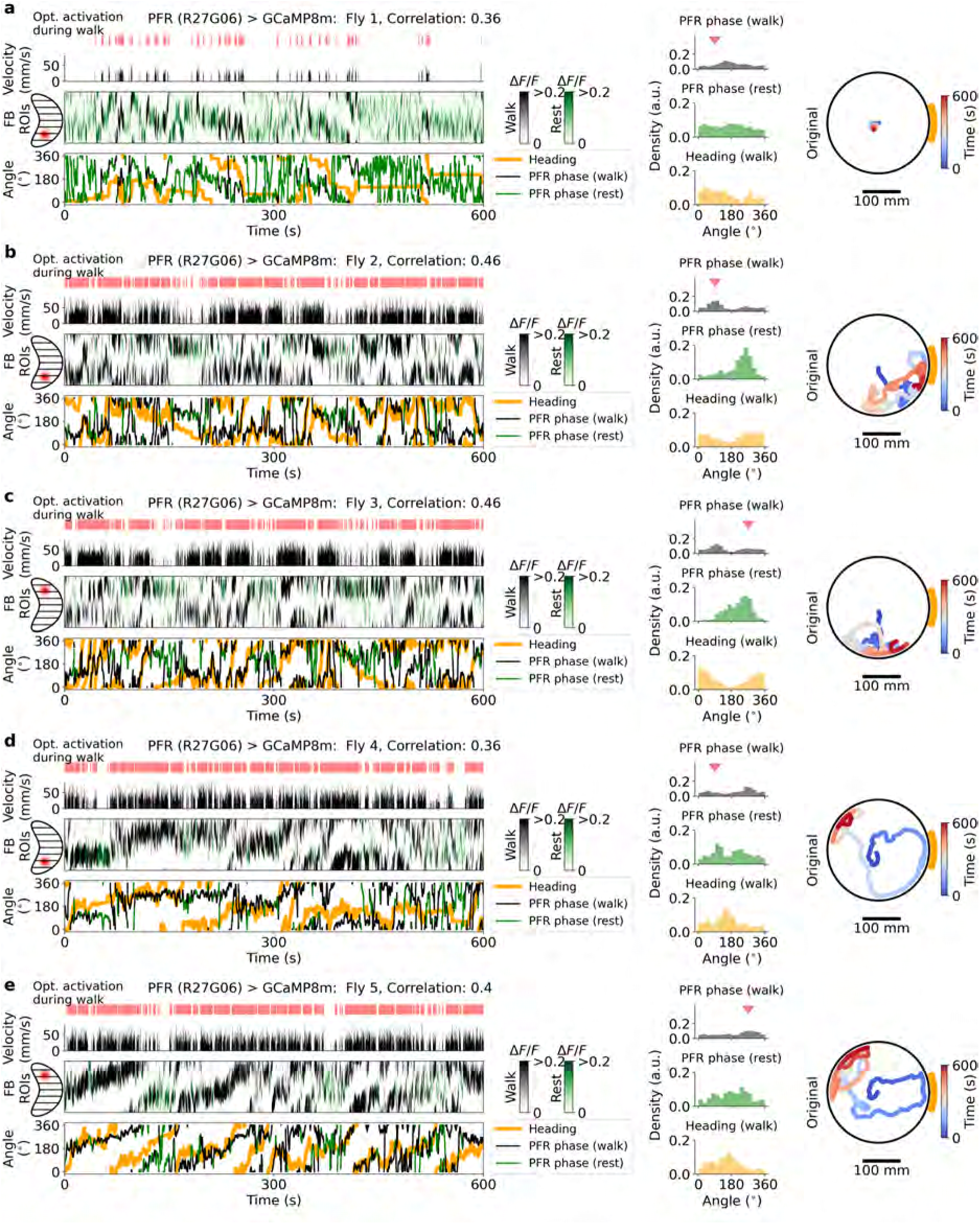
Experiments in flies 1-5 with GAL4 line R27G06 expressing GCaMP8m (control flies) in PFR neurons during optogenetic activation during walking periods. Data shown are from selected flies (10 minute trials) with *≥*60 s of walking and resting (selection criteria in Table S1). **a-e** Same as Supplementary Fig. S13a-e

**Figure S17.**
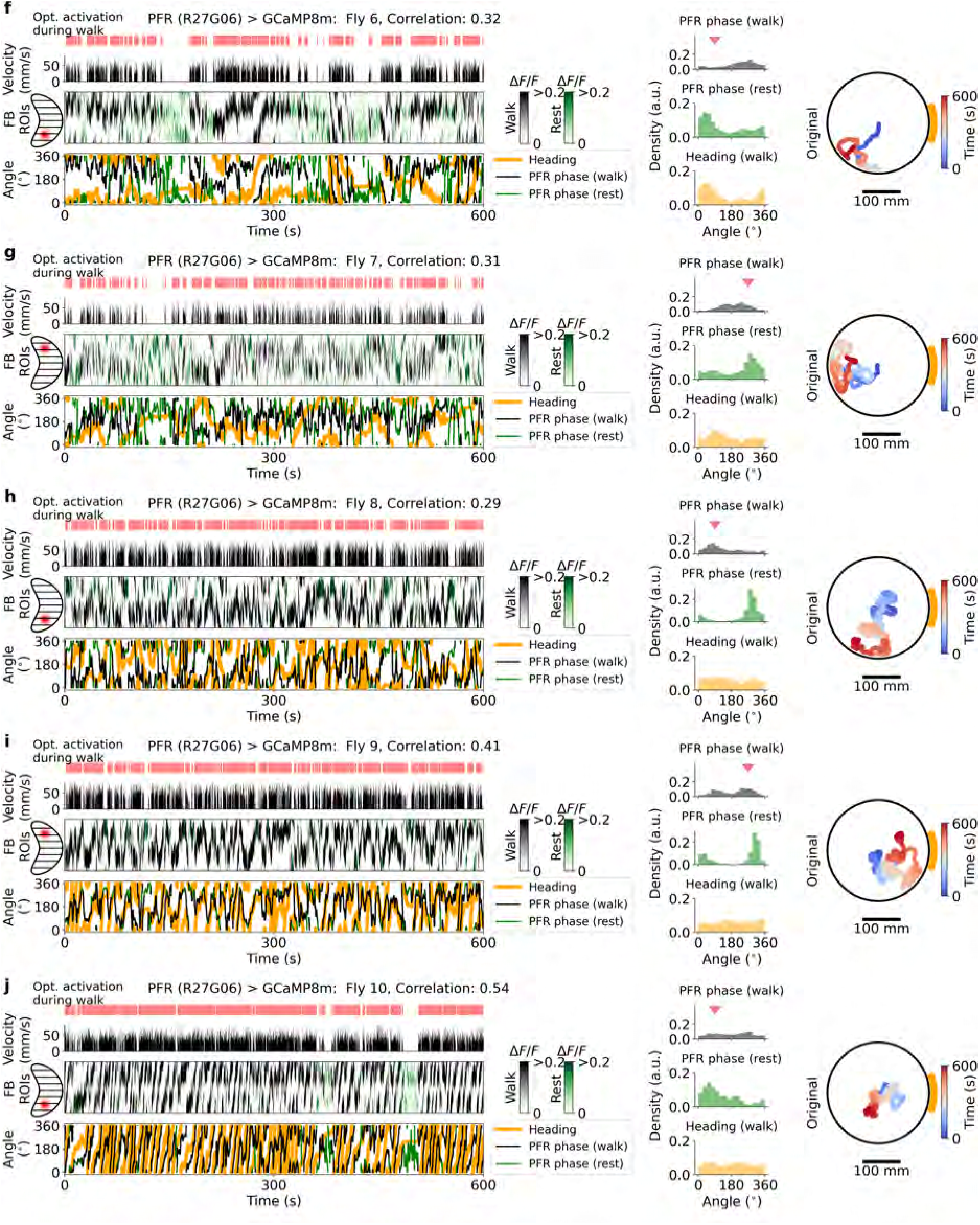
Experiments in flies 6-10 with GAL4 line R27G06 expressing GCaMP8m (control flies) in PFR neurons during optogenetic activation during walking periods. **f-j** Same as Supplementary Fig. S13a-e.

**Figure S18.**
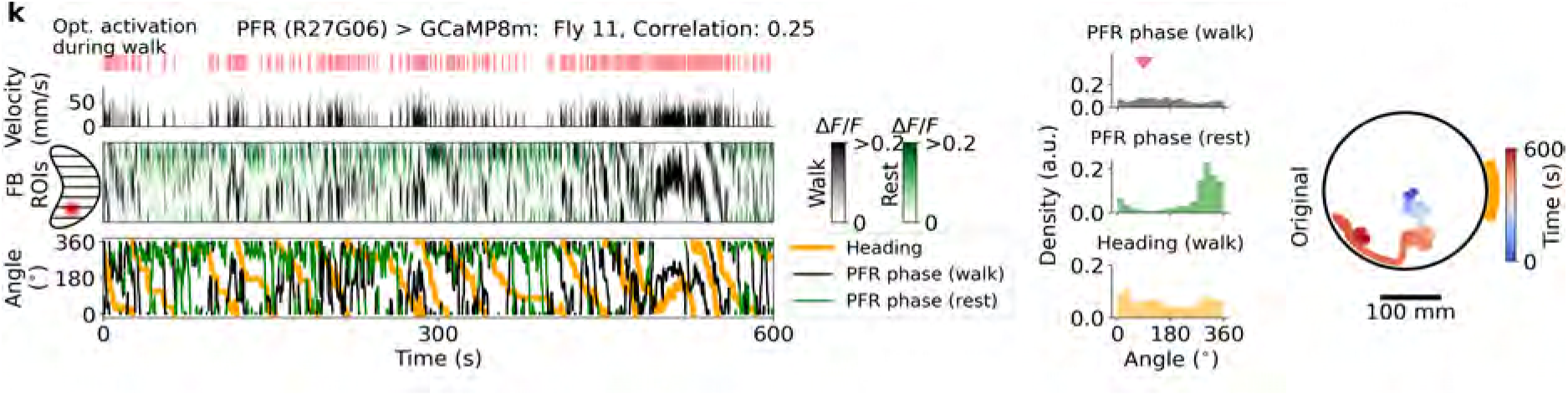
Experiment in fly 11 with GAL4 line R27G06 expressing GCaMP8m (control flies) in PFR neurons during optogenetic activation during walking periods. **k-l** Same as Supplementary Fig. S13a-e.

**Figure S19.**
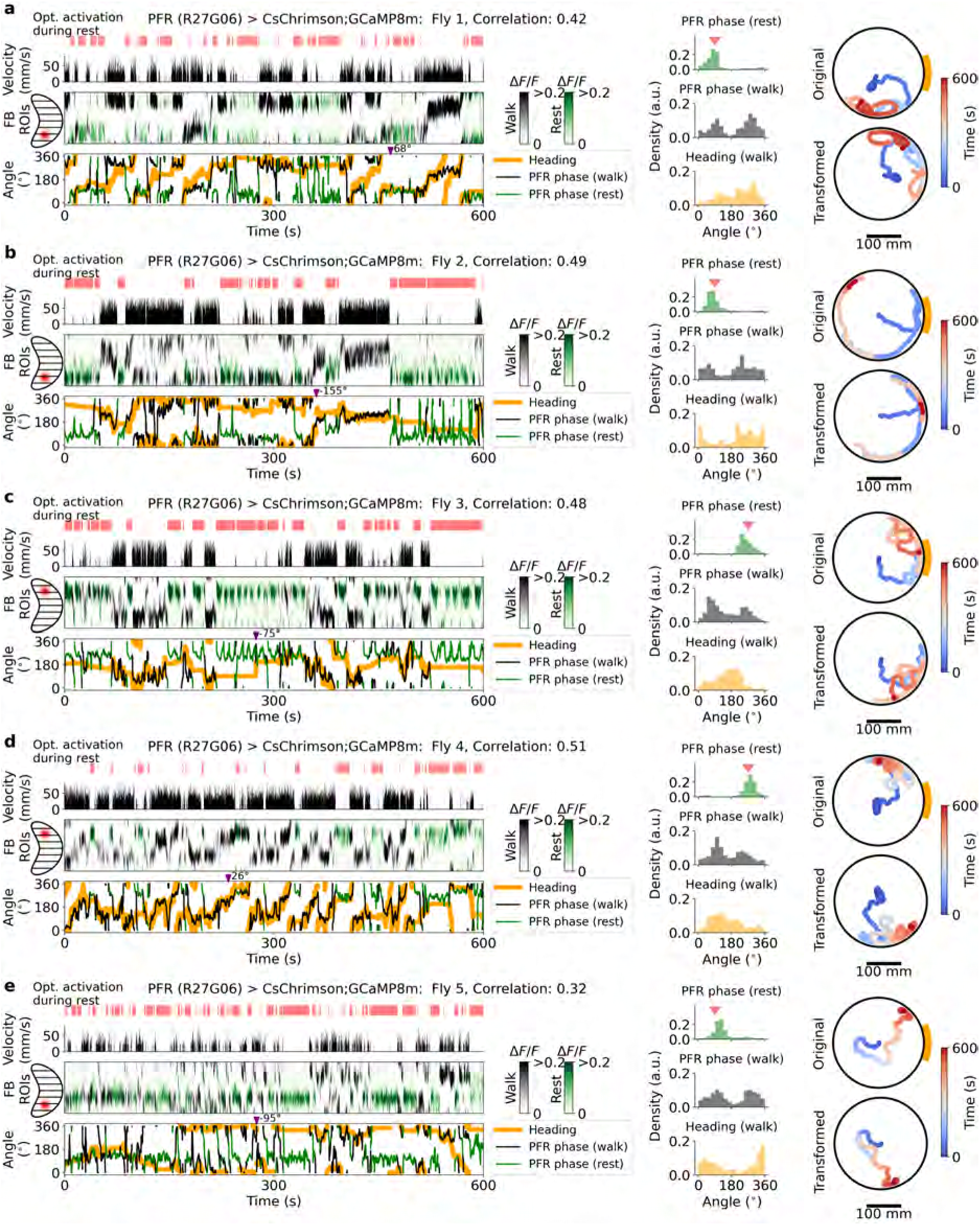
Experiments in flies 1-5 with GAL4 line R27G06 expressing CsChrimson and GCaMP8m in PFR neurons during optogenetic activation during rest periods. Data shown are from selected flies (10 minute trials) with *≥*60 s of walking and resting and a PFR phase-heading correlation *≥* 0.2 (selection criteria in Table S1). **a-e** Left panel: Fly velocity (top); fluorescence in FB ROIs during walking (black) and rest (green) (middle); PFR phase during walking (black) and rest (green), and fly heading (orange) (bottom). Middle panel: PFR phase density during rest (green, top) and walking (black, middle); heading density (orange, bottom). Right panel: Original fly trajectory in the mechanical VR (top) and transformed trajectory (buttom) using the two-segment offset model (see Methods and Extended Data Fig. 3h).

**Figure S20.**
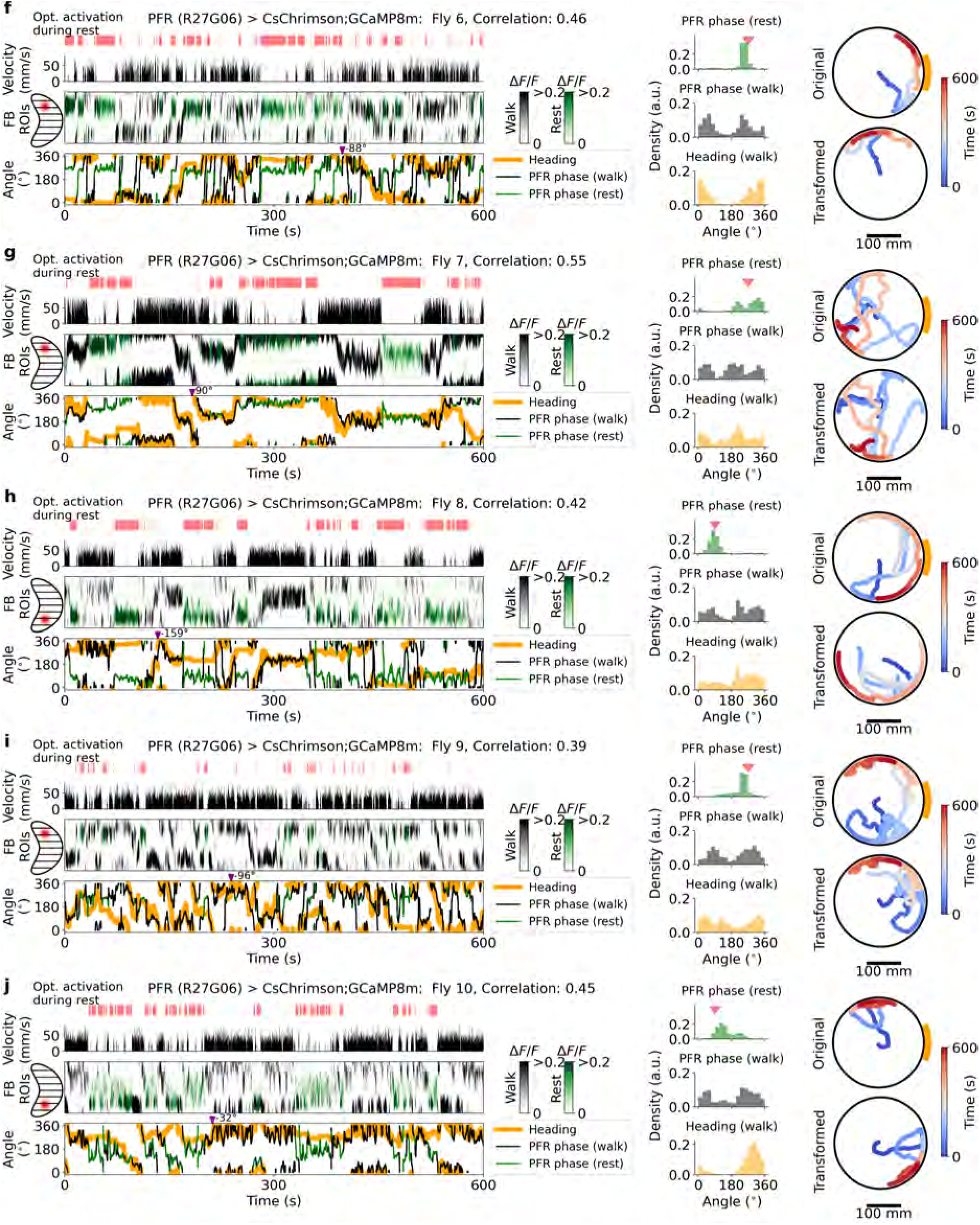
Experiments in flies 6-10 with GAL4 line R27G06 expressing CsChrimson and GCaMP8m in PFR neurons during optogenetic activation during rest periods **f-j** Same as Supplementary Fig. S19a-e.

**Figure S21.**
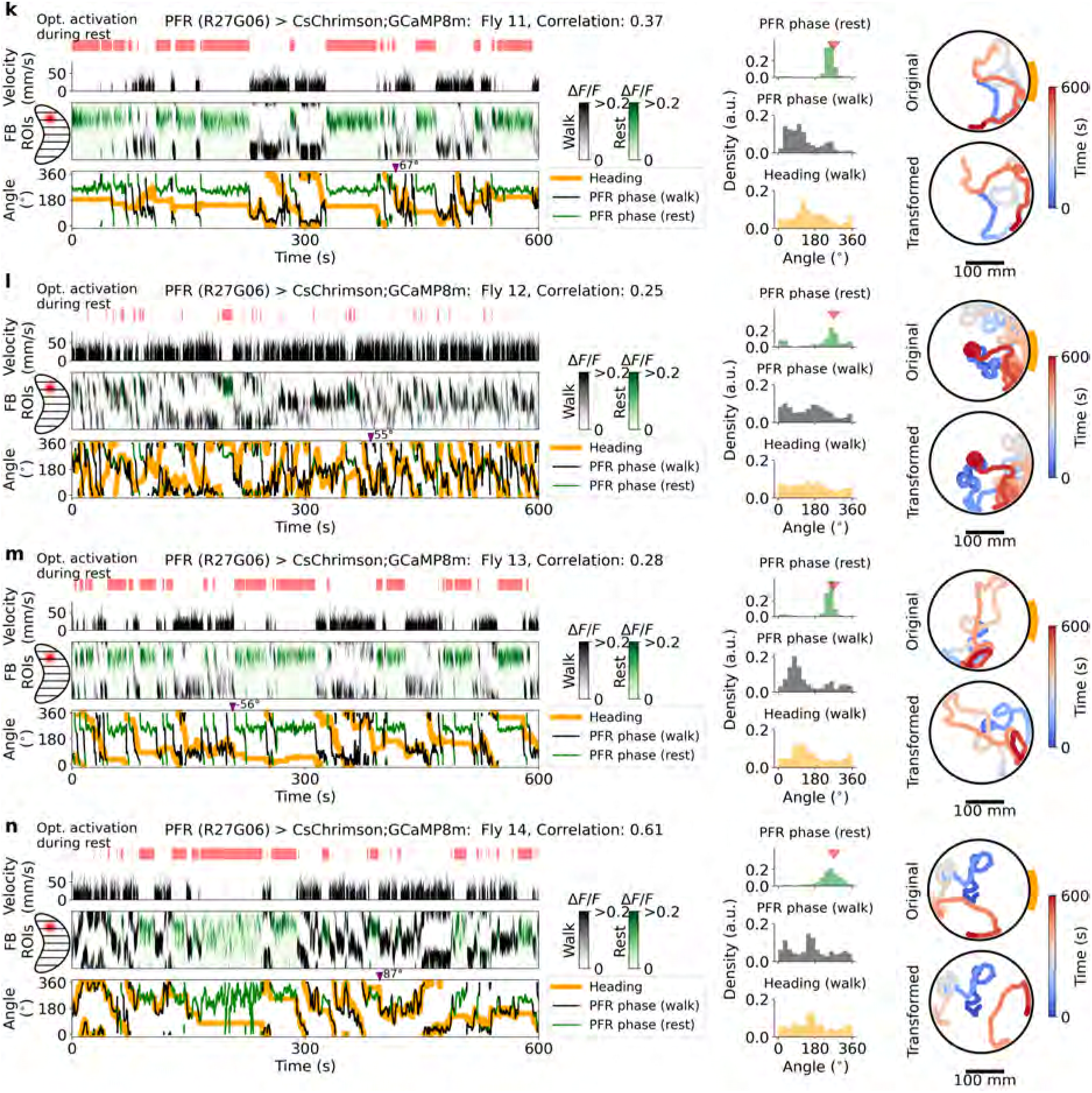
Experiments in flies 11-14 with GAL4 line R27G06 expressing CsChrimson and GCaMP8m in PFR neurons during optogenetic activation during rest periods **k-n** Same as Supplementary Fig. S19a-e.

**Figure S22.**
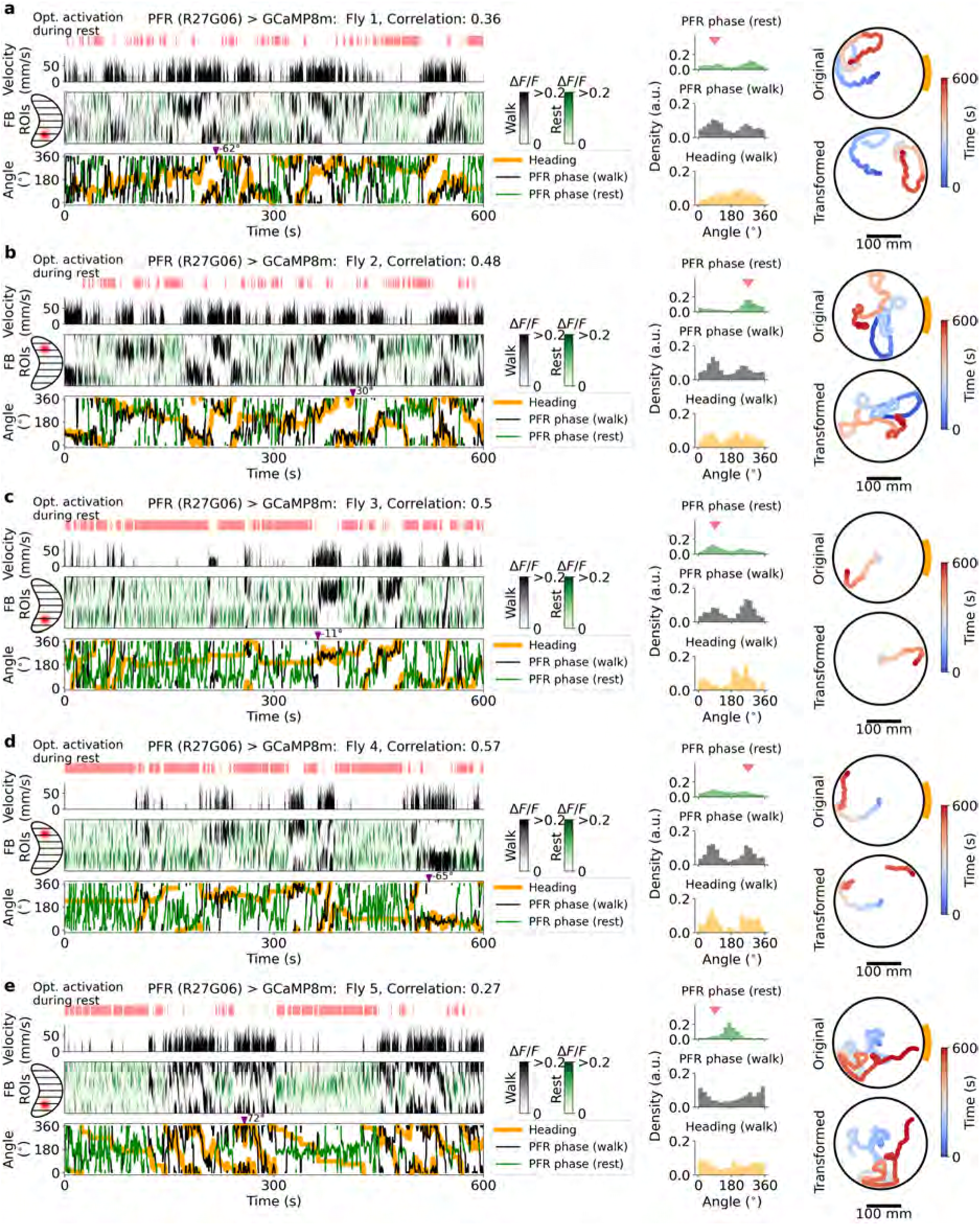
Experiments in flies 1-5 with GAL4 line R27G06 expressing GCaMP8m (control flies) in PFR neurons during optogenetic activation during rest periods. Data shown are from selected flies (10 minute trials) with *≥*60 s of walking and resting and a PFR phase-heading correlation *≥* 0.2 (selection criteria in Table S1). **a-e** Same as Supplementary Fig. S19a-e

**Figure S23.**
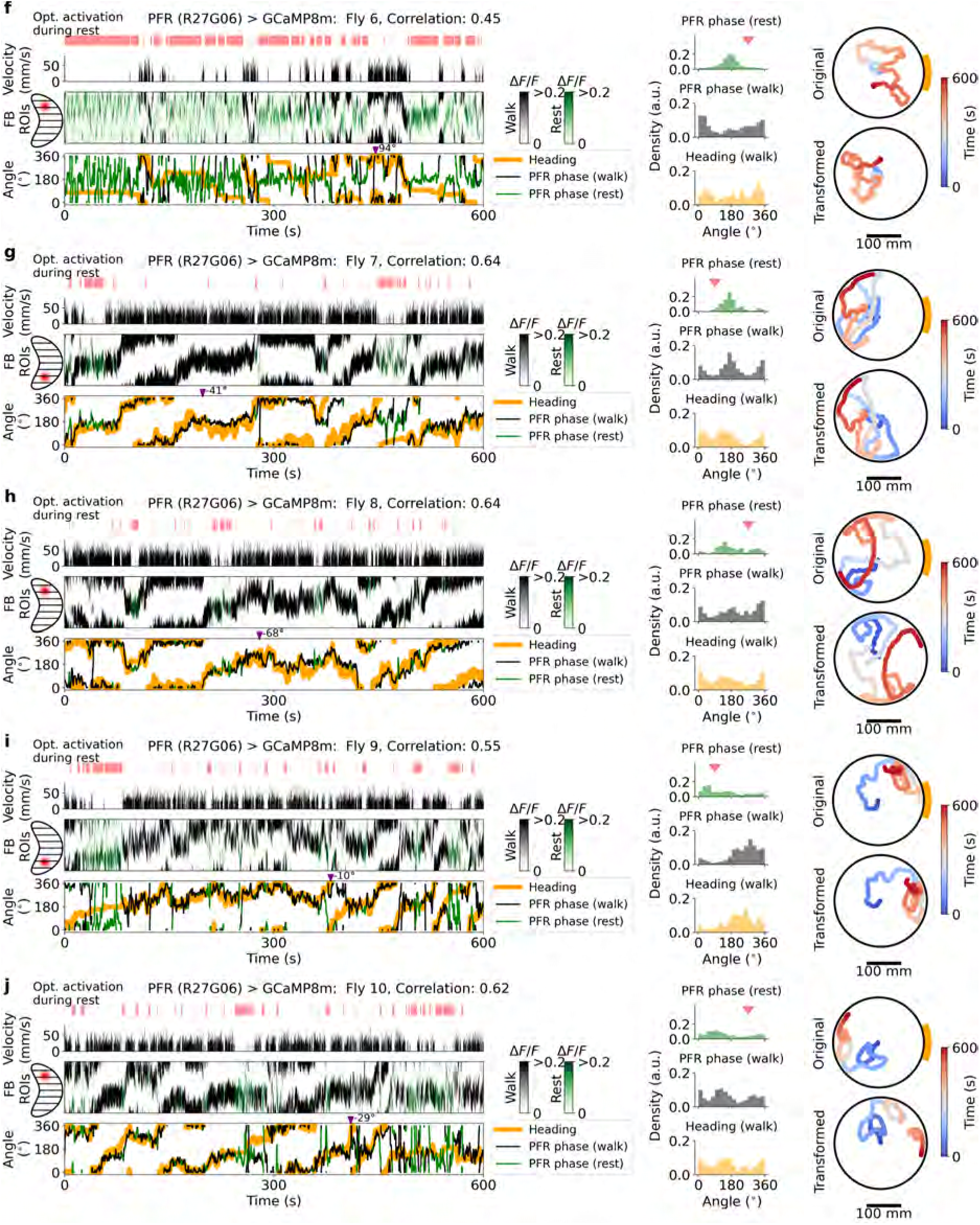
Experiments in flies 6-10 with GAL4 line R27G06 expressing GCaMP8m (control flies) in PFR neurons during optogenetic activation during rest periods. **f-j** Same as Supplementary Fig. S19a-e.

**Figure S24.**
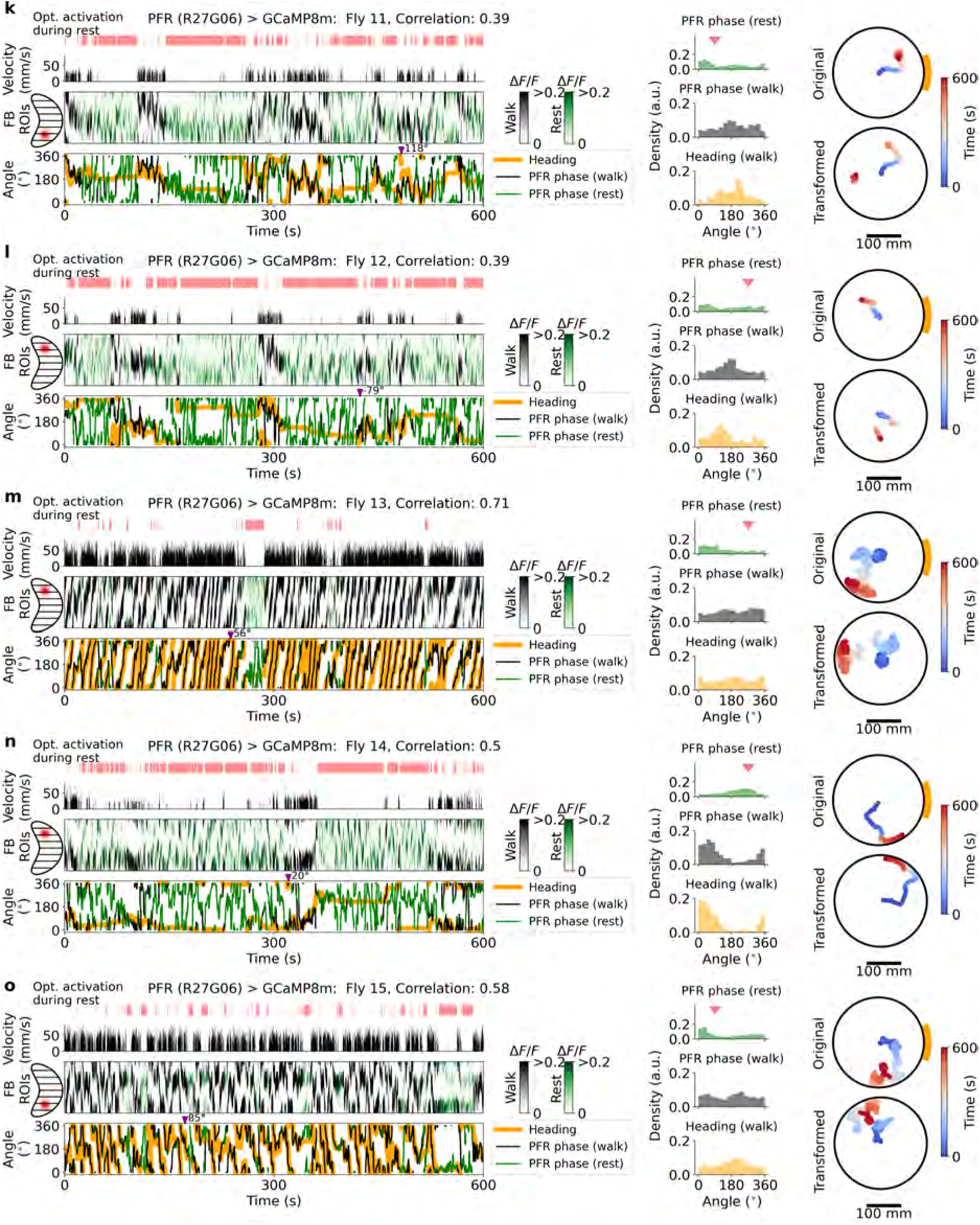
Experiments in flies 11-15 with GAL4 line R27G06 expressing GCaMP8m (control flies) in PFR neurons during optogenetic activation during rest periods. **k-o** Same as Supplementary Fig. S19a-e.

**Figure S25.**
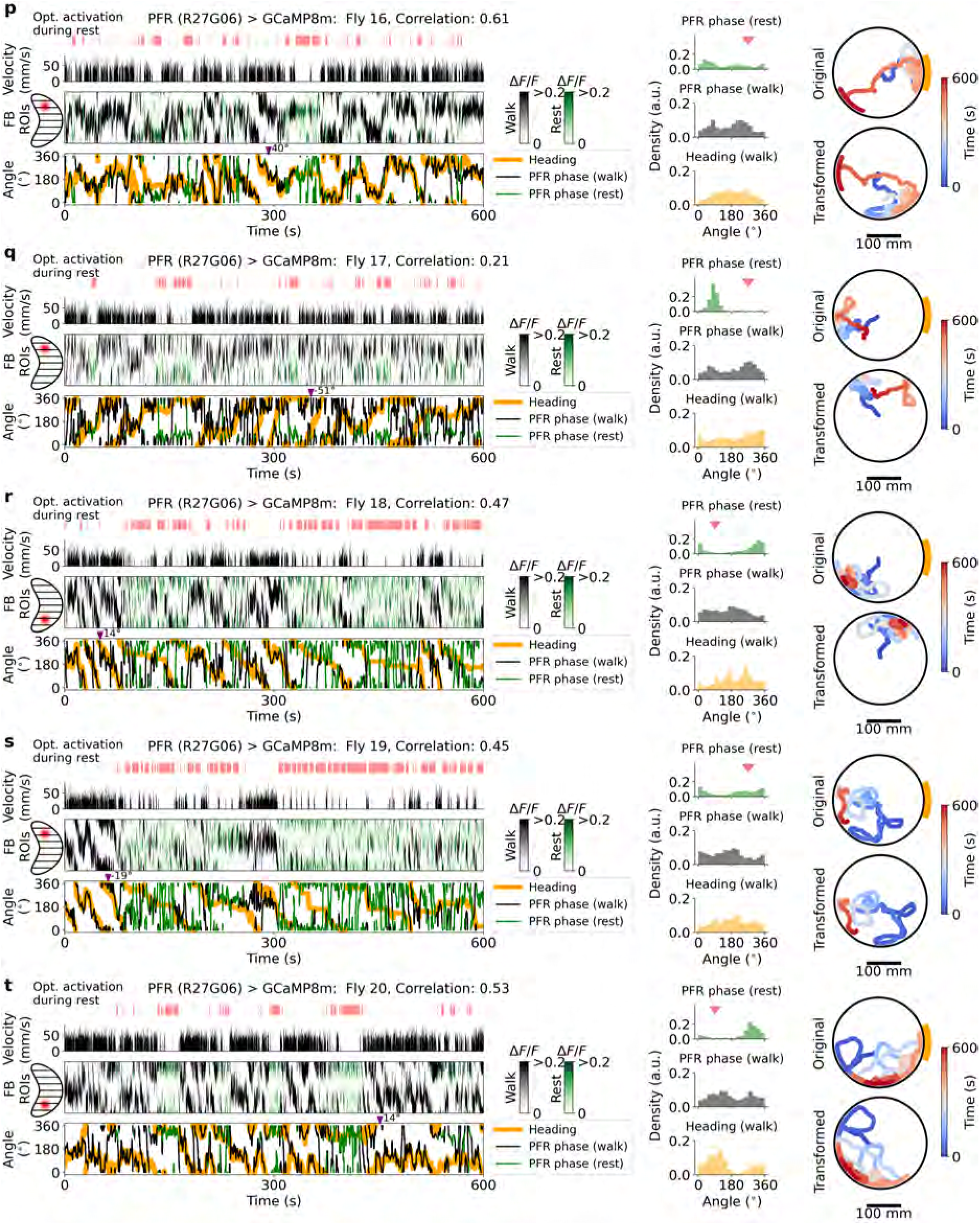
Experiments in flies 16-20 with GAL4 line R27G06 expressing GCaMP8m (control flies) in PFR neurons during optogenetic activation during rest periods. **p-t** Same as Supplementary Fig. S19a-e.

**Figure S26.**
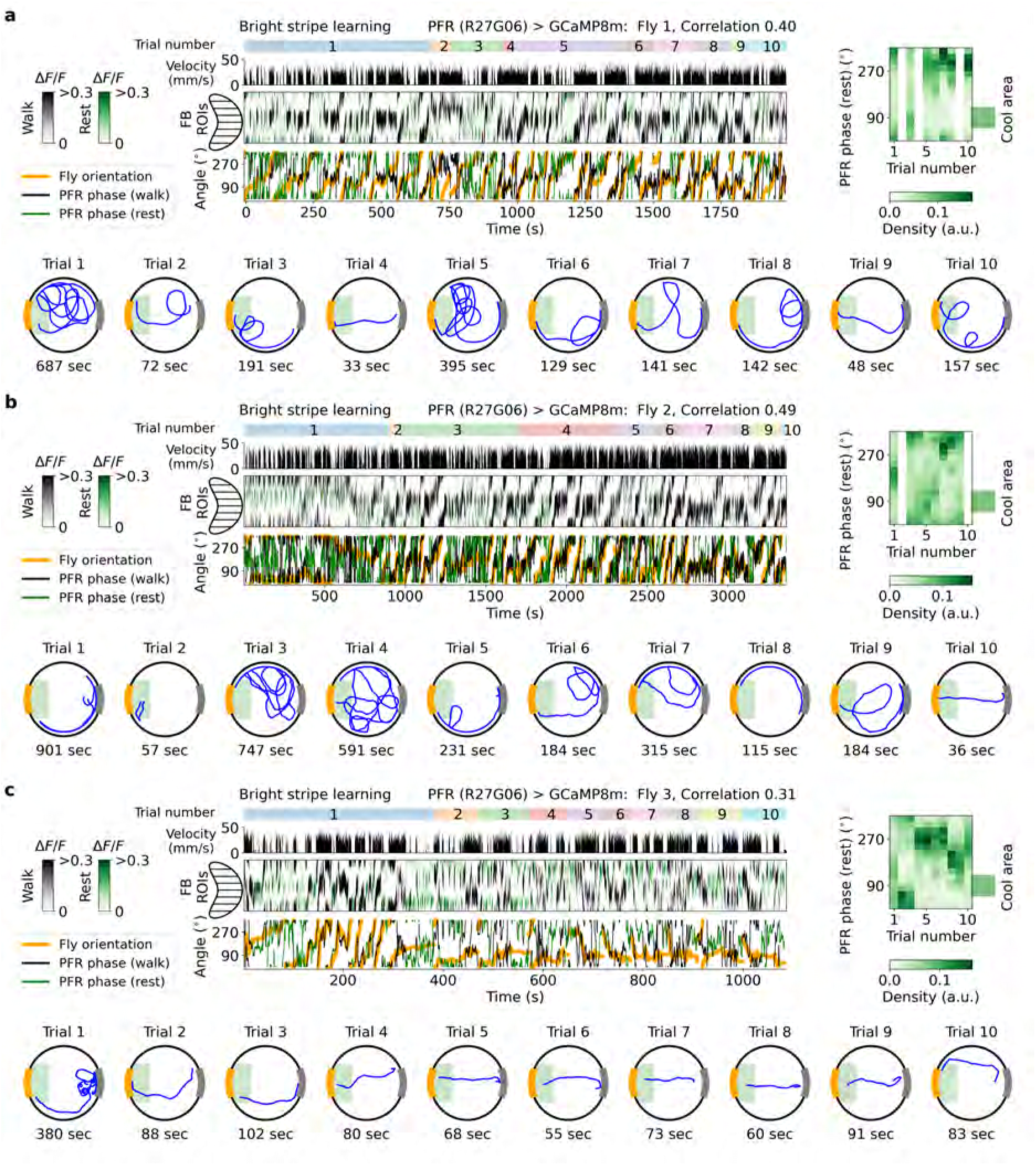
Experiments in flies 1-3 with GAL4 line R27G06 expressing GCaMP8m during place learning toward the bright stripe over 10 trials. Selected flies had a learning index > 1 (see Methods) and a PFR phase–heading correlation *≥* 0.2 (see Table S1). **a-c** Top left panel: Trial number (1-10, indicated by top colored regions) and velocity (top, black). Fluorescence during walking (black) and rest (green) across trials (middle), and PFR phase during walking (black) and rest (green) and heading (orange, bottom). Top right panel: PFR phase density during rest across trials. Vertical white regions indicate trials with less than 10 seconds of resting time; these distributions were excluded from the analysis. The green square indicates the direction of the cool area in the arena. Bottom panel: Original fly trajectories across trials.

**Figure S27.**
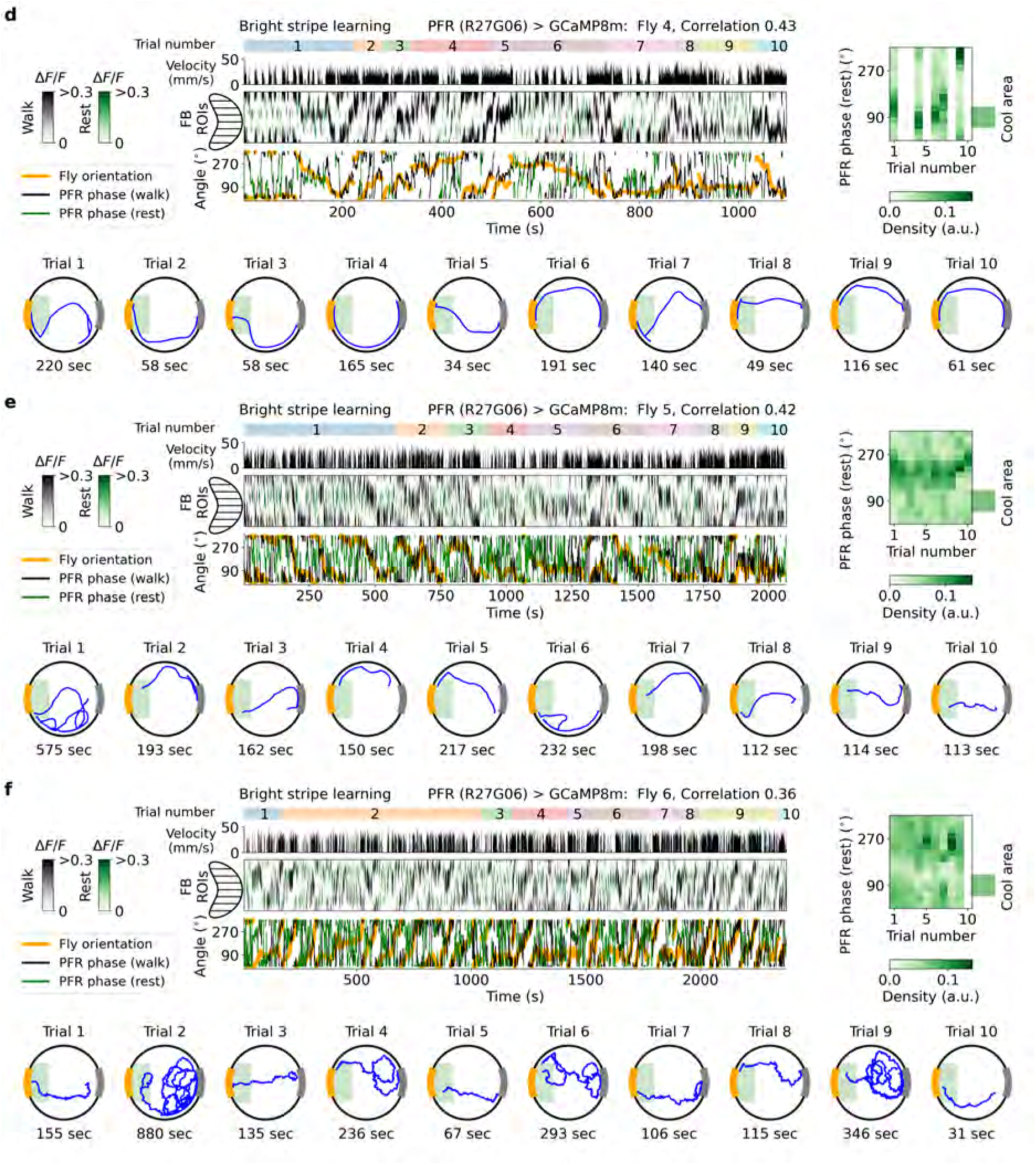
Experiments in flies 4-6 with GAL4 line R27G06 expressing GCaMP8m during place learning toward the bright stripe over 10 trials. **d-f** Same as Supplementary Fig. S26a-c.

**Figure S28.**
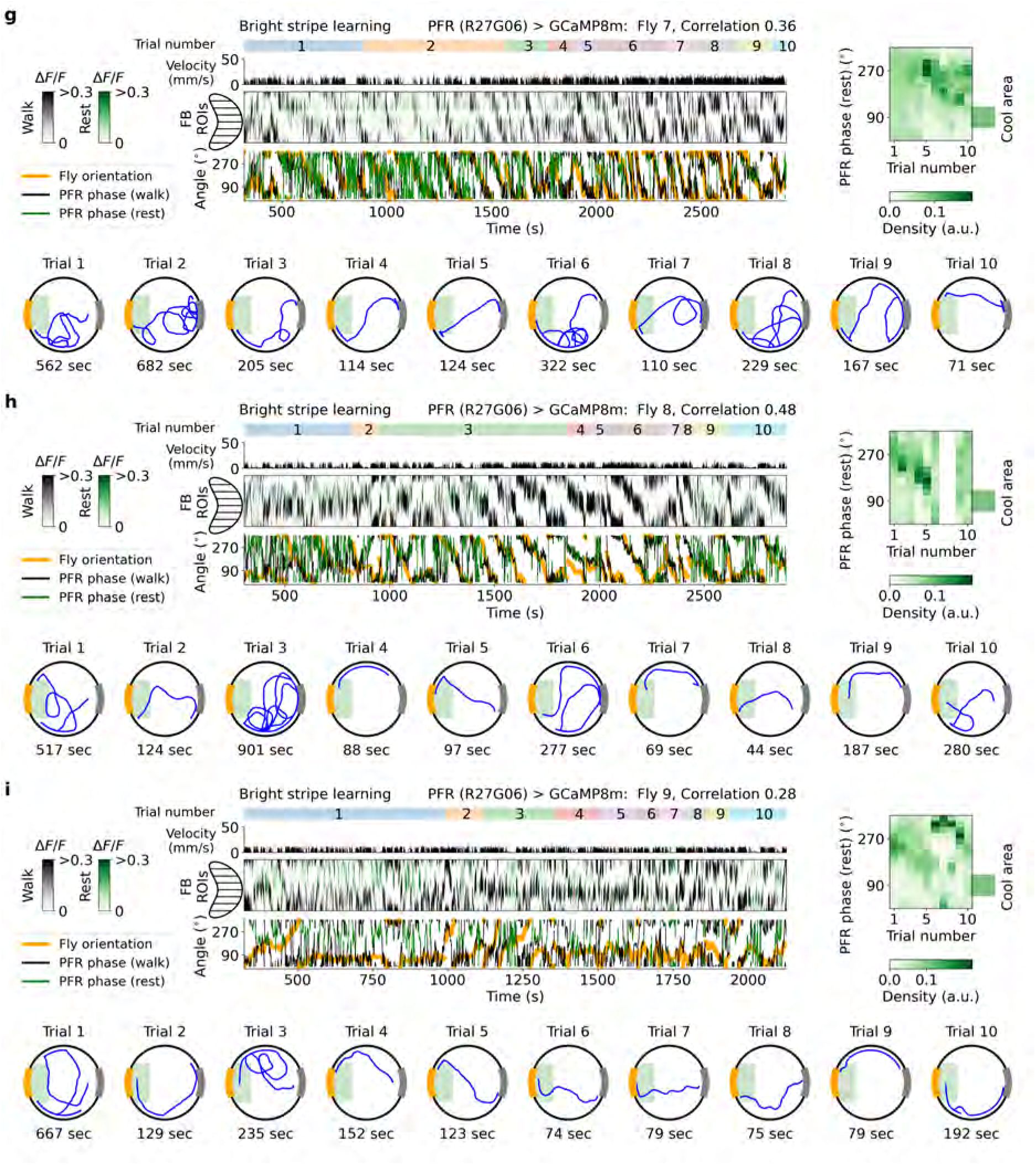
Experiments in flies 7-9 with GAL4 line R27G06 expressing GCaMP8m during place learning toward the bright stripe over 10 trials. **g-i** Same as Supplementary Fig. S26a-c.

**Figure S29.**
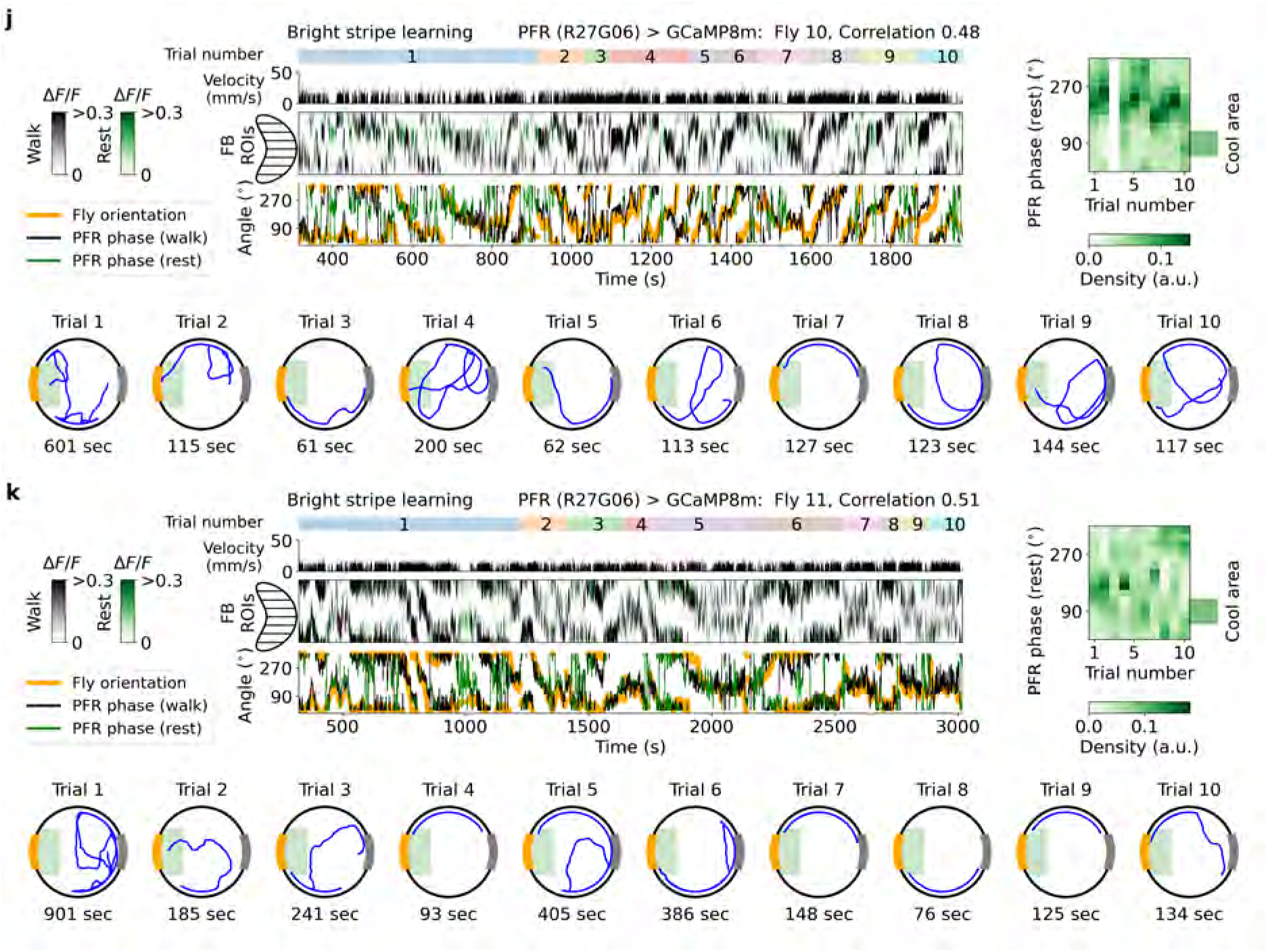
Experiments in flies 10-11 with GAL4 line R27G06 expressing GCaMP8m during place learning toward the bright stripe over 10 trials. **j-k** Same as Supplementary Fig. S26a-c.

**Figure S30.**
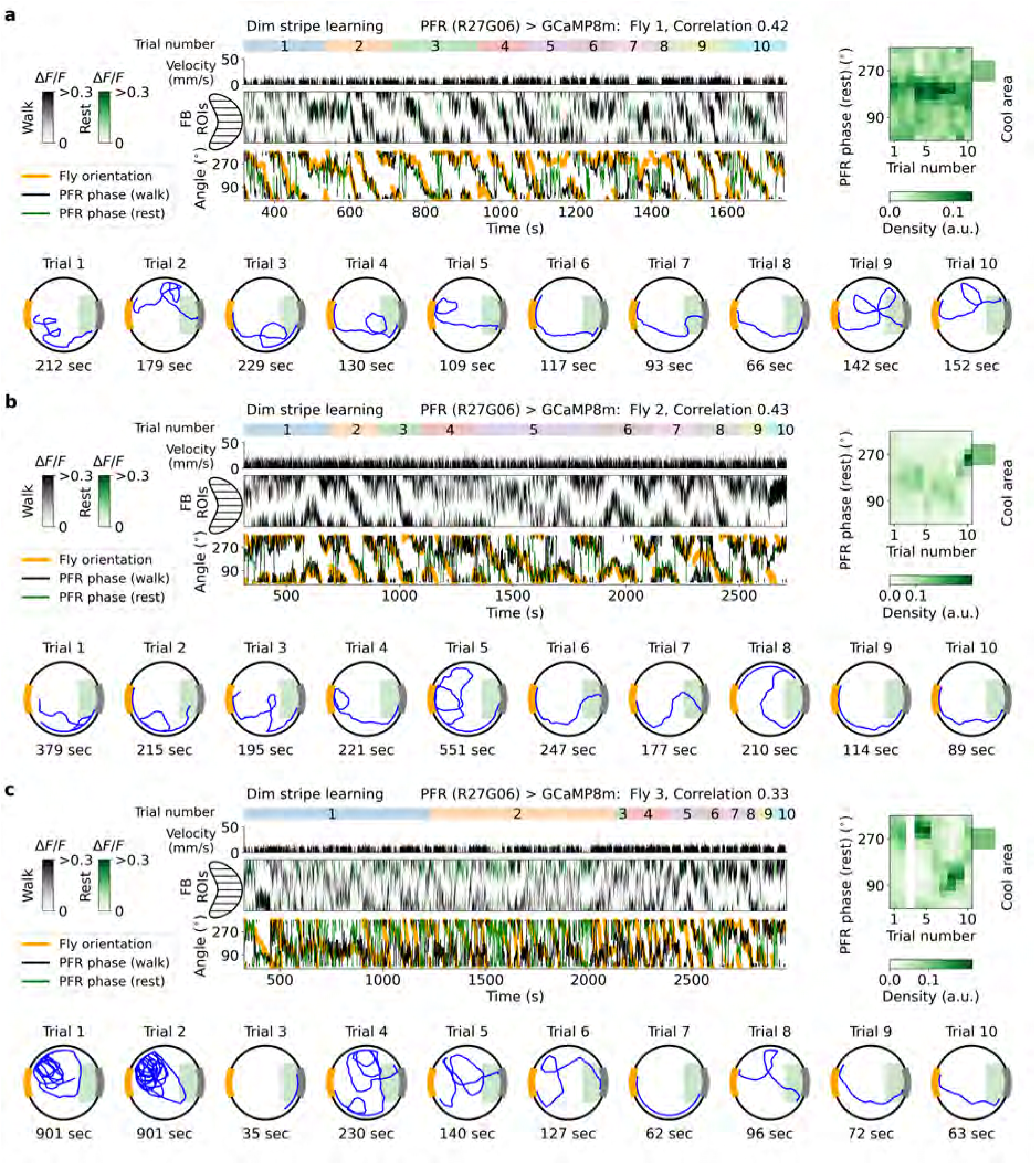
Experiments in flies 1-3 with GAL4 line R27G06 expressing GCaMP8m during place learning toward the dim stripe over 10 trials. **a-c** Same as Supplementary Fig. S26a-c.

**Figure S31.**
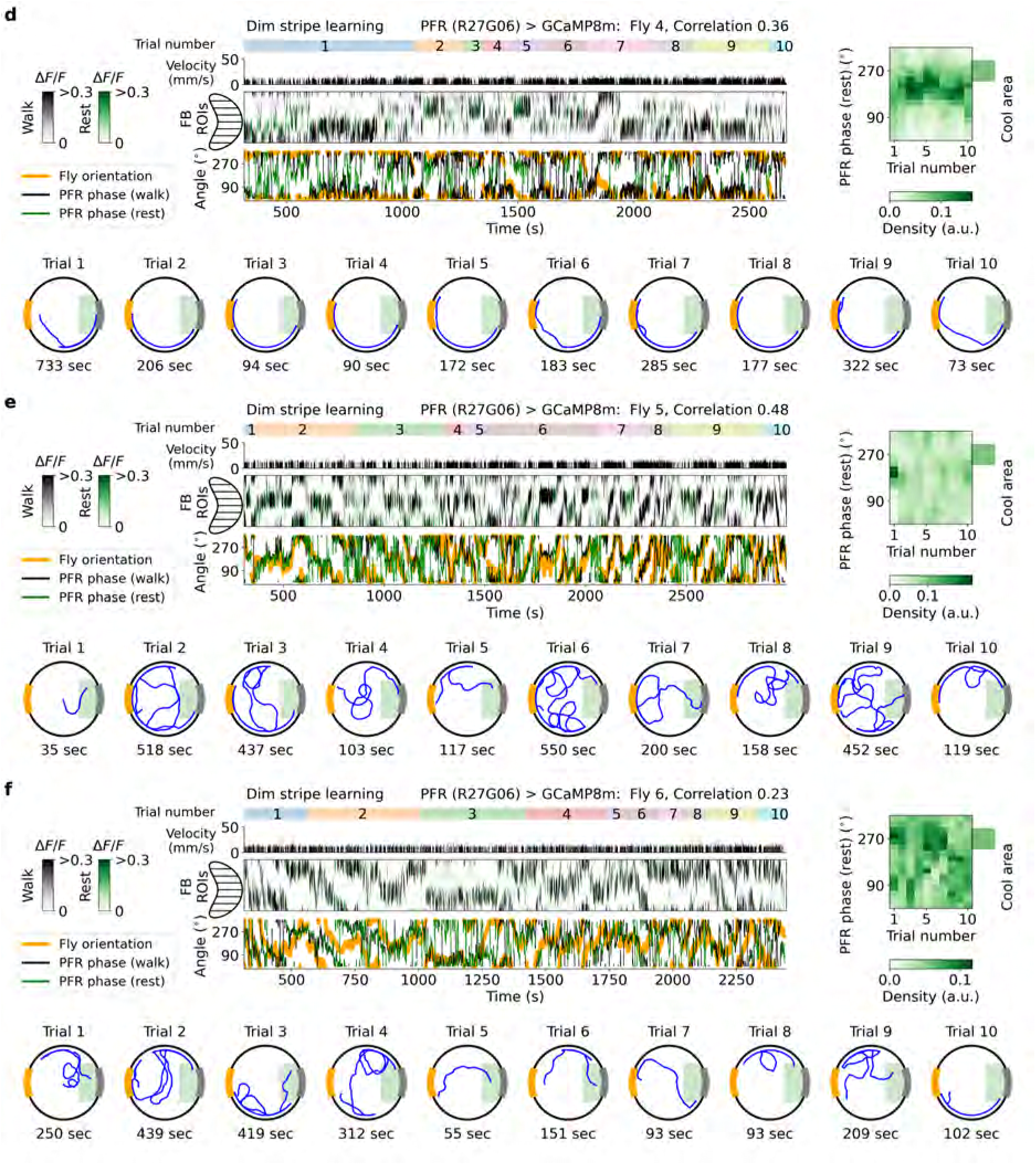
Experiments in flies 4-6 with GAL4 line R27G06 expressing GCaMP8m during place learning toward the dim stripe over 10 trials. **d-f** Same as Supplementary Fig. S26a-c.

**Figure S32.**
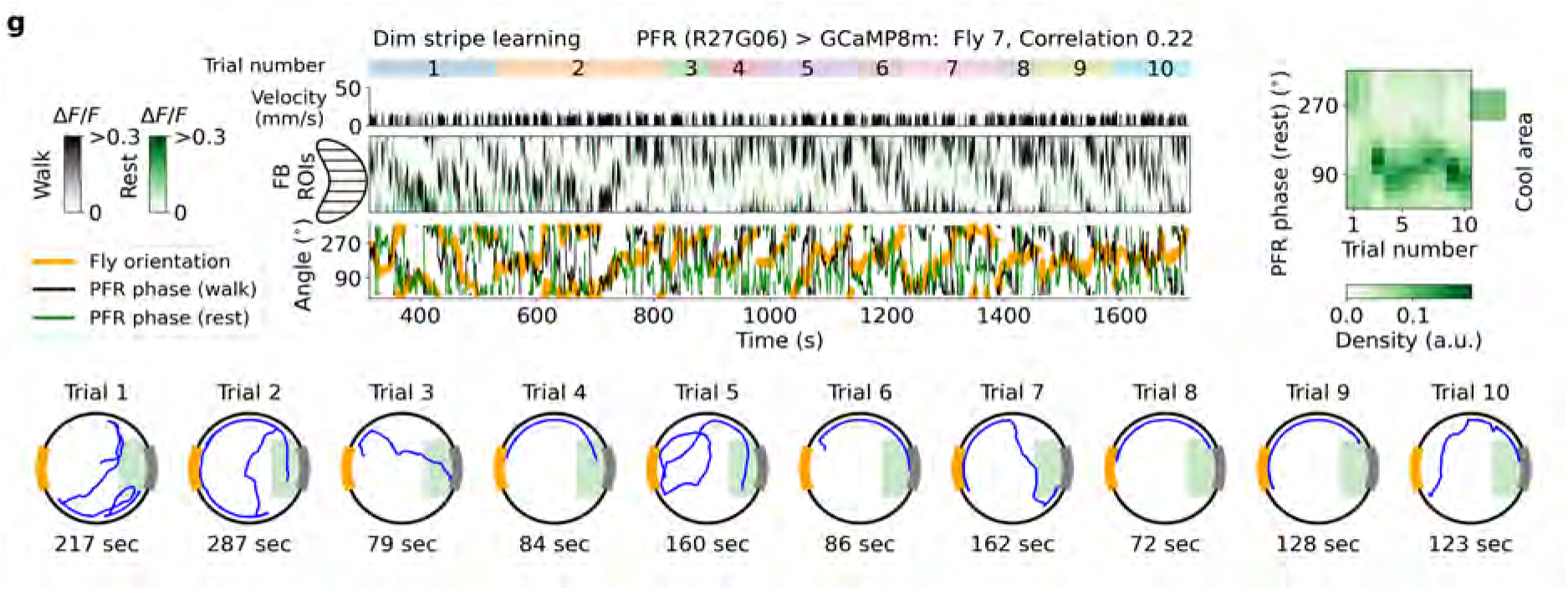
Experiments in fly 7 with GAL4 line R27G06 expressing GCaMP8m during place learning toward the dim stripe over 10 trials. **g** Same as Supplementary Fig. S26a-c.

**Figure S33.**
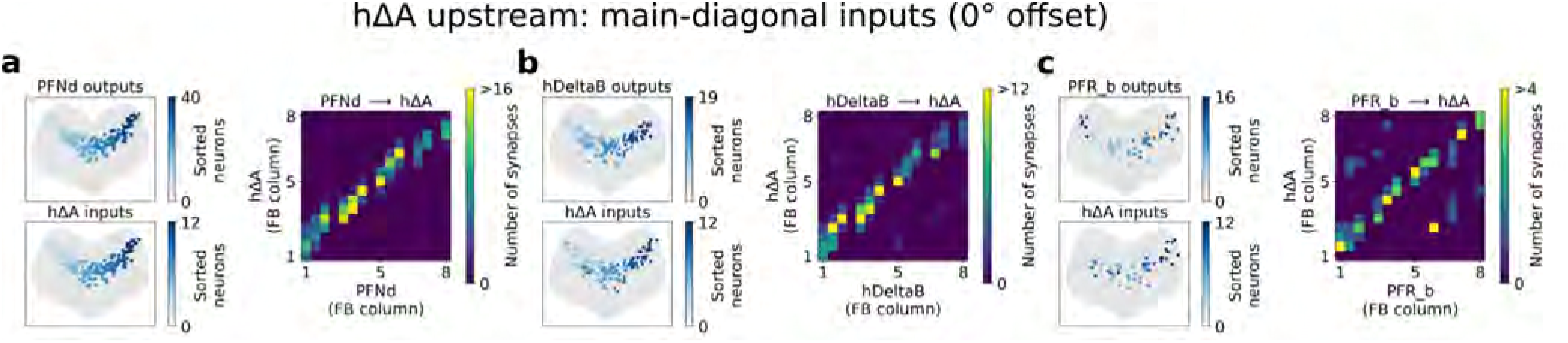
Connectivity profile between hΔA neurons and upstream populations that project to hΔA exclusively through main-diagonal connections. **All panels**: synaptic connections within the fan-shaped body (FB), shown in axon-view. Top left: synapses colored by the presynaptic upstream neuron; columnar upstream populations are sorted according to their output column in the FB (axonal column), from left to right. Left Bottom: the same synapses colored by the postsynaptic hΔA neuron, sorted by its axonal output column. Right: synapse counts between each upstream population and hΔA as a function of FB column, illustrating the main-diagonal connectivity pattern.

**Figure S34.**
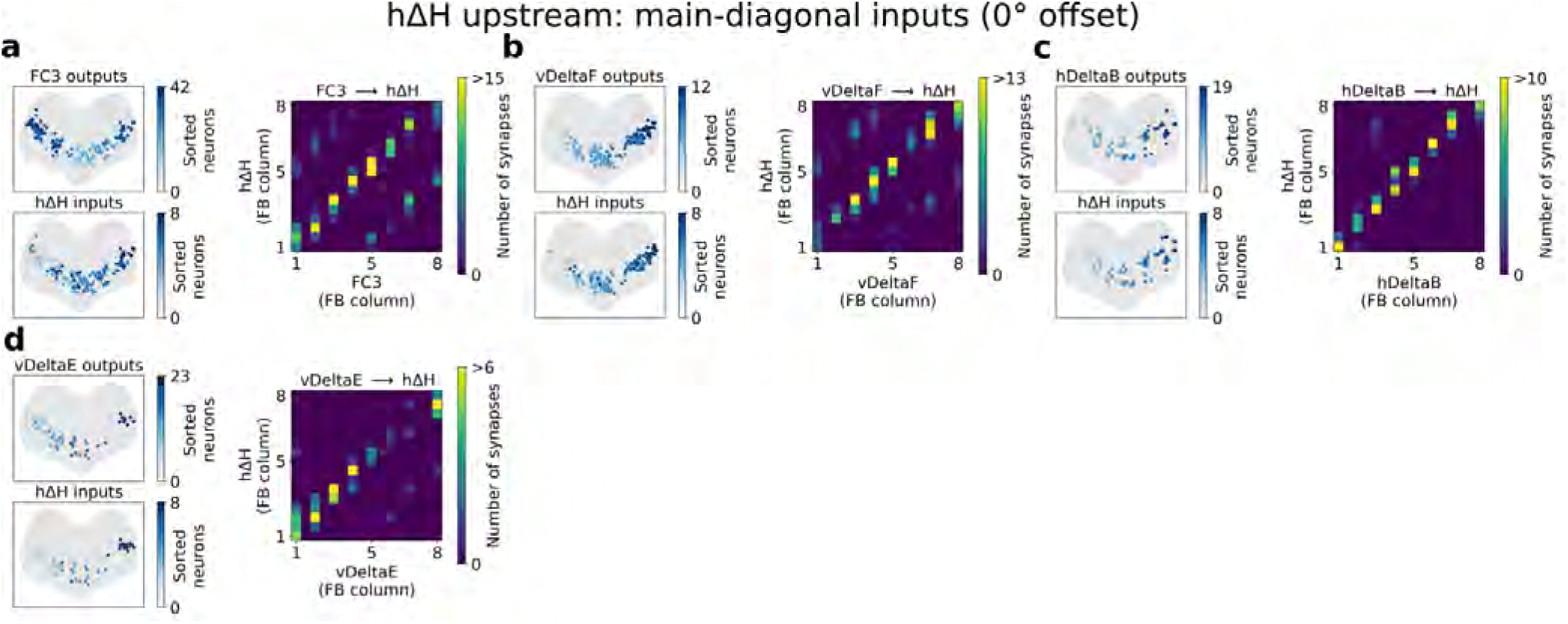
Same as Supplementary Fig. S33 but showing only upstream populations that target hΔH through the main diagonal.

**Figure S35.**
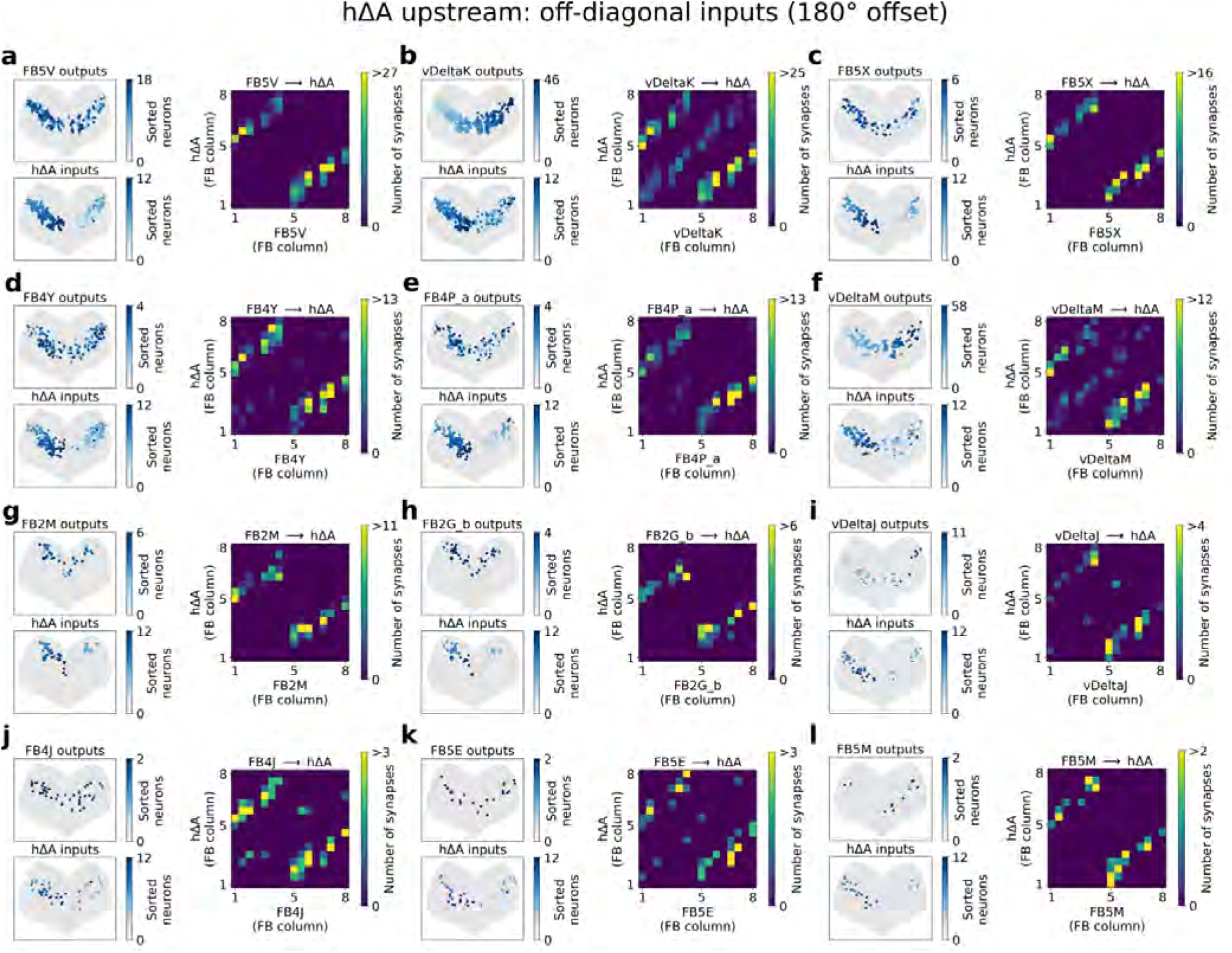
Same as Supplementary Fig. S33 but showing only upstream populations that target hΔA through the off diagonal.

**Figure S36.**
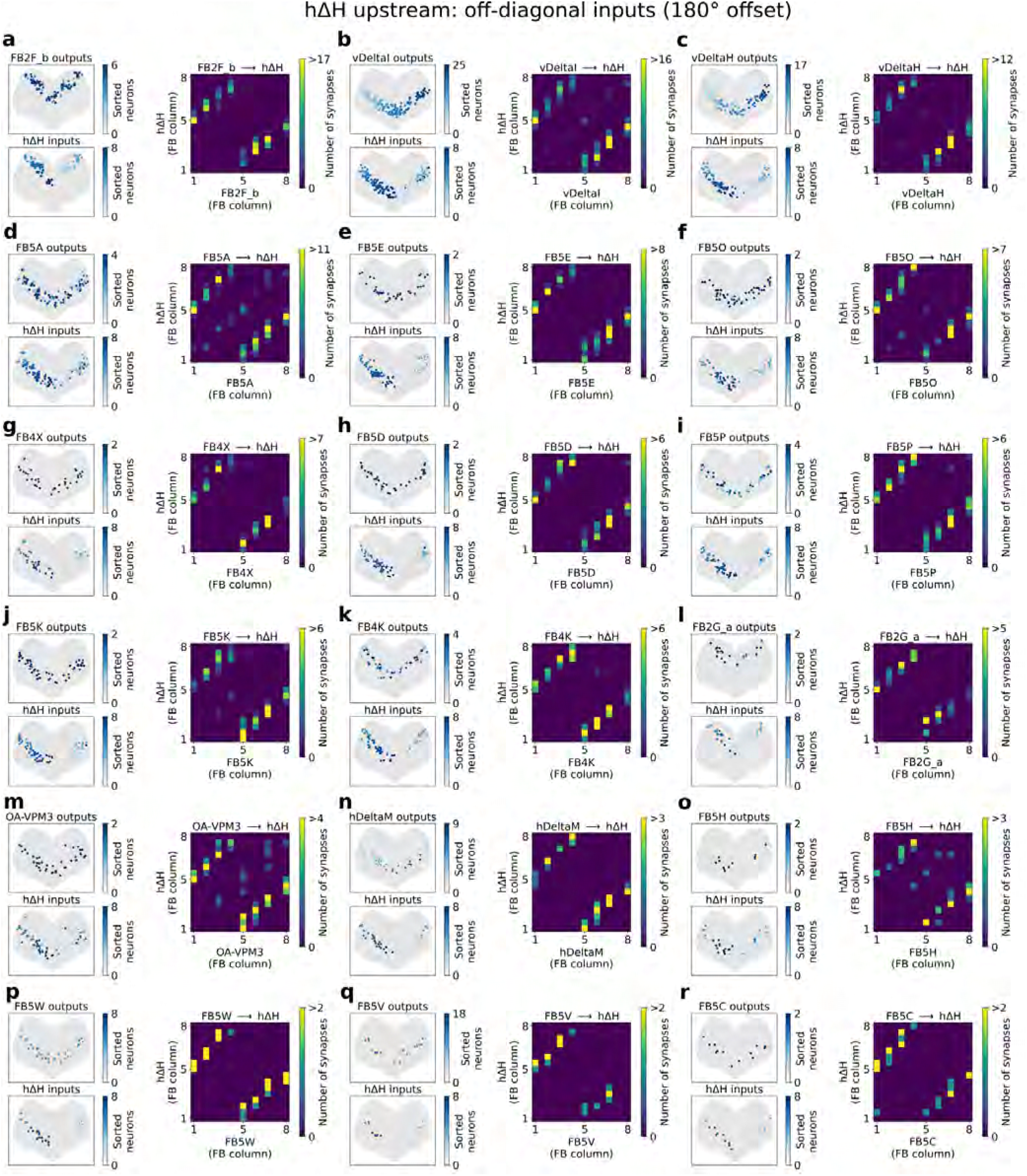
Same as Supplementary Fig. S35 but for hΔH.

**Figure S37.**
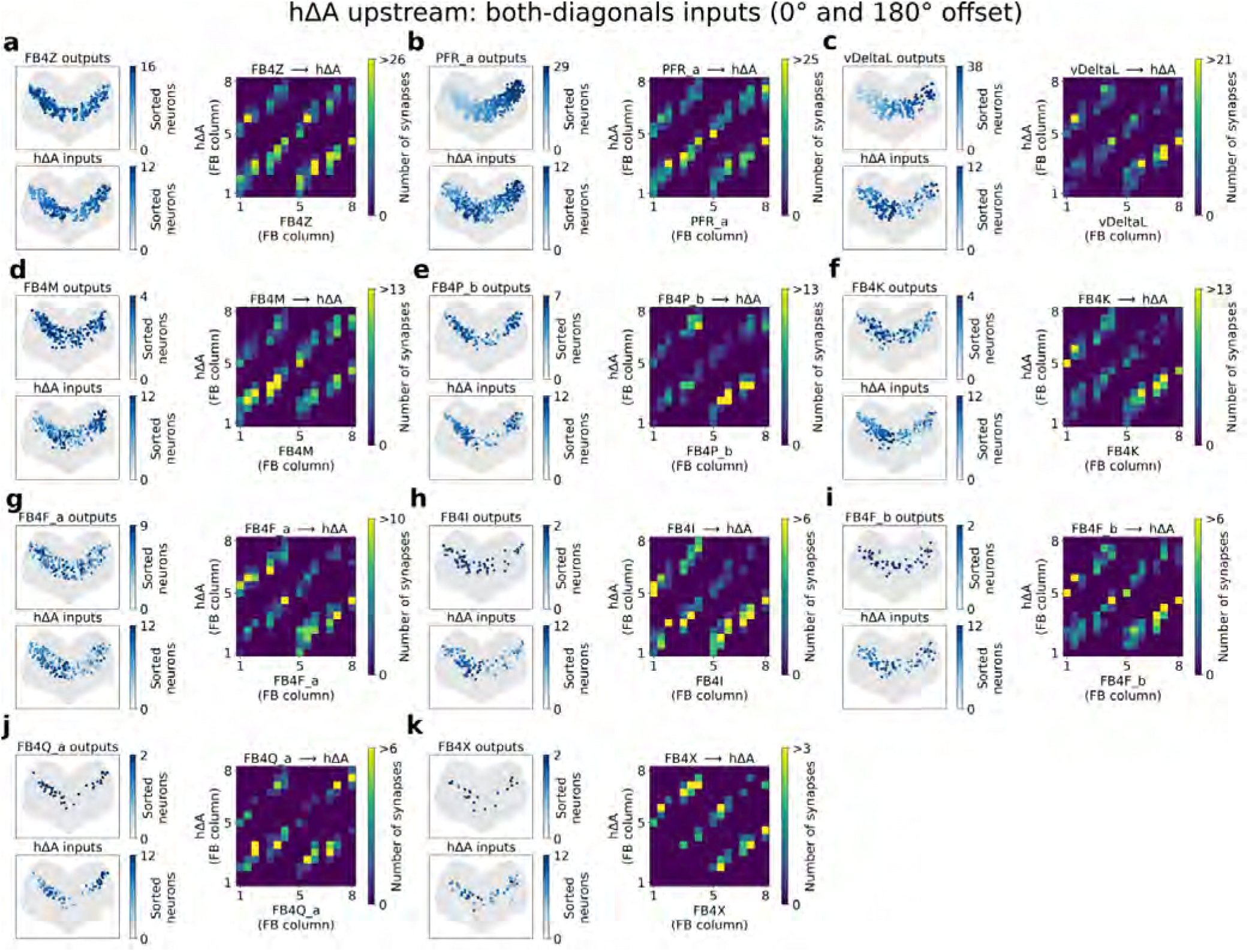
Same as Supplementary Fig. S33 but showing only upstream populations that target hΔA through both diagonals, including PFR neurons.

**Figure S38.**
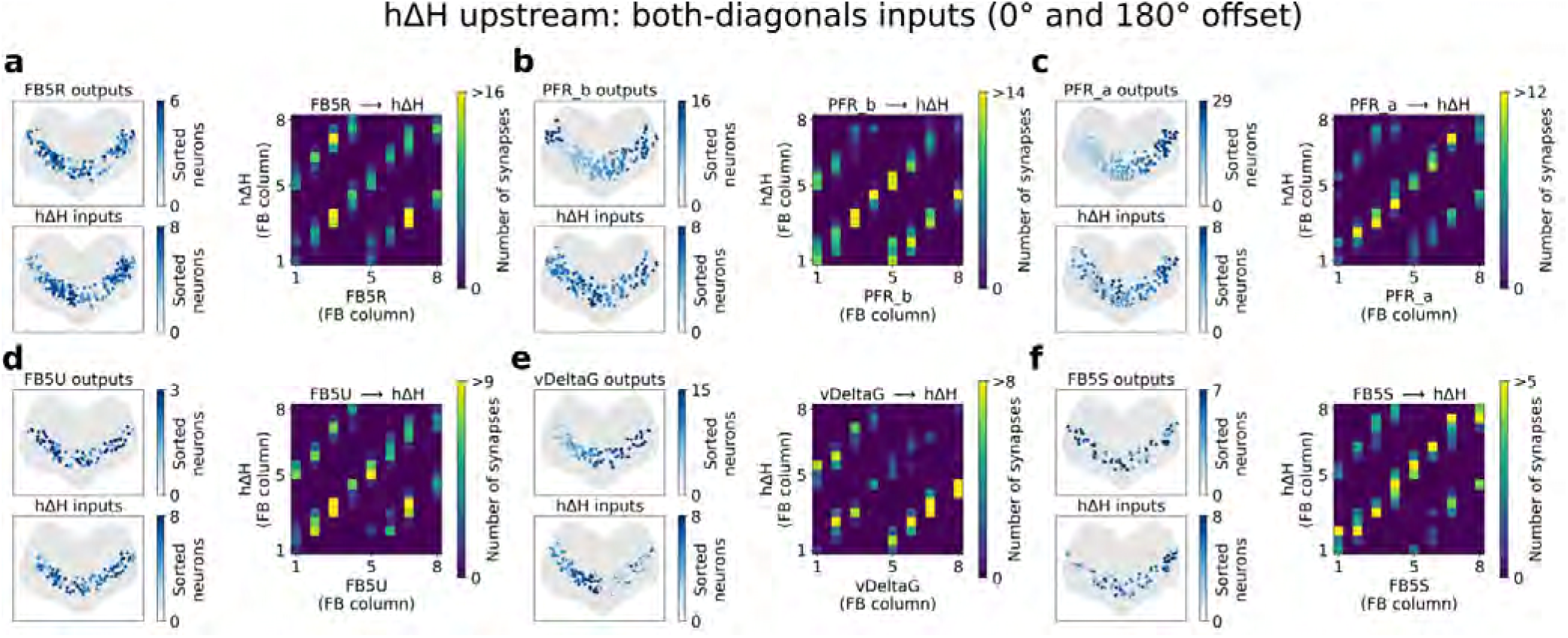
Same as Supplementary Fig. S34 but showing only upstream populations that target hΔH through both diagonals, including PFR neurons.

**Figure S39.**
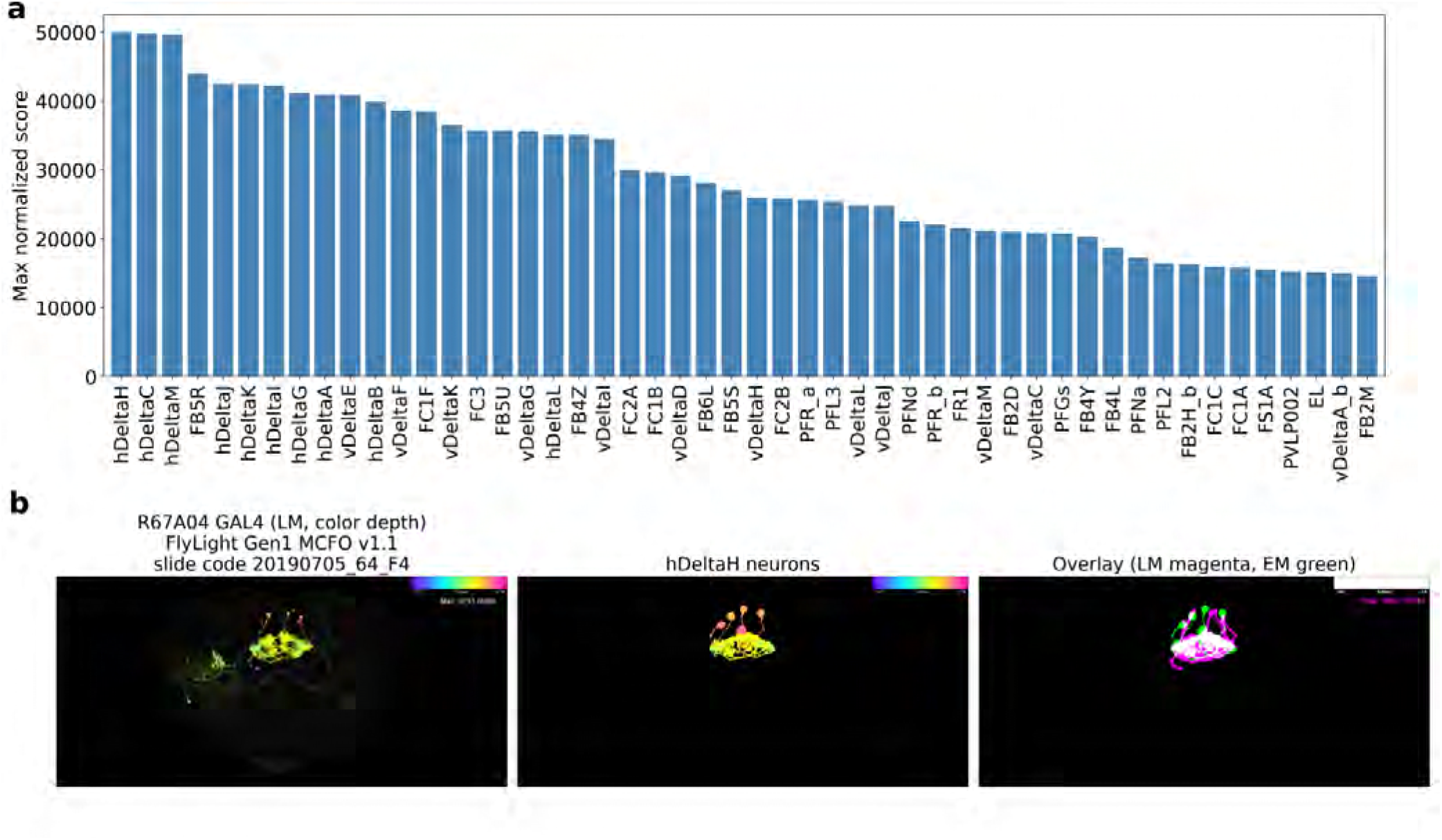
NeuronBridge matches between the GAL4 line R67A04 and EM neuron types. **(a)** Maximum normalized NeuronBridge color depth MIP search score for each EM neuron type across all R67A04 light microscopy (LM) images. Only EM neurons with inputs or outputs in the FB were included, and the 50 highest-scoring types are shown. **(b)**Example match for the R67A04 LM image with the highest hΔH score (FlyLight Gen1 MCFO v1.1, slide code 20190705_64_F4). Left: LM color depth MIP. Center: color depth MIPs of the best-matching hΔH EM neurons. Right: overlay of the LM signal (magenta) and the same EM neurons (green); overlap is shown in white. In the color depth MIPs, color indicates depth along the anterior–posterior axis.

**Figure S40.**
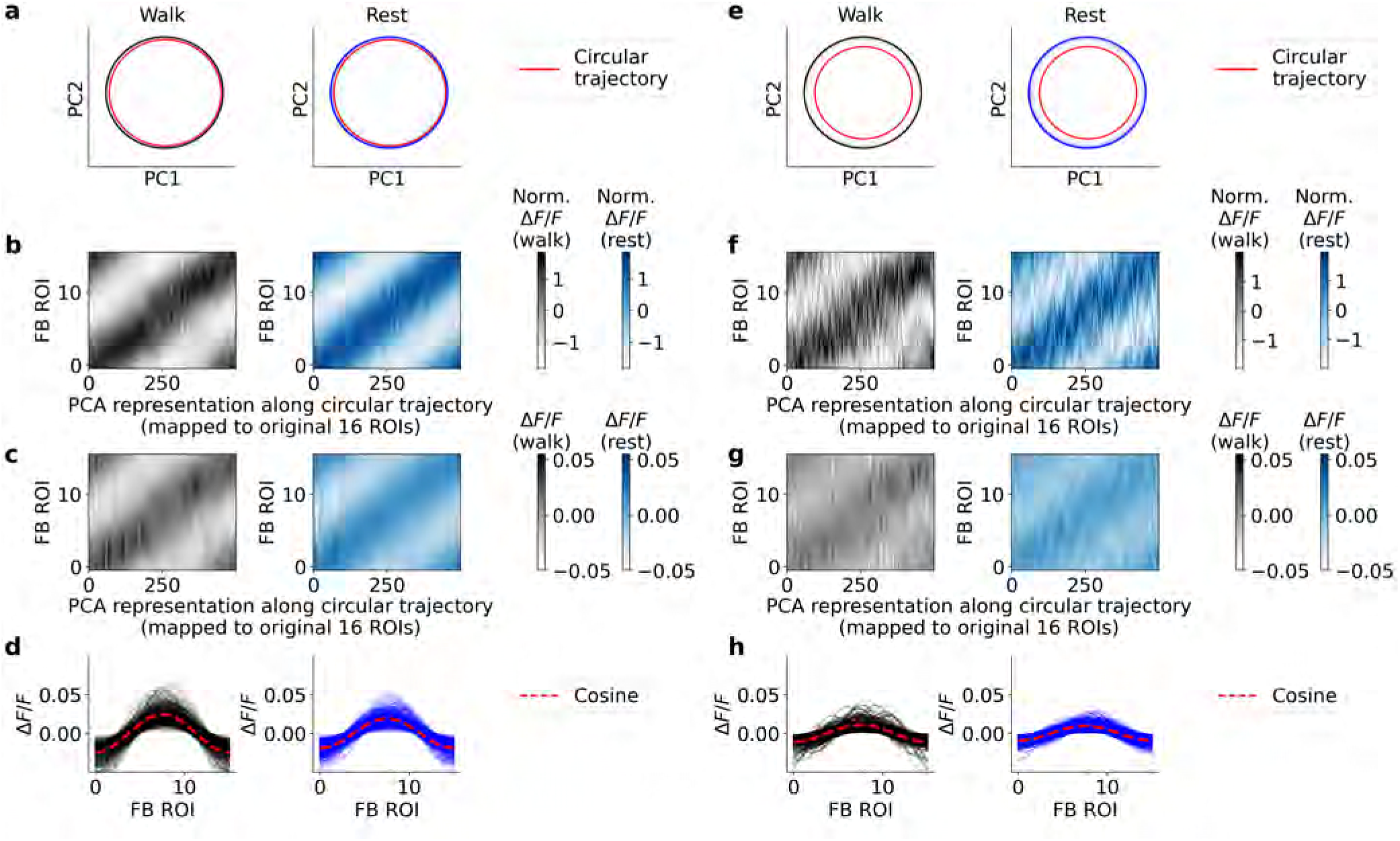
Same as Supplementary Fig. S5 but for *h*Δ neurons from a total of 12 flies, resulting in a total of 147,929 fluorescence traces.

**Figure S41.**
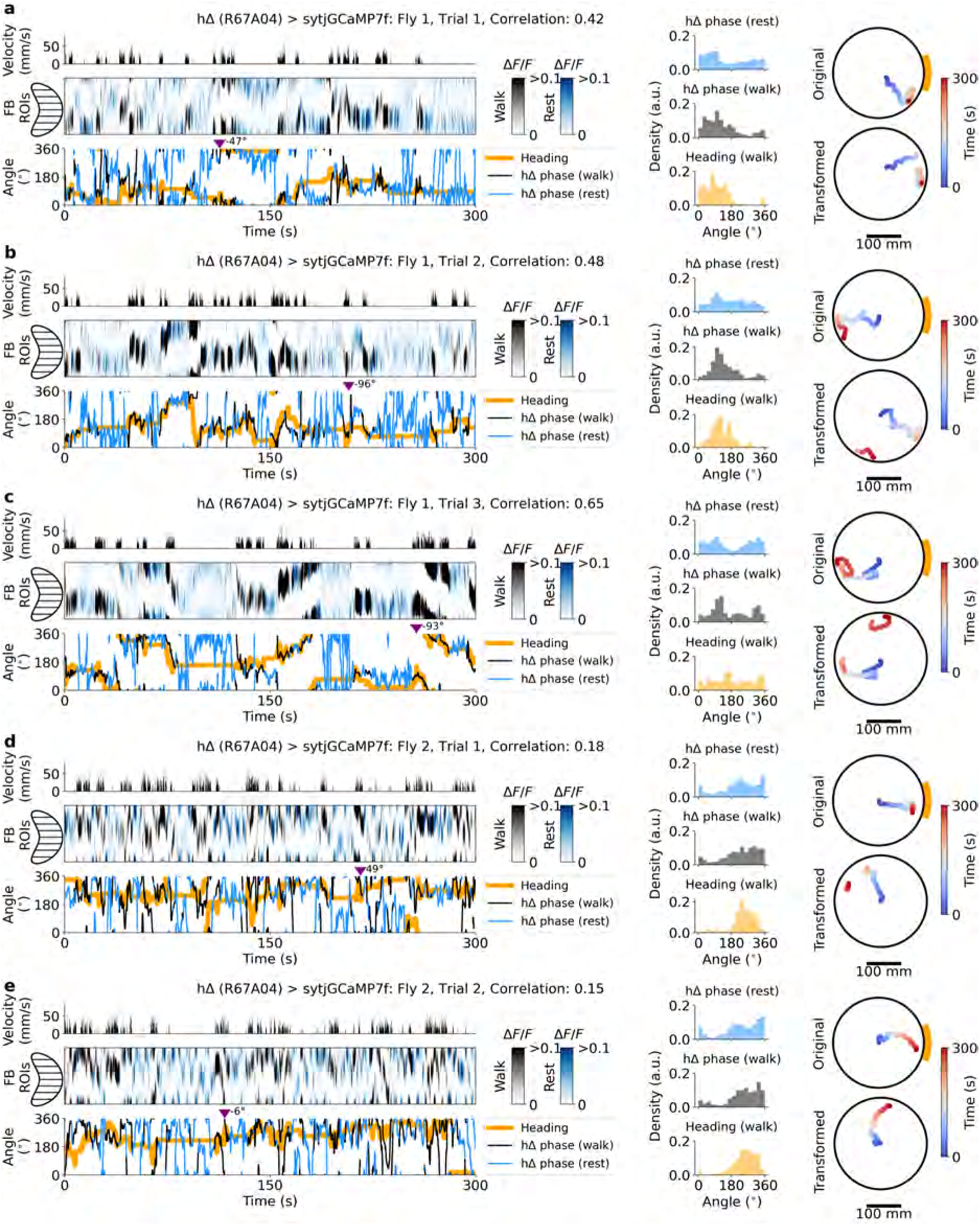
Neural activity and behavior in flies 1-2 expressing sytjGCaMP7f in hΔ neurons (GAL4 line R67A04) navigating the mechanical VR. Data shown are from selected trials (5 min duration) with *≥*60 s of walking and resting, and a PFR phase–heading correlation *>* 0.1 (selection criteria in Table S1). **a-e** Left panel: Fly velocity (top); fluorescence in FB ROIs during walking (black) and rest (green, middle); PFR phase during walking (black) and rest (blue), and fly heading (orange, bottom). Middle panel: PFR phase density during rest (blue, top) and walking (black, middle); heading density (orange, bottom). Right panel: Original fly trajectory in the mechanical VR (top); trajectory transformed by the two-segment offset model and rest-phase offset (bottom; see Methods and Extended Data Fig. 3).

**Figure S42.**
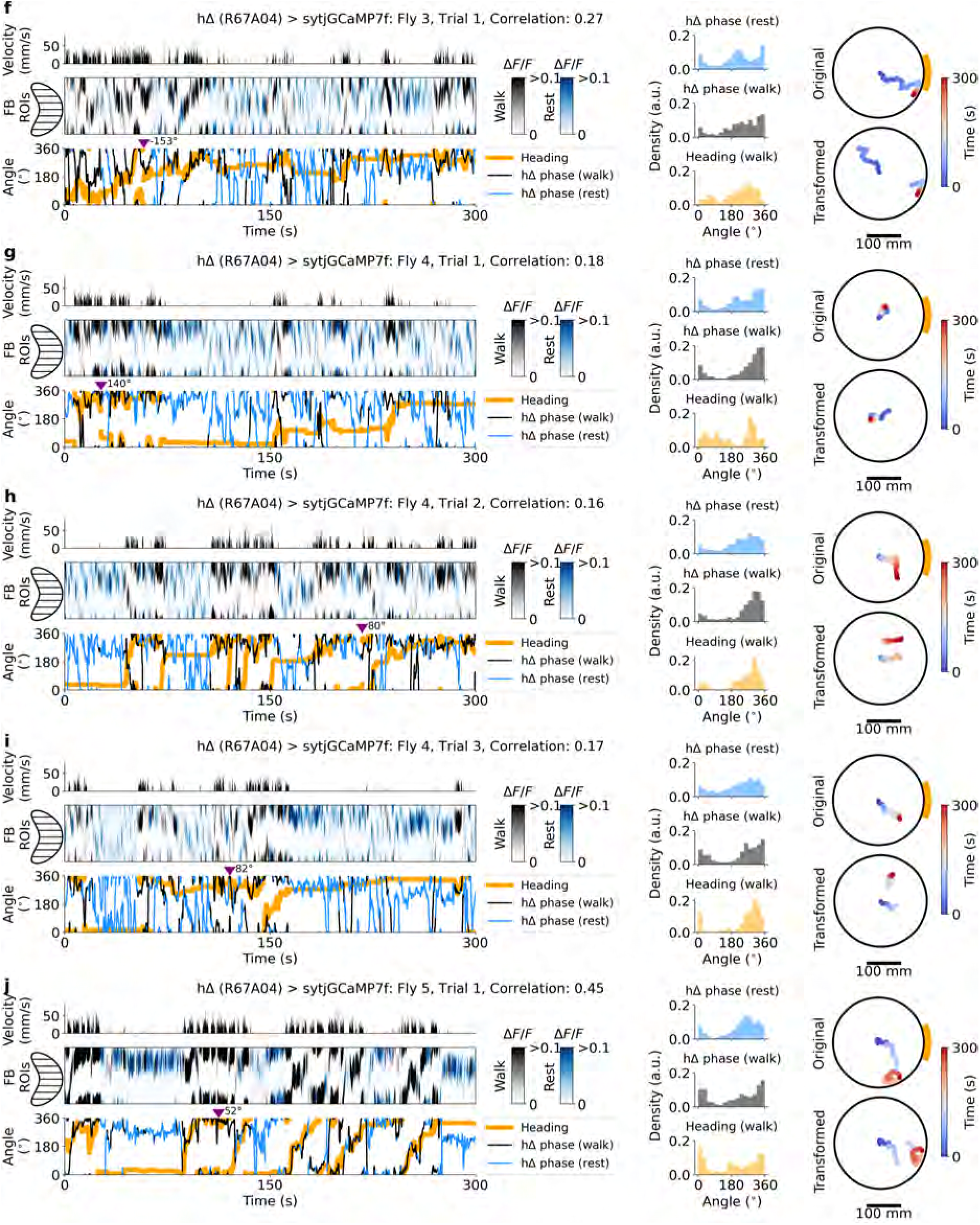
Neural activity and behavior in flies 3-5 expressing sytjGCaMP7f in hΔ neurons (GAL4 line R67A04) navigating the mechanical VR. **f-j** Same as Supplementary Fig. S41a-e.

**Figure S43.**
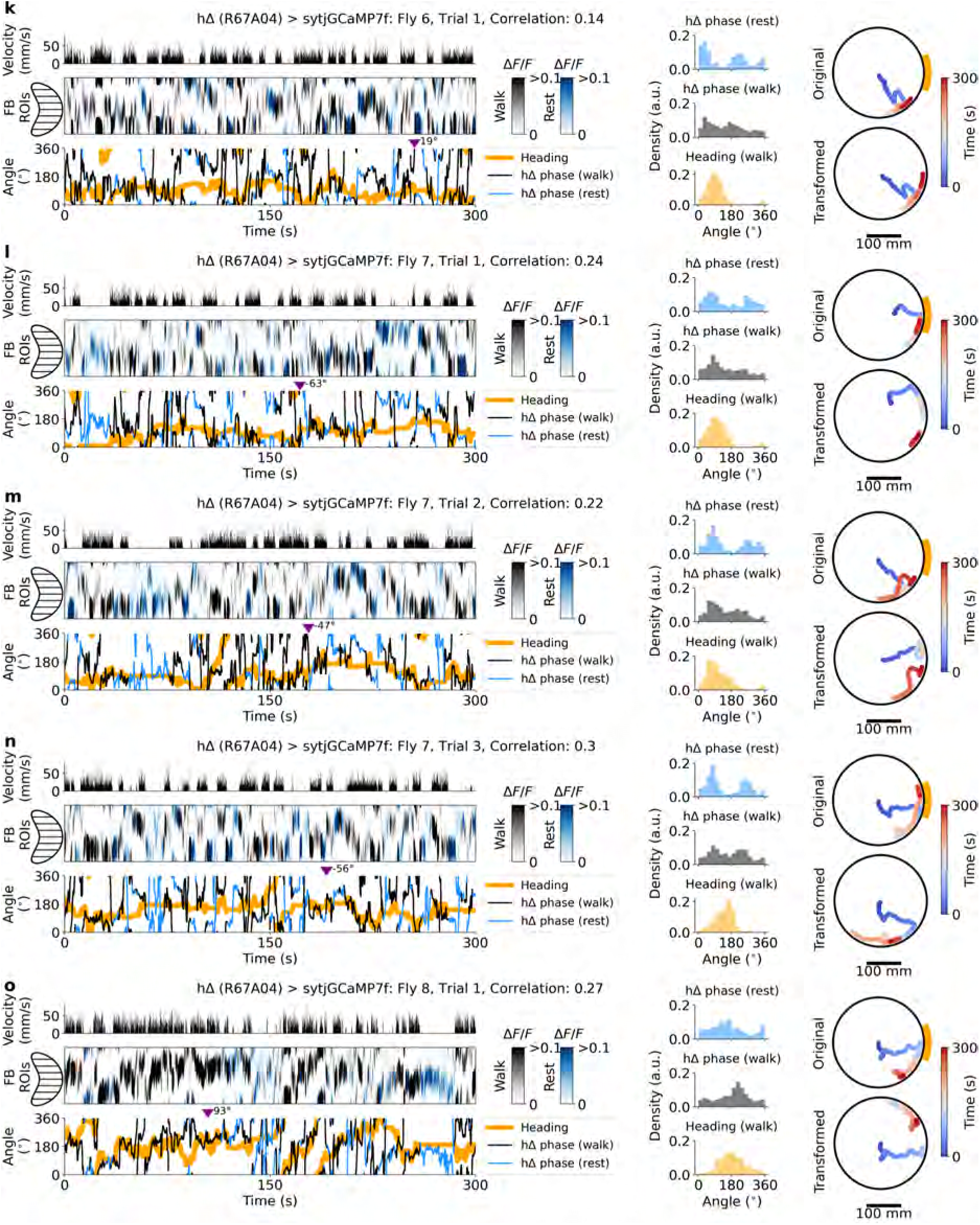
Neural activity and behavior in flies 6-8 expressing sytjGCaMP7f in hΔ neurons (GAL4 line R67A04) navigating the mechanical VR. **k-o** Same as Supplementary Fig. S41a-e.

**Figure S44.**
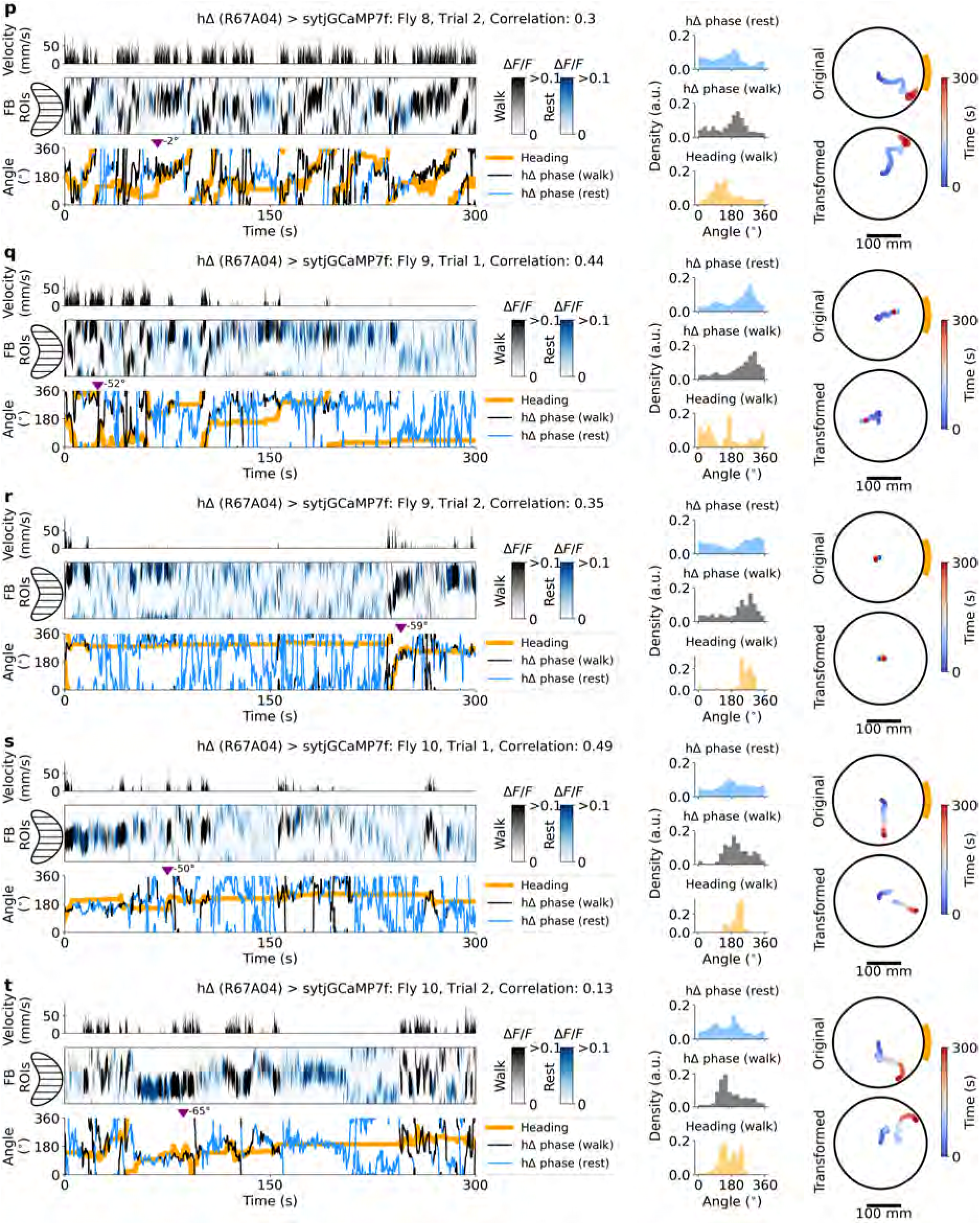
Neural activity and behavior in flies 8-10 expressing sytjGCaMP7f in hΔ neurons (GAL4 line R67A04) navigating the mechanical VR. **p-t** Same as Supplementary Fig. S41a-e.

**Figure S45.**
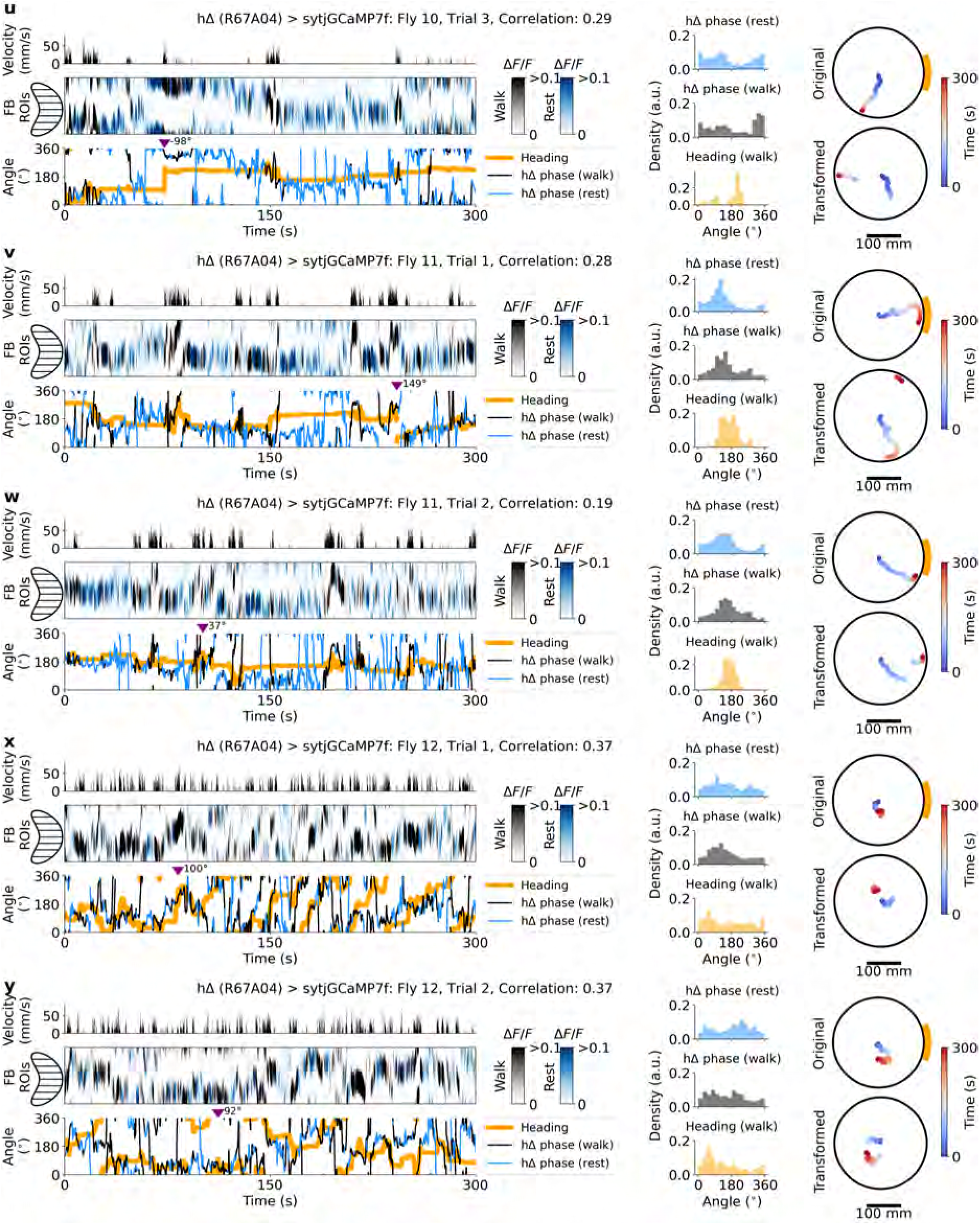
Neural activity and behavior in flies 10-12 expressing sytjGCaMP7f in hΔ neurons (GAL4 line R67A04) navigating the mechanical VR. **u-y** Same as Supplementary Fig. S41a-e.

**Figure S46.**
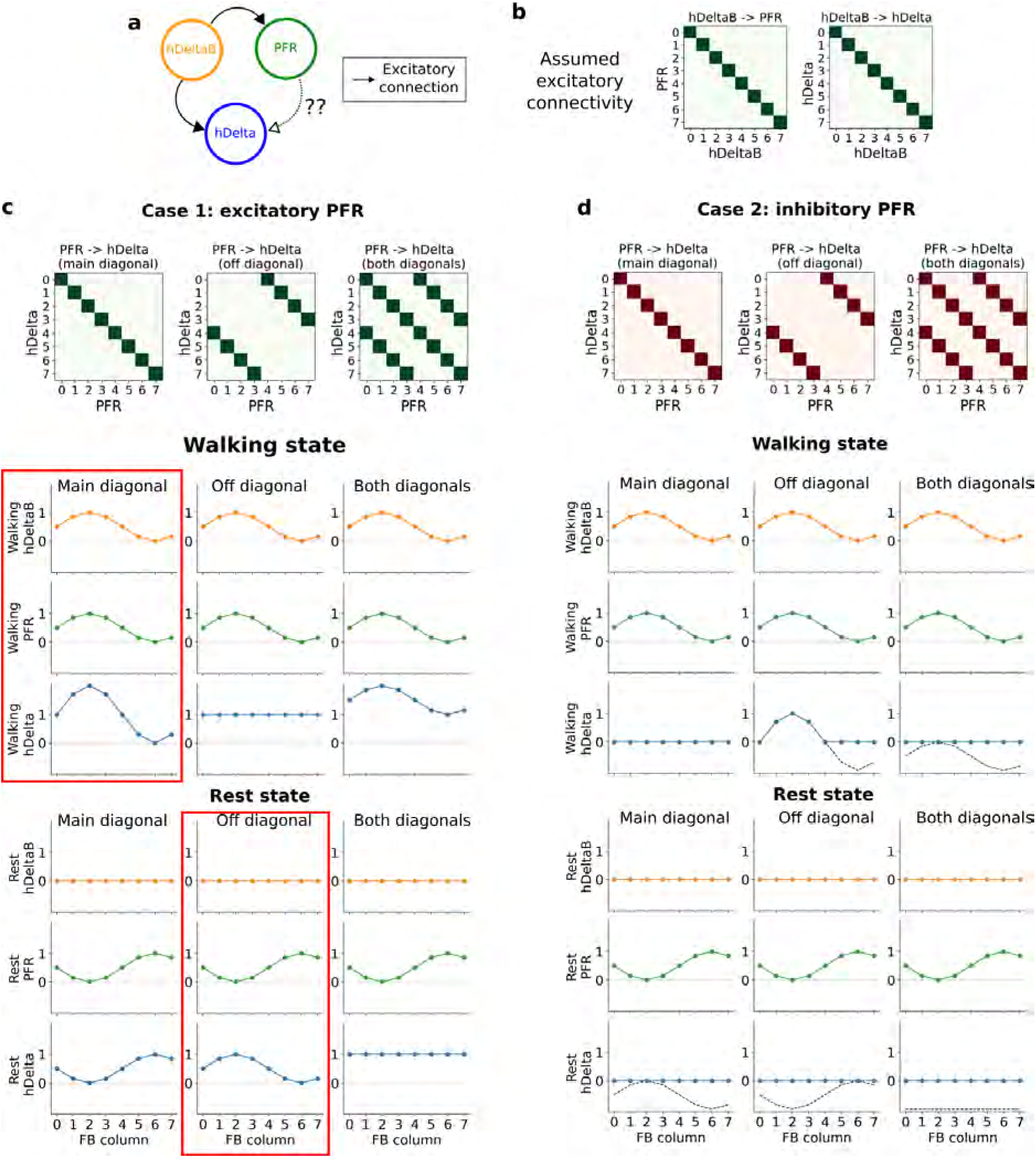
Anatomically constrained analysis of the sign and connectivity of PFR*→*hΔ synapses. **(a)** Three-population circuit including hΔB, PFR and hΔ neurons; the sign of the PFR*→*hΔ connection (dashed arrow) is unknown. **(b)** Assumed connectivity matrices (8 fan-shaped body columns) derived from the connectome^77^. **(c)** Case 1, excitatory PFR. Top: tested PFR*→*hΔ connectivity matrices (main diagonal, off-diagonal shifted by 4 columns, i.e. 180*^◦^*, or both). Middle and bottom: activity profiles during walking (hΔB: cosine peaking at column 2, transmitted to PFR via the main diagonal) and rest (hΔB: zero; PFR: cosine peaking at column 6, i.e. 180*^◦^* from the walking bump). Only the combination of main-diagonal transmission during walking and off-diagonal transmission during rest (red boxes) yields an hΔ bump at the same column in both states. **(d)** Case 2, inhibitory PFR: same layout as **c**. Dashed lines show hΔ activity before rectification. For all configurations, rectified hΔ activity at rest is flat or zero, so inhibitory PFR cannot reproduce the observed hΔ bump at rest without additional mechanisms (e.g. a constant excitatory input to hΔ combined with off-diagonal.)

